# Single-cell lineage tracing reveals hierarchy and mechanism of adipocyte precursor maturation

**DOI:** 10.1101/2023.06.01.543318

**Authors:** Guillermo C. Rivera-Gonzalez, Emily G. Butka, Carolynn E. Gonzalez, Wenjun Kong, Kunal Jindal, Samantha A. Morris

## Abstract

White adipose tissue is crucial in various physiological processes. In response to high caloric intake, adipose tissue may expand by generating new adipocytes. Adipocyte precursor cells (progenitors and preadipocytes) are essential for generating mature adipocytes, and single-cell RNA sequencing provides new means to identify these populations. Here, we characterized adipocyte precursor populations in the skin, an adipose depot with rapid and robust generation of mature adipocytes. We identified a new population of immature preadipocytes, revealed a biased differentiation potential of progenitor cells, and identified Sox9 as a critical factor in driving progenitors toward adipose commitment, the first known mechanism of progenitor differentiation. These findings shed light on the specific dynamics and molecular mechanisms underlying rapid adipogenesis in the skin.

## Main Text

White adipose tissue is central to the regulation of energy storage and metabolism, but it can also be involved in the immune response, wound healing, fibrosis and hair growth *(1–4)*. Mature adipocytes, the main cell type in adipose tissue, represent a highly specialized population that stores excess energy in the form of lipids. To accommodate an energy surplus, adipose tissue can expand through two main mechanisms: hypertrophy and hyperplasia. Hypertrophy refers to the increase in cell size in pre-existing mature adipocytes, while hyperplasia indicates the creation of new mature adipocytes which is a critical mechanism to maintain a healthy adipose tissue *(5, 6)*. Since mature adipocytes are post-mitotic under most conditions *(7, 8)*, the generation of new mature adipocytes relies on the proliferation and differentiation of adipocyte precursor cells (APCs) which encompass progenitor and preadipocyte cells. Several surface markers have traditionally been used to identify APCs in different murine white adipose depots *(9–12)*. Recently, single-cell RNA sequencing (scRNA-seq) of these depots has provided a more unbiased approach for identifying and classifying APC populations *(13–18)*. For example, preadipocyte cells committed to the generation of new mature adipocytes can be identified by *Icam1* gene expression. In contrast, progenitor cells are labeled by the expression of *Dpp4*. Transplant assays have suggested a coarse hierarchical relationship between progenitors and preadipocytes; according to this model, progenitors give rise to cells with a preadipocyte-like phenotype *(14)*. This model assumes synchronous replacement as differentiation progresses. However, adipose depots exist in different anatomical locations and have distinct biological properties *(19)*. For example, in response to a high-fat diet, it can take up to 8 weeks for the subcutaneous and visceral depots to produce new mature adipocytes *(20, 21)*. In contrast, it only takes the dermal depot 11 days on average to produce new mature adipocytes under homeostatic conditions, a process tightly coordinated with the hair follicle cycle *(22, 23)*. Studying these different microenvironments, developmental stages, and other biological conditions (such as high-fat diet inflammation and cell cross-talk) could uncover new phenotypes of adipocyte precursor cells, hierarchical relationships, process of adipogenesis, and mechanisms that regulate adipogenic differentiation. Ultimately, identifying the properties of adipose precursors in a depot-specific manner could lead to therapeutic interventions that increase mature adipocyte production to achieve positive health benefits.

Understanding adipogenic differentiation requires the use of high-resolution lineage tracing. Current approaches to lineage-trace APCs either lack specificity or restrict the tracing to a small subset of cells and make competition assays technically challenging, which limits the interpretation of the results. Therefore, the use of cell barcoding is a very attractive option to characterize, with unparalleled resolution, the hierarchy of cell populations. Here, we identified and validated *Dpp4* and *Cd9* as markers for progenitor and preadipocyte populations in the skin using cell-barcoding and transplantation approaches. We profile a previously uncharacterized population of immature preadipocytes (*Egfl6*^+^) that occupies a developmental niche along this differentiation trajectory and represents a stable cell state in the skin. Using our unbiased genetic barcoding method, ‘CellTagging,’ *(24–26)* we label and track progenitor and preadipocyte cells isolated from the dermal depot to measure their unique differentiation potential during homeostatic adipogenesis. We identify progenitor cells as an important source of an uncharacterized population of immature preadipocytes and describe how progenitor cells are more likely to acquire markers of commitment to the adipose lineage, such as *Fabp4*. In contrast, pre-existing preadipocytes acquire other transcriptional characteristics and do not follow a linear path, providing the first evidence of divergent differentiation potential between progenitors and preadipocytes. Moreover, we identify *Sox9* as a key factor that preferentially drives progenitors toward both immature and committed preadipocyte stages. Overall, our results provide evidence for uncharacterized preadipocyte phenotypes, detailed hierarchical dynamics between progenitors and preadipocytes and establish the first known molecular mechanism of differentiation in progenitor cells.

## Results

### Identification of an immature preadipocyte population in the skin

We performed scRNA-seq on Lin-and CD29+/CD34+ double-positive cells isolated from the skin of postnatal day 21 (P21) mice, which contain a mixture of APCs, to identify putative populations of progenitors and preadipocytes *(16)* (Fig. 1A). After performing standard quality control and filtering, we obtained 14,478 APCs across 12 transcriptionally different clusters (Fig. 1B). Our data recapitulated previous findings of two major populations: putative heterogenous progenitors (clusters 0, 1, 2, 4, 5, 6, 9, and 10), and preadipocytes (clusters 3, 7, 8, and 11), based on selected markers from inguinal APCs. (Fig. 1C). Surprisingly, cluster 7 exhibited a small proportion of APCs expressing genes such as *Fabp4* and *Pparg*, which are characteristic of committed preadipocytes *(27)* (fig. S1A). We also observed high expression of *F3* (*Cd142*) in cells across multiple clusters (fig. S1B), in contrast to the restriction of *F3* expression to a single cluster, as previously reported in inguinal adipose. Conflicting reports label F3-expressing cells as either adipogenic, or a subset of non-adipogenic cells *(13, 14)*.

**Fig. 1.**
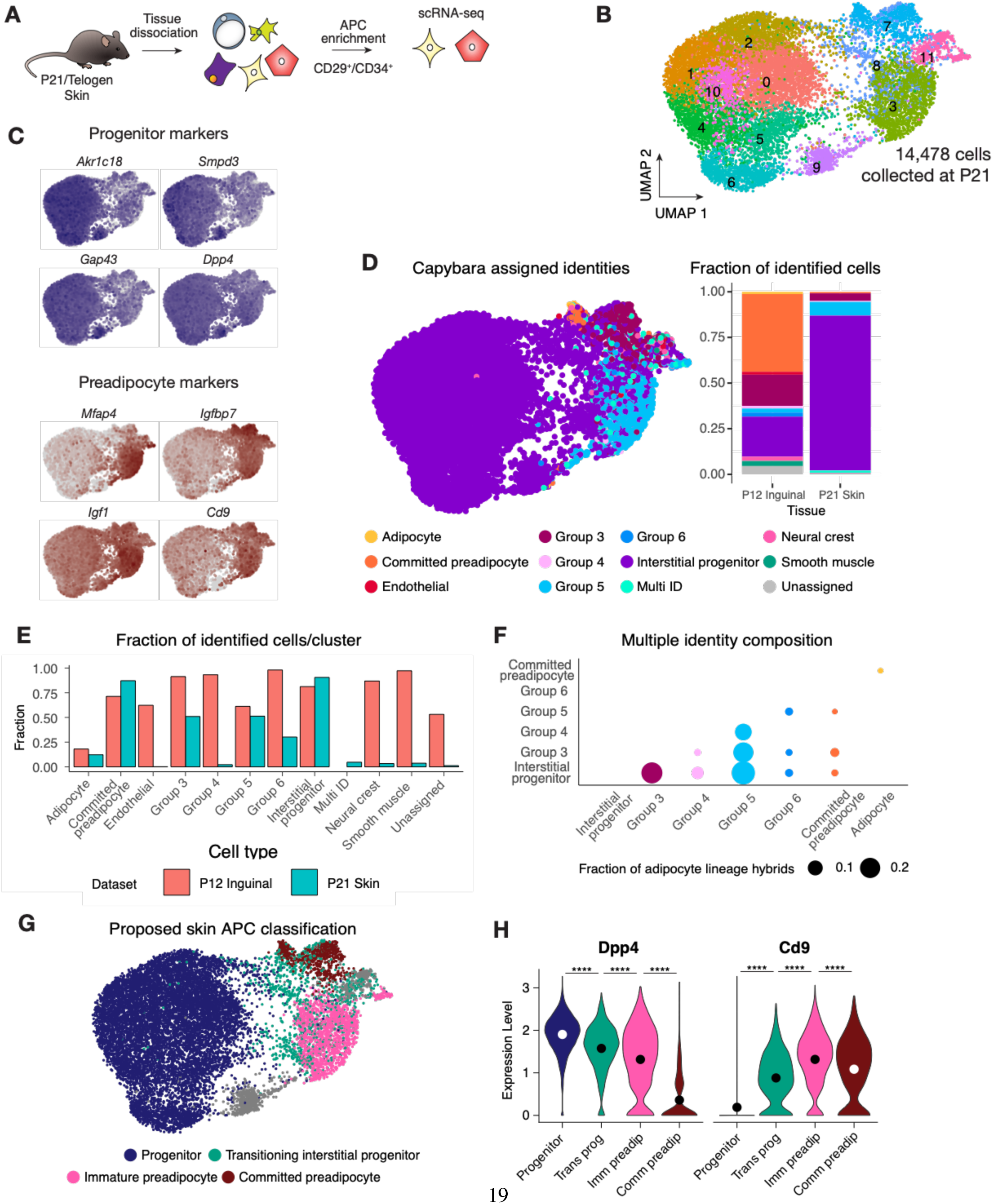
Identification of a putative immature preadipocyte population in the skin. (A) Graphic description of APC isolation and enrichment for scRNA-seq experiment. (**B)** UMAP of sequenced APCs isolated from mouse skin (14,478 cells pooled from 2 male and 2 female C57BL/6J mice). **(C)** Feature expression of progenitor and preadipocyte markers of skin APCs. **(D)** Capybara classification of skin APCs using the inguinal dataset by Merrick et al. *(14)* as the reference (left) and breakdown of relative cell type abundance (fraction of total cells) per dataset, inguinal (reference) or skin (right). **(E)** Fraction of Capybara identified cells per cluster in inguinal and skin datasets. **(F)** Composition and the fraction of skin APCs identified as containing multiple identities (Multi ID) in the adipocyte lineage. **(G)** Proposed classification of dermal APCs. **(H)** *Dpp4* and *Cd9* expression gradients for proposed classification of dermal APCs; dots indicate mean expression values (unpaired t test with Welch’s correction, two-tailed, ****p < 0.0001).

We utilized an unbiased approach to characterize our single-cell data in the context of known populations of APCs, in light of our identification of multiple F3^high^ populations. This approach prevents misidentification of cell types resulting from the use of subjective markers and enables us to compare transcriptional identities between skin and inguinal depots. Subsequently, Capybara, a computational single-cell classifier, assigned cell identities relative to a known reference using whole transcriptomic information, eliminating the need for manual marker selection *(28)*. To construct the reference, we manually annotated the existing inguinal atlas according to the same markers used by Merrick *et. al. (14)* (fig. S1C-D) and used Capybara to label skin APCs based on this inguinal classification (Fig. 1D). By applying Capybara, we confidently identified the cells originally considered putative progenitor cells as transcriptionally similar to interstitial progenitor cells in the reference (fig. S1E). Although the match between skin preadipocytes and inguinal cell types lacked statistical significance, Capybara suggested the best fitting identities for several of our distinct skin preadipocyte populations (Fig. 1D). Most frequently, the strongest matches for skin preadipocytes (cells in clusters 3, 7, 8, and 11) were the committed preadipocyte, group 3 and group 5 cell identities. Our preadipocyte cluster 7 aligns most closely with inguinal group 3 (*F3*) and committed preadipocytes. Notably, this cluster does not exhibit the highest *F3* expression in our dataset (fig. S1B), like group 3 cells do in inguinal tissue. Surprisingly, skin preadipocyte cluster 3, the population with the highest *F3* expression (fig. S1B), matches best with previously uncharacterized inguinal group 5. Among the most interesting differences between datasets is the strong enrichment for interstitial progenitors in the skin (Fig. 1D, right). Also, while inguinal group 5 cells were not previously suggested to have any adipogenic potential, our data is significantly enriched for this population of cells compared to the inguinal dataset. Subsetting and reclustering of preadipocyte cells did not provide better separation among different Capybara classifications, suggesting that in contrast to progenitors, preadipocytes do not display identical phenotypes across depots (fig. S1F).

Taking an orthogonal approach to compare cell identities across datasets, we integrated skin and inguinal datasets *(14)*. Briefly, we used anchors to co-embed the two datasets, and then performed Louvain clustering on the super-embedding (fig. S1G). We observed varying degrees of co-clustering for each cell type, with generally higher fractions of inguinal cells co-clustering with cells of the same cell type (fig. S1H). Interstitial progenitors, committed preadipocytes, group 3 and group 5 cells have the highest co-clustering, which is in line with Capybara classification (Fig. 1E). Taken altogether, skin interstitial progenitors show a strong identity match to inguinal progenitors. However, the preadipocyte populations found in the skin exhibit clear differences not only in identity but also in proportion from those found in the inguinal depot. Most of our putative preadipocyte population consists of skin preadipocyte cells resembling inguinal group 5 cells, while the number of cells identified as committed preadipocytes and group 3 cells remain low. Interestingly, Capybara identified 203 multi-ID cells (cells displaying multiple identities that meet predetermined confidence thresholds). These cells co-clustered with various types of preadipocytes (fig. S1I). Most of the multi-ID cells (154; 76%) shared interstitial progenitor, group 3 (*F3+*), group 4, and group 5 identities (Fig. 1F). From this we hypothesize that these rare multi-ID cells may occupy adipogenic transition states between these APC identities, suggesting that group 5 cells could be part of the adipogenic lineage. The observed differences in identity and relative abundances highlight the likely presence of depot-specific preadipocyte phenotypes in the skin.

These findings led us to generate a new classification paradigm specific to the skin (Fig. 1G, fig. S1J; table S1). Cells from clusters 0, 1, 2, 4, 5, 6 and 10 were assigned a progenitor identity. Cells identified as progenitors by Capybara that showed increasing levels of Cd9 expression, occupying clusters 3 and 8, were labeled as transitioning progenitors. Non-progenitors in clusters 3 and 8 were labeled as immature preadipocytes given their preadipocyte phenotype and lack of *Pparg* expression. Interestingly, 99% of cells identified as Group 5 are labeled as immature preadipocytes. Lastly, committed preadipocytes and group 3 cells were labeled as committed preadipocytes since they cluster together in our dataset (cluster 7) and further subclustering failed to segregate them (fig. S1F). Cells in clusters 9 and 11 were omitted from this reference construction due to the heterogeneity of these clusters.

In order to understand whether our putative skin APC populations are capable of producing mature adipocytes and taking advantage of the *Dpp4* and *Cd9* expression gradient which correlated with putative differentiation stages (Fig. 1H), we devised a FACS isolation strategy to enrich for progenitors (DPP4^high^), immature preadipocytes (CD9^high^/F3^high^), and committed preadipocytes (CD9^high^/F3^low^; fig. S2). We identified a population of DPP4+/CD9^low^ cells and a continuum of CD9+ cells with variable DPP4 expression (fig. S3A), similar to our scRNA-seq data (Fig. 1C). As observed in our scRNA-seq dataset, preadipocytes can be further separated based on cell surface F3 protein expression, with putative immature preadipocytes displaying high F3 expression and committed preadipocytes displaying low F3 expression (fig. S3A). We validated that populations isolated by FACS matched selected differentially expressed genes identified during scRNA-seq analysis by qPCR (fig. S3B). Through transplant and *in vitro* differentiation assays, we confirmed the adipogenic potential of isolated progenitors, immature preadipocytes, and committed preadipocytes (fig. S3C-E). These results confirm the adipogenicity of isolated progenitors and committed preadipocytes and establishes for the first time the adipogenic potential of a novel population of immature preadipocytes, marked by high levels of *Cd9* and *F3* expression.

### Lineage tracing highlights complex APC hierarchy and differentiation potential

While there is evidence that committed preadipocytes and group 3 cells originate from progenitors *(14)*, the origin of immature preadipocyte cells is unknown. Given our understanding of the adipogenic potential of adipocyte progenitors to generate multiple populations of preadipocytes, we hypothesized that progenitors produce this immature preadipocyte population. We performed single-nuclei RNA sequencing (snRNA-seq) of whole skin to identify potential markers that exclusively label adipocyte progenitors. Cell types were annotated using known gene expression markers (fig. S4A and table S2). We were unable to identify a single marker to specifically lineage trace progenitors due to the gene expression overlap between multiple populations in this tissue. For example, the selection of *Pdgfra*-Cre mouse models *(23, 27, 29)* would not be sufficient to distinguish between adipocyte progenitors and preadipocytes since both populations showed high *Pdgfra* expression (fig. S4B-D). The same conclusion is reached when considering the use of the *Dpp4*-Cre model *(30)*, since several preadipocyte clusters display lingering *Dpp4* expression, an observation we also confirm by FACS analysis (fig. S4B-D and fig. S3A).

Given the lack of precise tools to label our populations of interest *in vivo*, we decided to use the genetic barcoding approach designed by our group, CellTagging *(25, 31)*. CellTagging labels cells via lentiviral vectors encoding a *GFP* gene with a genetic barcode in the 3’ UTR. scRNA-seq detects CellTags, capturing transcriptional and lineage information in the same assay, from the same cells. To identify the origin of immature preadipocytes, we CellTagged both progenitors and a mixed population of preadipocytes with two non-overlapping CellTag libraries. This allowed us to discriminate between progenitors and preadipocytes and perform competitive transplant assays (Fig. 2A). We introduced a dominant-negative form of CEBP-A (DN-CEBPA) before cell injection to prevent the loss of transplanted cells due to mature adipocyte formation *(32)* This prevented the lipid filling and loss of APCs with little expected side effects on APC biology given the lack of biological effects on cultured preadipocytes before hormonal induction *(33)* and since *Cebpa* expression in our cells is turned on in committed preadipocytes (fig. S5A). Briefly, cells were isolated from P21 mouse skin, transduced with identifying barcoding libraries and DN-CEBPA virus through viral transduction (the v2 library was used for adipocyte progenitors, and has a unique set of barcodes completely different from the v1 library which was used for preadipocytes, allowing competition assays to be performed). Then cells were mixed in equal numbers and transplanted into the skin of P21 mice during the telogen stage (Fig. 2A). At around P21, the hair follicle cycle starts transitioning from the telogen stage to the anagen stage, exhibiting a natural uptick of adipogenesis in the skin. Taking advantage of this process allowed us to inject labeled cells into an environment primed for promoting adipogenic potential. Between 11 and 13 days after transplantation, cells were recovered from the skin during the anagen stage when hair growth-induced adipogenesis occurs *(4)*, and transplant (GFP+) and host cells (GFP-) were isolated by FACS and processed together for scRNA-seq (Fig. 2B).

**Fig. 2.**
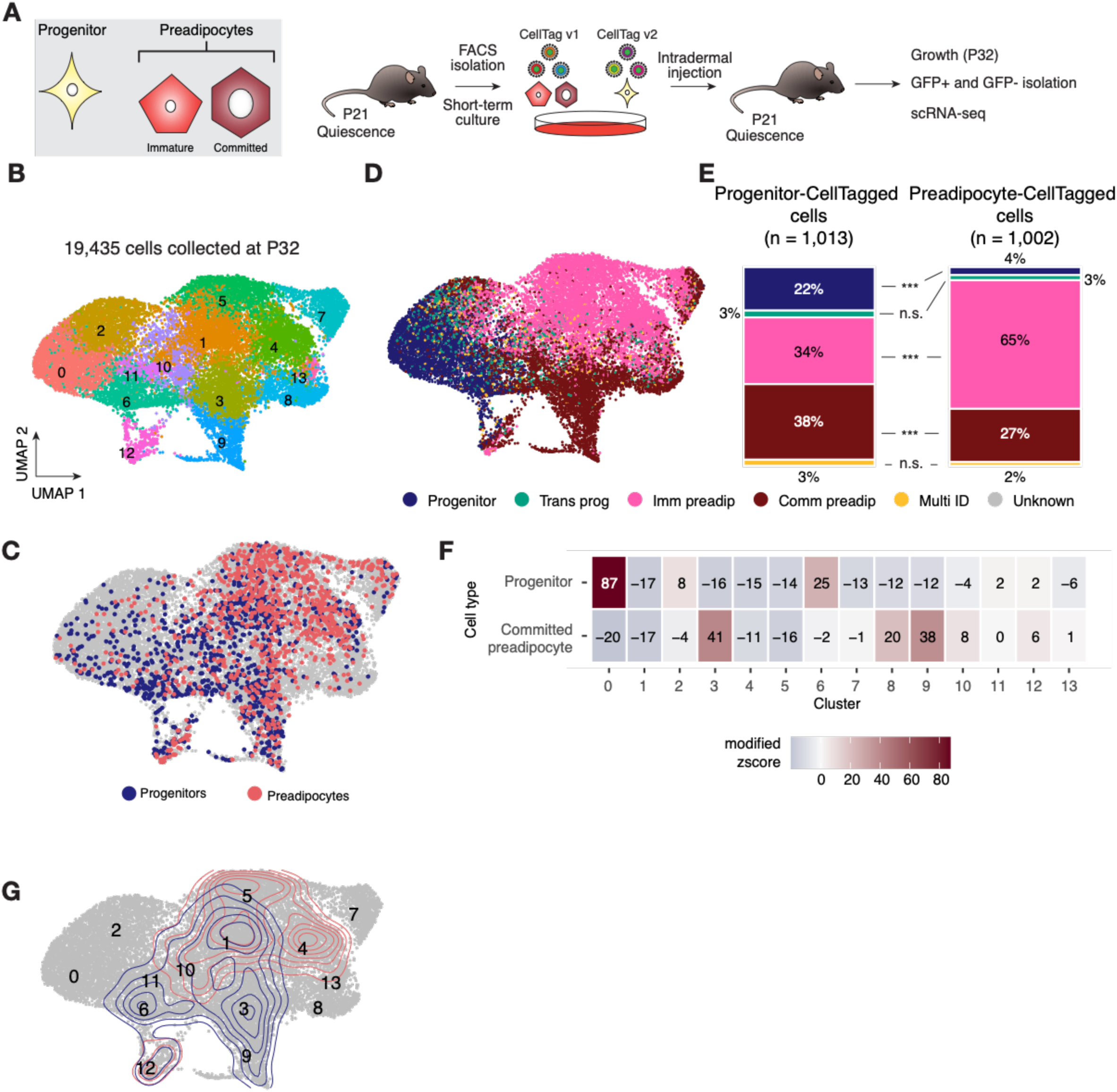
Lineage tracing of progenitors and preadipocytes highlights complex APC hierarchy and differentiation potential. (A) Graphical representation of *in vivo* lineage-tracing competition assay for skin progenitors and preadipocytes. Both immature and committed preadipocytes were tagged as a single population (See cell key on the left). **(B)** UMAP representation of 19,435 sequenced APCs from 4 different biological replicates (n = 4 male C57BL/6J mice) at age P32 (P32 APC dataset) after identification, QC analysis, and integration. **(C)** Identification by CellTag barcode of transplanted progenitors and preadipocytes in the UMAP space of the P32 APC dataset. **(D)** Capybara classification of age P32 APCs using our previous P21 dataset as a reference. **(E)** Quantification of Capybara cell identities of CellTagged progenitors and CellTagged preadipocytes (***p < 0.0001, randomization test, two-tailed). **(F)** Modified z-scores of Jaccard similarities for each cluster in the P32 APC dataset compared to the progenitor or committed preadipocyte identities (1,000 Jaccard similarities were calculated using randomized data for background distributions). **(G)** Contour plots identifying Louvain clusters significantly enriched by permutation testing for transplanted progenitors, preadipocytes or both in the P32 APC dataset; colors correspond to populations in **(F)** (analysis was performed with 1,000 permutations).

APCs were identified based on mesenchymal-associated genes as well as *Cd34* and *Itgb1* co-expression which identify adipocyte precursors *(23, 34)* (table S3). After quality control, we analyzed 19,435 APCs from four independent biological replicates, of which 10,590 (54.4%) were host cells and 8,845 (45.5%) were transplanted cells, identified by GFP and CellTag expression (Fig. 2C and fig. S5B-D). Capybara classified adipocyte progenitors, transitioning progenitors, immature preadipocytes, committed preadipocytes, and multi-ID cells (Fig. 2D) using our reference P21 skin dataset (Fig. 1E). Differential gene expression analysis between host and transplanted progenitors suggests the transplantation process induced only minor changes in gene expression (table S4). 78 genes were significantly differentially-expressed between the two progenitor populations, representing 0.4% of the total 19,594 unique features captured in the dataset. Identifying the origin (adipocyte progenitor or preadipocyte) uncovered the differentiation potential of transplanted cells. Cells of progenitor origin (identified by unique barcodes) differentiate into committed preadipocytes (38%), and immature preadipocytes (34%), while a smaller proportion retained a progenitor signature (22%) (Fig. 2E left panel, fig. S5E-G). Transplanted preadipocytes (identified through unique barcode expression) remained as or generated a high proportion of immature preadipocytes (65%) and a lesser number of committed preadipocytes (27%) (Fig. 2E, right panel, table S5). These findings suggest that progenitor cells generate immature preadipocytes and are likely an important source for these cells.

To assess differentiation potential of our distinct populations, we used Jaccard indices to assign each Louvain cluster a relative similarity to the progenitor or committed preadipocyte identity (Fig. 2F). These identities mark previously known and widely-accepted early and late points of the adipogenesis differentiation trajectory in the skin and other depots *(14, 27)*. Ranking Jaccard indices allowed us to identify clusters with high committed preadipocyte and low progenitor identity (clusters 3 and 9). Permutation tests identified that clusters 3 and 9 (mostly committed preadipocytes) and 6 (progenitors) were only enriched for cells of progenitor origin (p = 0.00015, p < 0.0001, p < 0.0001, respectively, permutation test, one-sided). Clusters 4 and 5 (immature preadipocytes) were only significantly enriched for cells of preadipocyte origin (p < 0.0001, p < 0.0001, permutation test, one-sided). Finally, clusters 1 (immature preadipocytes), 10 (heterogenous preadipocytes), and 12 (proliferating cells) were enriched for cells of progenitor and preadipocyte origins (progenitor origin: p < 0.0001, p < 0.0001, p < 0.0001; preadipocyte origin: p < 0.0001, p = 0.0282, p < 0.0001, respectively, permutation test, one-sided; Fig. 2G, fig. S5H-I, table S6). Thus, we can conclude that cells of progenitor origin have greater potential to reach the committed preadipocyte state than that of cells of preadipocyte origin. These findings were surprising given that the current adipogenesis model proposes a step-wise differentiation process where progenitors give rise to preadipocytes, which in turn terminally differentiate into mature adipocytes *(35)*. Yet, cells of progenitor origin displayed a higher potential to generate committed preadipocyte cells, a cell state with high *Pparg* and *Cebpa* expression (master regulators of the terminal differentiation process), while pre-existing preadipocytes, remained as or generated additional immature preadipocytes and lower numbers of committed preadipocytes. This suggests that progenitors and preadipocytes have unique differentiation potentials and that progenitors attain a privileged state that allows them to become committed preadipocytes more efficiently than pre-existing preadipocytes.

### Sox9 promotes the transition from adipocyte progenitor to preadipocyte

The molecular mechanisms that regulate adipocyte progenitor identity and how these cells become preadipocytes remain poorly understood. Merrick *et. al. (14)* identified TGFB signaling as a key factor in maintaining inguinal adipocyte progenitor identity *in vitro (14)*. To evaluate whether the same signaling mechanism is important to maintain the identity of skin adipocyte progenitors, we isolated skin adipocyte progenitors, cultured them with TGFB for five days, and harvested them for RNA isolation. Gene expression of selected progenitor markers by qPCR showed reduced or no change in expression after TGFB treatment (fig. S6A). Additionally, the expression of preadipocyte marker genes in progenitors treated with TGFB showed no significant change except for *Igf1*, which showed a five-fold increase with TGFB treatment (fig. S6A). To assess whether *in vitro* culture obscures the role of TGFB in maintaining skin adipocyte progenitor identity, we tested the impact of TGFB treatment *in vivo*. For this, we isolated adipocyte progenitors, CellTagged them, and treated them with TGFB for 5 days before transplanting them using a Matrigel matrix mixed with TGFB. Cells were recovered 11 days after transplantation and stained with antibodies to determine whether transplanted progenitors displayed progenitor or preadipocyte markers. Like previous experiments, control progenitors (treated with BSA) showed a significant degree of cells expressing DPP4, a progenitor marker (fig. S6B and C). TGFB-treated adipocyte progenitors lost DPP4 expression (fig. S6B and C), suggesting that the effect of TGFB on skin adipocyte progenitors has little to do with maintaining their identity.

Further analysis of our CellTag-adipogenesis dataset (Fig. 2B) showed high expression of the transcription factor *Sox9* in clusters 3 and 9, which are exclusively enriched for adipocyte progenitors, and among CellTagged progenitors (Fig. 3A-B). We therefore hypothesized that *Sox9* could be a regulator in the transition from adipocyte progenitor to preadipocyte. To test this, we isolated adipocyte progenitors and CellTagged them with either an empty library or with one that included *Sox9* under the control of the SFFV promoter. Five to seven days after CellTagging, cells were transplanted, separately, into mouse skin and recovered 11-13 days after transplantation (Fig. 3C). Flow analysis of surface markers shows that adipocyte progenitors that overexpress *Sox9* transition into a preadipocyte phenotype (CD9^high^ expression) in a higher proportion than GFP-only expressing cells, suggesting that *Sox9* is sufficient to promote adipocyte progenitor differentiation (progenitor origin: p = 0.00061; preadipocyte origin: p = 0.00125, unpaired t test with Welch’s correction, two-tailed; Fig. 3D-E). Interestingly, using the same *in vivo* approach with shRNA to knockdown *Sox9* expression (Fig. 3F and fig. S6D) resulted in an increased proportion of progenitor cells; however, adipocyte progenitors were still able to transition into the preadipocyte phenotype, suggesting that *Sox9* is not required by progenitors to transition to the preadipocyte identity (progenitor origin: p = 0.02924; preadipocyte origin: p = 0.70279, unpaired t test with Welch’s correction, two-tailed; Fig. 3G-H).

**Fig. 3.**
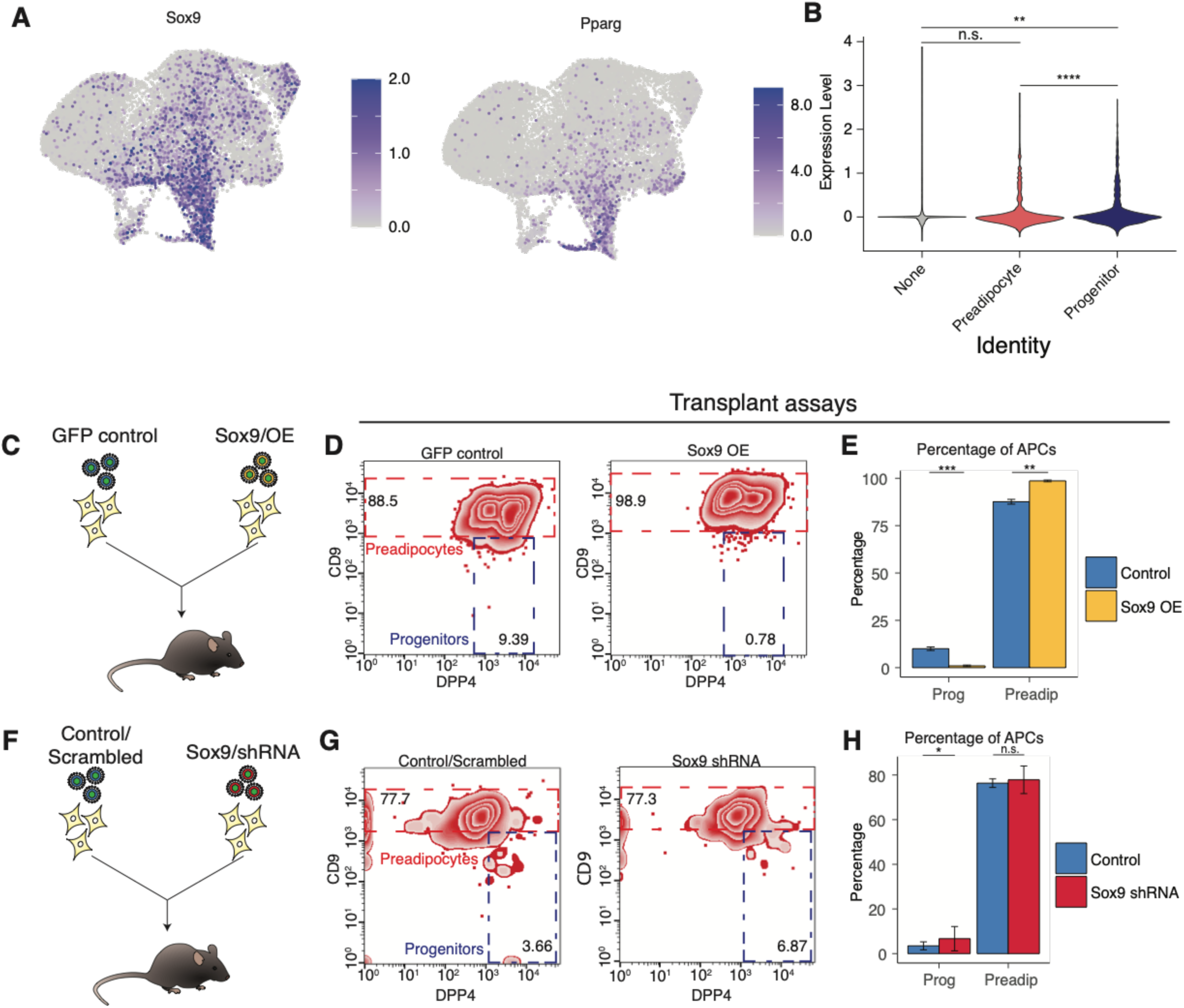
Sox promotes the transition from progenitor to preadipocyte. (A) Feature expression of *Sox9* and *Pparg* in the UMAP space of the P32 APC dataset. **(B)** Violin plot of *Sox9* feature expression by CellTag population (unpaired t test with Welch’s correction, two-tailed, ****p < 0.0001, **p < 0.01). **C.** Graphic representation of *Sox9* OE experiment in progenitor cells. **(D) and (E)** Representative FACS plots (D) and quantification (E) of transplanted progenitors from male and female C57BL/6J mice transduced with a *GFP*-only (control) or with *GFP*/*Sox9* expression cassette 11-14 days after initial injection into the skin of male C57BL/6J mice (n = 3 biological replicates, unpaired t test with Welch’s correction, two-tailed, ***p < 0.001, **p < 0.01). **(F)** Graphic representation of Sox9 knockdown experiment in progenitor cells. **(G) and (H)** Representative FACS plots (G) and quantification (H) of transplanted progenitors transduced with a *GFP* and scramble shRNA (control) or with *GFP* and *Sox9* shRNA expression cassette 11-14 days after initial injection into the skin of male C57BL/6J mice (n = 3 biological replicates; unpaired t test with Welch’s correction, two-tailed; *p < 0.05).

### Sox9 preferentially drives progenitor cells towards preadipocyte commitment

To gain a better understanding of the specific fate choices of progenitors driven by *Sox9* overexpression and knockdown, we performed an *in vivo* competition assay. For this, we isolated progenitor cells as previously described and divided them into three groups: one group was transduced with a CellTag construct that contained a *Sox9* expression cassette, another group received a GFP-only CellTag and a *Sox9* shRNA cassette, and the third group received a GFP-only CellTag and a scrambled shRNA control cassette. These groups were mixed in equal numbers and injected into the skin of P21 mice and harvested for scRNA-seq 11 days later (Fig. 4A). With two independent biological replicates, we were unable to recover an adequate number of shRNA (2) or control (144) CellTagged cells in our sequencing data. We hypothesized that *Sox9* overexpression may confer a competitive advantage in this system. To test this, we conducted an EdU proliferation assay with the same treatments *in vitro* (Fig. 4B). We observed that cells with the overexpression cassette showed a slight but significant increase of EdU incorporation (49.8%) compared to the control (43.2%), while cells with low *Sox9* expression (*Sox9*-shRNA-treated) displayed extremely low levels of EdU incorporation relative to the control (9.7%; Fig. 4C). This may further explain the discrepancy in cell recovery for this experiment. Using these data to model relative abundances in a competition assay, we determined that cells with the *Sox9* overexpression cassette would outcompete both other populations to abundances below 10% by 16 doubling periods (Fig. 4D).

**Fig. 4.**
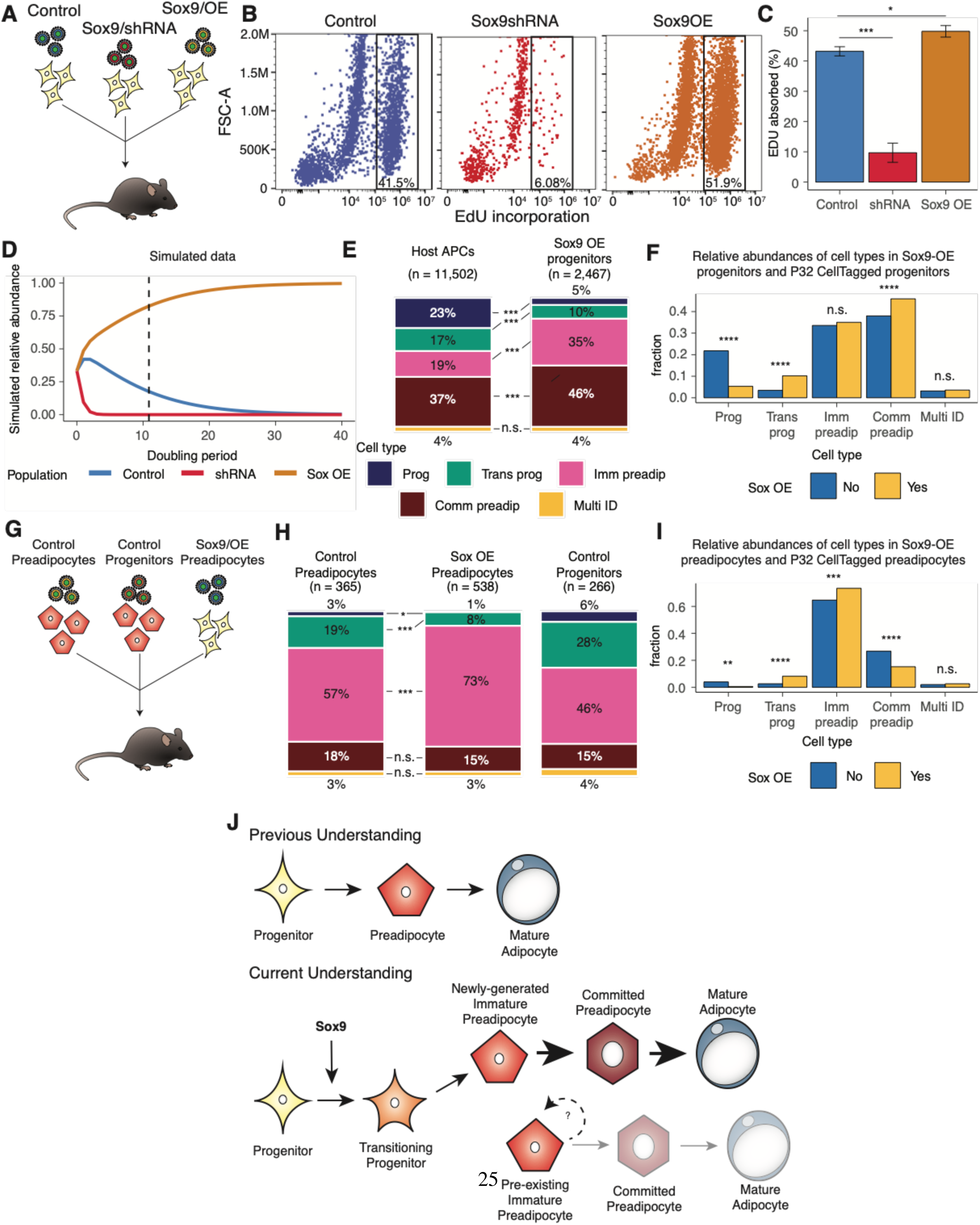
Sox9 expression drives progenitor cells into preadipocyte phenotype. (A) Graphic representation of *Sox9* overexpression/knockdown experiment in progenitor cells isolated from male and female C57BL/6J mice. **(B)** FACS plots of *in vitro* EdU incorporation in isolated progenitor cells treated with: control, *Sox9* overexpression (OE), and *Sox9* knockdown on day 5 after transduction. **(C)** Average percentage of EdU positive progenitors in control, *Sox9* overexpression (OE; *p < 0.05, unpaired t-test with Welch’s correction), and *Sox9* knockdown treatment groups (***p < 0.001, unpaired t-test with Welch’s correction; n = 3 biological replicates) **(D)** Abundance competition model generated with proliferation data to predict when dominance of *Sox9* OE cells could occur. Dashed line indicates approximate point of collection, 11 days. **(E)** Capybara-assigned identity proportions of host and *Sox9* OE progenitors based on P21 adipocyte precursor identities (*** p < 0.0001, randomization test, two-tailed). **(F)** Proportion of progenitor-originating cells identified as each cell type by Capybara, using our P21 dataset as reference for *Sox9*-overexpressed preadipocytes and progenitor CellTagged cells collected previously at P32 (Fig. 2; Chi square test; ****p < 0.0001). **(G)** Graphic representation of *Sox9* OE experiment in preadipocyte cells. **(H)** Capybara-assigned identity proportions of control progenitors, control preadipocytes, and *Sox9* OE preadipocytes. Colors correspond to legend in (E). (**E**) Randomization testing was performed to test for enrichment of cell types in *Sox9* OE preadipocyte population relative to control preadipocyte population (***p < 0.0001, *p < 0.01, randomization test, two-tailed). **(I)** Proportion of preadipocyte-originating cells identified as each cell type by Capybara, using our P21 dataset as reference for *Sox9*-overexpressed preadipocytes and CD9+ CellTagged cells collected previously at P32 (Fig. 2; Chi square test; ****p < 0.0001, ***p < 0.001, **p < 0.01). **(J)** Depiction of step-wise differentiation process where cells that differentiate are replaced by new cells from progenitors following the same track (Upper). Depiction of proposed differentiation process where activation of *Sox9* during the progenitor state increases potential for cells to transition toward the committed preadipocyte state. Pre-existing preadipocytes do not exhibit the same potential toward the committed lineage.

We observed 2,467 cells (18%) for which *Sox9* was overexpressed. Using Capybara with our P21 dataset and new classification as reference, we compared relative abundances of cell types among our CellTagged and host cells. In agreement with our flow cytometry data, we observed an enrichment of immature and committed preadipocytes in the *Sox9* overexpressing population relative to the host population (p < 0.0001, Pearson’s Chi-Square test with Yates’ continuity correction), as well as a reduction in the relative abundances of progenitor and transitioning progenitor cells (p < 0.0001, p < 0.0001, randomization test; Fig. 4E, table S5). 56% of the host population are immature or committed preadipocytes, while 81% of the *Sox9* OE CellTagged population are comprised of these cells. We confirmed these classifications by projecting these data onto our P32 CellTag-adipogenesis dataset, and observed strong co-clustering for each cell type (fig. S7A-B). Given the poor numbers of control progenitor cells in the competition assay, we also compared *Sox9* OE progenitors to progenitors transplanted in our P32 dataset (Fig. 2). This comparison showed a significant decrease in the number of progenitors (p < 0.0001, Pearson’s Chi-Square test with Yates’ continuity correction) and an increase in committed preadipocytes in *Sox9* OE cells (p < 0.0001, Pearson’s Chi-Square test with Yates’ continuity correction; Fig. 4F).

The data we present here shows diverging differentiation potential of progenitors and preadipocytes in our P32 dataset, in which progenitors acquired immature and committed preadipocyte phenotypes more frequently than pre-existing preadipocytes. Further, we showed that *Sox9* plays a key role in promoting the committed phenotype among progenitors. Therefore, we wanted to understand if the effects of *Sox9* observed in progenitor cells can also switch the fate of pre-existing preadipocytes from immature to committed. For this we isolated progenitors and preadipocytes from the skin as previously described. Preadipocytes were CellTagged and transduced with the *Sox9* overexpression cassette, and we additionally CellTag indexed progenitors and preadipocytes with a GFP-only construct as control populations (Fig. 4G). Here, we did not observe the same strong competitive advantage that *Sox9* conferred to progenitors, as we were able to recover cells from all 3 conditions in significant numbers (Fig. 4H). Based on our CellTag indices, we recovered 538 preadipocytes in which *Sox9* was overexpressed, 365 control preadipocytes in which *Sox9* was not overexpressed, and 266 CellTagged control progenitors. Among preadipocytes, we observed an enrichment for immature preadipocytes (p < 0.0001, randomization test; table S5) and a nonsignificant difference of committed preadipocytes in *Sox9* overexpressed cells relative to CellTagged preadipocyte controls (p = 0.2693, randomization test; Fig. 4H, table S5). Comparing CellTagged control preadipocytes from our P32 dataset (Fig. 2) to *Sox9* overexpressing preadipocytes also shows an increase in the relative abundance of immature preadipocytes (p = 0.0006, Pearson’s Chi-Square test with Yates’ continuity correction), with a depletion in the relative abundance of committed preadipocytes (p < 0.0001, Pearson’s Chi-Square test with Yates’ continuity correction; Fig. 4I, fig. S7A, C). These results suggest that *Sox9* overexpression does not alter the preadipocyte differentiation potential. Altogether, our results indicate that *Sox9* acts specifically on progenitor populations to induce differentiation toward immature and committed preadipocytes (Fig. 4J). *Sox9* overexpression in preadipocytes is unable to shift the fate of these cells toward a committed phenotype.

## Discussion

In this study, we provide evidence of a previously uncharacterized immature preadipocyte population in the skin, the divergent differentiation potential of progenitors and preadipocytes, and the role of *Sox9* in promoting the differentiation of progenitor cells into both immature and committed preadipocytes. While single-cell approaches haven been used in the past to understand the biology of fibroblasts and adipocyte precursors in the skin during homeostasis, disease, and repair *(34, 36–39)*, this is the first effort to our knowledge to individually lineage-trace the differentiation paths of adipocyte progenitors and preadipocytes in the skin during homeostatic differentiation using a combination of genetic barcoding and scRNA-seq. Lineage-tracing technologies have experienced a rapid evolution, especially when combined with scRNA-seq *(40)*. Our CellTag approach has the advantage of being able to label specific populations and assay their differentiation potential in the same microenvironment. While isolation and manipulation of cells *in vitro* is not the perfect approach, it is currently the most precise way to target progenitors and preadipocytes in a specific manner and its overall results are comparable to similar transplant assays carried out without in vitro manipulation *(14)*. It will be interesting to identify genetic elements that drive gene expression specifically in progenitor or preadipocyte populations and use *in vivo* barcoding, such as the CARLIN mouse model *(41)*, to follow differentiation trajectories of adipocyte precursor cells in different tissues and under different biological conditions.

We also provide the first dermal-inguinal comparison of adipocyte precursors and assigned dermal adipocyte precursors identities using our cell classifier Capybara. This approach allowed us to identify a prominent immature preadipocyte population in the skin which was previously overlooked given its low prevalence in the inguinal adipose depot *(14)*. Interestingly, adipocyte progenitors from both the skin and inguinal depots display a high degree of shared identity. This could suggest that progenitors in the skin and in the inguinal depot share a common or very similar microenvironment. There is evidence of a somewhat homogeneous mesenchymal population in a wide array of tissues that expresses high levels of *Dpp4* and stemness markers *(42)*, which could indicate that the adipocyte progenitor population is present in many other tissues and that they give rise to more than one type of specialized mesenchymal cell. While there is evidence of intrinsic differences between adipose precursors from different depots *(43, 44)*, the differences between skin and inguinal preadipocytes could also point to a differentiation process that occurs in dissimilar environments *(45)* or responds to different needs.

One of the most obvious differences between skin and inguinal preadipocytes is the different proportion of immature and committed cells between depots. The skin preadipocyte population only contains a minor proportion of committed preadipocytes, while the committed population is very prominent in the inguinal depot. These differences could be partially explained by the ever-changing nature of the dermal adipose depot that undergoes natural cycles of expansion and regression under the influence of the hair follicle cycle *(46, 47)* compared to the relatively quiescent state of the inguinal depot under homeostatic conditions where only a small proportion of new mature adipocytes are renewed *(48)*. It is possible that the presence of immature preadipocytes in the skin responds to the need to generate new mature adipocytes in a cyclic manner while committed preadipocytes could be geared towards responding to a sudden demand for energy storage. Understanding the differences in adipogenesis between the skin and other depots could produce the necessary knowledge to promote generation of mature adipocytes to prevent hypertrophy-associated negative effects *(49)*.

In this study, we also provide proof that progenitors and preadipocytes have divergent differentiation potential. We identify these differences by performing the first reported competitive assay between adipocyte progenitors and preadipocytes. This surprising finding suggests that, contrary to the step-wise differentiation paradigm of mature adipocytes, progenitor cells are more capable of reaching a committed preadipocyte state than preexisting preadipocytes. At the same time, preexisting preadipocytes appear to especially accumulate in immature states different from immature preadipocytes generated from progenitor cells (Fig. 3E). These findings raise questions about the replenishment of preadipocytes by progenitor cells following differentiation. Could it be possible that preadipocytes have some self-renewing capabilities and do not completely rely on progenitors to maintain their population? When the skin is subject to depilation (an injury model), preadipocytes are activated to proliferate *(23)*. Examples of populations that do not rely on more stem-like progenitors to be maintained occur in hematopoiesis and epithelial differentiation *(50–52)*. Additionally, given the high capacity of progenitor cells to reach the committed preadipocyte stage, it might be possible that homeostatic and induced (excess energy) adipogenesis could promote selective differentiation of progenitors or preadipocytes populations. Moreover, an alternative explanation for this different biological potential could be that the isolated preadipocyte population is heterogenous and contains a mixture of adipogenic and non-adipogenic mesenchymal cells.

Finally, we identify *Sox9* as a key transcription factor capable of promoting the differentiation of progenitors into mature and committed preadipocytes but it has little to no effect on the differentiation of pre-existing preadipocytes, suggesting that the effect of Sox9 is cell-state specific. This is, to our knowledge, the first description of a molecular mechanism that controls differentiation of adipose progenitors into preadipocytes. *Sox9* has been shown to regulate chondrocyte differentiation in a stage-specific manner *(53)* and to induce proliferation following injury in the kidney *(54)*. In adipose tissue, *Sox9* has been shown to prevent terminal differentiation in *Pref1/Dlk1*+ inguinal cells *(55)*. This population of inguinal *Dlk1*+ cells most likely consists of a majority of committed preadipocytes according to scRNA-seq data, therefore it remains unclear if the role of *Sox9* in skin progenitors is specific to this depot or a general mechanism.

In summary, using scRNA-seq and Capybara, a unique tool for cell classification, we have identified a previously uncharacterized population of immature preadipocytes in the skin. Barcoding of cells with our CellTag libraries provided unprecedented cell-tracking resolution, allowing us to identify the existence of divergent differentiation potential between progenitors and preadipocytes, and identified *Sox9* as a key factor driving the differentiation of adipose progenitor cells. These findings show the existence of a previously unknown stable adipogenic state, provide detailed biological information on the process of generation of new mature adipocytes and could be used to design therapeutic approaches that aim to increase mature adipocyte production in specific adipose depots. This would help controlling local adipose tissue inflammation, mature adipocyte size and its associated negative effects *(5)*.

## Acknowledgements

We thank members of the Morris laboratory for helpful discussions, the Washington University Center for Cellular Imaging (WUCCI) at Washington University School of Medicine, and the Alvin J. Siteman Cancer Center at Washington University School of Medicine and Barnes-Jewish Hospital in St. Louis, MO., for the use of the Siteman Flow Cytometry, which provided FACS service.

## Funding

National Institute of General Medical Sciences R01 GM126112 (SAM)

Silicon Valley Community Foundation, Chan Zuckerberg Initiative Grant DAF2021-238797 (SAM)

Allen Distinguished Investigator Award through the Paul G. Allen Frontiers Group (S.A.M.) Vallee Scholar Award (SAM)

Sloan Research Fellowship (SAM)

New York Stem Cell Foundation Robertson Investigator Award (SAM)

Rita Levi-Montalcini Postdoctoral Fellowship from the Center of Regenerative Medicine, Washinton University in St. Louis (GCRG)

NIH-T32HG000045-21 (EGB)

NIGMS-R01GM126112-03S1 supplement (CEG)

## Author contributions

Conceptualization: SAM, GCRG Methodology: SAM, GCRG, EGB

Experimental procedures: GCRG, EGB, CEG, KJ Data Analysis: SAM, GCRG, EGB, CEG, WK Funding acquisition: SAM

Writing – original draft: SAM, GCRG, EGB, CEG

## Data and materials availability

All single-cell RNA sequencing data will be deposited on the Gene Expression Omnibus (GEO; accession number to be provided). All code used for analysis will be deposited on Github (link to be provided).

## Supplementary Materials

Materials and Methods

Figs. S1 to S7

Tables S1 to S6

References *(14, 23, 31, 56, 57, 58, 59, 60)*

