## Supplementary material for "Single-cell lineage tracing reveals hierarchy and mechanism of adipocyte precursor maturation": Methods and supplementary information

**The PDF file includes:**

Materials and Methods
Supplementary Text
Figs. S1 to S7
Tables S1 to S6
References

### Materials and Methods

Mice

All experiments conducted on mice were done following the guidelines issued by Washington University in St. Louis' Institutional Animal Care and Use Committee (IACUC) Protocol ID: 21-0317. C57BL/6J (Strain #:000664) and ROSA$^{nTnG}$ (Strain #023035) mice were purchased from Jackson Laboratories and used at least 1 week after their arrival.

Immunofluorescence and Tissue Staining

*C57BL/6J* mouse skin was embedded in optimum cutting temperature (OCT) compound (Tissue-Trek, 4583) and frozen on dry ice. OCT blocks were cryosectioned at 14 µm and fixed for 10 min with 4% formaldehyde. Skin sections were stained as previously described *(*23*)*. The following antibodies were used: Perilipin A (ab3526, 1:1000), tdTomato (goat; LSBio [LS-C3406] 1:100). After staining, slides were incubated in DAPI solution (300nM) for 5 min and washed. Sections were mounted in Prolong Gold anti-fade reagent (Thermo Fisher, P36934). Images were taken with a Zeiss AxioImager Z2. Image analysis was done using Fiji software (NIH) and Adobe Photoshop.

Adipocyte Precursor Cell Culture and differentiation

FACS-isolated adipocyte precursors were cultured as described previously from P21 male and female C57BL/6J mice (*23*). Briefly, cells sorted via FACS were plated on collagen-coated plates (Collagen I, Cat# A1048201, Thermo Fisher) in DMEM supplemented with 10% fetal bovine serum (FBS). For differentiation experiments, cells were allowed to reach 100% confluency then switched to 0.5% FBS DMEM for 48h. After starving, cells were treated with insulin (Sigma, St. Louis, MO, 1882, 2 µg/ml) every 48h until collection time. Cells were treated with Bodipy (1:1,000; Thermo Fisher, D-3922) and imaged using a Nikon eclipse Ts2 inverted microscope before RNA isolation.

RNA Extraction and Real-Time PCR

RNA extraction and purification were done using the RNeasy mini kit (QIAGEN, 74104) or RNeasy micro kit (QIAGEN, 74004), following the manufacturer's instructions. RNA was reverse transcribed using the Maxima RT kit (ThermoFisher K1672). Real-time PCR was performed uisng TaqManTM Gene Expression Master Mix (ThermoFisher Scientific) and gene-specific TaqManTM probes in a 20ul reaction volume and processed according to manufacturer's instructions (4371135) on the StepOne Plus qPCR system. Probes: Akr1c18, (Mm00506289_m1), Smpd3 (Mm00491359_m1), Gap43 (Mm00500404_m1), Dpp4 (Mm00494538_m1), Cd9 (Mm00514275_g1), Igfbp7 (Mm03807886_m1), Mfap4 (Mm00840681_m1), Igf1 (Mm00439560_m1), Eln (Mm00514670_m1), Lox (Mm00495386_m1), Smoc2 (Mm00491553_m1), Plin1 (Mm00558672_m1), Adipoq (Mm00456425_m1), Pparg (Mm00440940_m1), Sox9 (Mm00448840_m1).

FACS and Analysis

FACS analysis of adipocyte precursors was performed as described previously *(23)*. Briefly, male and female P21 *C57BL/6J* mouse skin was dissected and digested in collagenase buffer (Hank's balanced salt solution [HBSS] containing 3% BSA, Collagenase 1A 1:100, Worthington LS004196, 1.2 mM calcium chloride, and 0.8 mM zinc chloride) for 60 min at 37°C in a shaking water bath. Undigested tissue was separated from released cells through filtration using 100 and 70-µm filters. Floating mature adipocytes were separated from the stromal vascular fraction (SVF) by centrifugation at 300× g for 3 min. To identify adipocyte precursors, the SVF was stained in 3% BSA in HBSS with the following antibodies: CD31-PE-Cy7 (1:500; BD Biosciences, 561410), CD45 APC-PE-Cy7 (1:5,000; BD Biosciences, 561868), CD29 PE/Dazzle 594 (1:400; BioLegend, 102232), CD34 PE (1:200; BioLegend, 119301), DPP4 (CD26) APC (1:500; BioLegend, 137807), CD9 BV421 (1:250; BD Biosciences, 564235) and F3 (1:100, AF3178 R&D). For live/dead discrimination, cells were stained with Live/Dead Fixable Violet (1:1,000; Thermo Scientific, L23105). For analysis of proliferation by EdU incorporation, the Click-iT EdU Flow Cytometry Assay Kit (Invitrogen, C10419) was used, following the manufacturer's instructions. Samples were sorted or analyzed with a Sony iCyt Synergy BSC or a CytoFLEX SRT machine. Analysis of flow cytometry data was performed using FlowJo software.

CellTag and plasmids

CellTag v1 and v2 lentiviral constructs were generated by introducing an 8-bp variable region into the 3′ UTR of GFP in the pSmal plasmid (*57*) using a gBlock gene fragment (Integrated DNA Technologies) and megaprimer insertion (https://www.addgene.org/pooled-library/morris-lab-celltag/). Cebpa-DN (Addgene, 33352) was cloned into the CellTag A plasmid (Addgene, 24591). Individual clones from CellTag-multi (*31*) were picked and Sanger sequenced to generate predefined barcodes. Sox9 was cloned into into CellTag_F (ATAGTATTCTGACAGGTATGAGCCATCT) under the control of the SFFV promoter, Sox9-shRNA-treated cells were tagged using CellTag_D (TAGGTGTGCTATTAGATATGTCACATAG) and control cells were tagged using CellTag_A (TTCGTAGCCTGTCAGCTATGGTTCATAG).
pLKO.1-sh-mSOX9-5 (40646) and pLKO.1 Puro shRNA Scramble (162011) were obtained from Addgene.

Lentivirus production

CellTag lentiviruses were produced by transfecting HEK293T cells with lentiviral pSMAL vector and packing plasmids pCMV-dR8.2 dvpr (Addgene plasmid 8455) and pCMV-VSV-G (Addgene plasmid 8454) using X-tremeGENE 9 (Sigma-Aldrich). ShRNA lentiviruses were produced by transfecting HEK293T cells with pLKO.1 plasmids and packaging plasmids psPAX2 (Addgene, 12260) and pMD2.G (Addgene, 12259). Viruses were collected 48 and 72 h after transfection.

Virus transduction

CellTag, CEBP-DN, Sox9-shRNA and scrambled virus-containing supernatant collected from virus-producing HEK293T cells was kept at 4 °C and used within 1 week. Prior to transduction, protamine sulfate (Sigma-Aldrich) was added to the viral solution to a final concentration of 4 µg/ml. Cells were aspirated of media, and the virus was added to the cells in combination with complete media in 12-h transduction periods. This transduction was repeated as needed (2 incubations with CellTag, CEBP-DN, or shRNA viruses) for a total of 24 h for adipocyte precursors that received only one virus type or 48h for cells that received two different viruses. shRNA knockdown was confirmed by sampling cells and measuring Sox9 expression before transplantation into the skin.

Transplantation assay for ROSA$^{nTnG}$ adipocyte precursors

Adipocyte precursors were isolated by FACS from male and female ROSA$^{nTnG}$ mice as outlined above. Live/lineage negative cells were tested for CD34 expression and positive cells were then selected and tested for DPP4 and CD9 expression. DPP4$^{high}$/CD9$^{low}$ cells were isolated as progenitors and CD9$^{high}$ cells were further separated based on F3 expression. F3$^{high}$ cells were isolated as immature preadipocytes and F3$^{low}$ cells isolated as committed preadipocytes. Cell populations (250k) were resuspended in PBS and mixed with Matrigel (Corning #356231) (1:1 volume), and individually injected intradermally into the back skin of P20-21 C57BL/6J mice. Tissue was collected for sectioning and staining at P32.

Transplantation Assay for virus-treated cells

Adipocyte precursors were isolated by FACS from P21 C57BL/6J male and female mice, treated with corresponding viruses, resuspended in PBS and mixed with Matrigel (Corning #356231) (1:1 volume), and injected intradermally into the back skin of P21 C57BL/6J mice. For progenitor/preadipocyte competition assays (v1/v2 CellTag libraries) each recipient mouse was injected with 400k-500k total cells (50% v2 progenitors and 50% v1 preadipocytes). For non-competitive *Sox9* overexpression and knockdown transplants, 300k-400kprogenitor were injected into the skin. In *Sox9* competitive assays, 200k cells from each condition were injected into the skin. At P32-35 the skin of host mice was analyzed for the presence of both host (GFP-) and transplanted (GFP+) cells, using the FACS strategy described previously we isolated GFP+ and GFP- total cells (no antibody labeling) for the competitive P32 datasets and mixed host and transplanted cells in equal numbers for single-cell library preparation. For the *Sox9* non-competitive overexpression and knockdown assays, GFP+ and GFP- adipocyte precursors were identified using CD31/45, CD34, DPP4 and CD9 as markers. For the *Sox9* competitive overexpression assays, GFP+ and GFP- adipocyte precursors were isolated by sorting CD31/45$^{neg/}$CD34$^{high}$ cells and mixing host and transplanted cells in equal numbers for single-cell library preparation.

Single-nucleus isolation from P32 skin

Nuclei isolation was performed on the skin of C57BL/6J 2 male mice as described in (*58*).

Single-cell and nucleus library preparation and sequencing

Single-cell and single-nucleus library preparation was performed using the Chromium Single-Cell Gene Expression Kit from 10X Genomics. Libraries were sequenced using an Illumina NextSeq-500.

CellTag amplification for scRNA-seq (CellTag-RNA PCR)

A PCR step was used to amplify CellTag barcodes from the single-cell cDNA library of Sox9OE and knockdown assays, obtained after step 2.4 of the 10x Genomics Single Cell Gene Expression Kit user guide (CG000315). 5ul (or at least 60ng) of cDNA was mixed with 2x Q5 HF PCR Master Mix (New England Biolabs) and 500nM of *P5/R1-par* and *P7/SI-R2* primers in a 50ul reaction volume and subjected to the following PCR program: 98 C for 30 seconds; N cycles (98°C for 10 seconds; 54°C for 30 seconds; 72°C for 30 seconds); 72°C for 2 minutes. The number of PCR cycles (N) was kept the same as the number of cycles used during sample index PCR of the main scRNA-seq library. CellTag amplicon library was purified using double-sided bead purification (0.4x-0.64x) and quantified on an Agilent TapeStation using the D1000-HS tape. Libraries were along with scRNA-seq libraries. CellTag amplicon libraries were sequenced on an Illumina NextSeq-500. Primer sequences: P5/R1-par 5'AATGATACGGCGACCACCGAGATCTACACTCTTTCCCTACACGACGCTC3' and P7/SI-R2_2 Indexed primer 5'CAAGCAGAAGACGGCATACGAGATNNNNNNNNGTGACTGGAGTTCAGACGTGTG CTCTTCCGATCTACAGgtactggagccgaga3'

Basic data alignment and processing

After demultiplexing, data were aligned using CellRanger versions 5.0.1, 6.0.2, 6.1.2, and 7.0.1 from 10X Genomics (https://support.10xgenomics.com/single-cell-gene-expression/software/downloads/latest). A custom reference genome was used, comprised of the mm10 genome as well as transgenes for GFP and the dominant-negative form of CEBP-A (DN-CEBPA). CellRanger-filtered digital gene expression matrices were generated for each sample for downstream analysis; bam files were simultaneously generated for CellTag processing. All single-cell data were processed using R 4.2.2 and Seurat 4.3.0. Using a standard Seurat analysis pipeline, we filtered out low-quality cells that had a high fraction of mitochondrial reads (greater than 0.05) and low total counts (less than 500; https://satijalab.org/seurat/articles/pbmc3k_tutorial.html). Cell cycle scoring and regression was performed. UMAP dimension reduction was performed with 10 principal components; UMAP and Louvain clustering algorithms were otherwise implemented with Seurat default parameters.

Data visualizations were generated using the R package ggplot2 3.4.2. Unless otherwise mentioned, statistical tests were performed using the R package stats 4.2.2.

Integration

P21 skin data were integrated with P12 inguinal data from Merrick et. al. (*14*) using the Seurat functions *FindIntegrationAnchors()* and *IntegrateData()* before further standard re-processing procedures. 4 P32 independent biological replicates were identically prepared and sequenced in pairs at two different times, so pairs of datasets sequenced together were integrated using the Seurat functions *FindIntegrationAnchors()* and *IntegrateData()*. Integrated objects were re-scaled, re-clustered, and re-embedded. Louvain clusters expressing high levels of features identified in Supplementary Table 3 were removed from both datasets, while clusters expressing high levels of adipocyte precursor markers or GFP were retained. Finally, both datasets were integrated again using the same Seurat integration procedure. The same re-embedding and clustering procedures were re-implemented downstream of integration on the new object. Finally, Sox9 perturbation datasets were projected onto the merged P32 object using the Seurat functions *FindTransferAnchors()* and *MapQuery()*.

CellTag index extraction

A standard workflow was utilized in indexing experiments to perform clone calling, with "clones" approximated as cells with shared CellTag index (https://github.com/morris-lab/newCloneCalling). Here, the bam file was parsed for CellTag reads using the "multi-v1" CellTag version. One CellTag per cell was considered an adequate CellTag signature in filtering due to the length of the barcode, and cells with greater than 20 counts of a CellTag were removed. Allowlisting and binarization were performed, but Jaccard and additional downstream clonal analyses were not performed due to the nature of these indexing experiments and lack of biologically informative clonal information.

Cell type classification by Capybara

We evaluated our datasets for automatic cell type classification with various references according to the published Capybara workflow and the R package Capybara (v0.0.0.9; https://github.com/morris-lab/capybara). We did not perform tissue-level classification; instead, raw counts were used for each dataset to generate a custom reference with classifications constructed as described in each case. Quadratic programming, discrete cell type classification, and multiple identity scoring were performed for downstream analysis.

Differential expression analysis

Differentially-expressed features were identified using the *FindMarkers()* function in Seurat. In all instances, default parameters were used: a default log fold-change required between the two groups of 0.25, a Wilcoxon Rank Sum test was used to identify features, and a minimum fraction of .1 cells expressing the feature in either population was required for testing.

Jaccard similarities and modified z score procedure

To calculate Jaccard similarities of Louvain clusters of our P32 data and Capybara classifications of progenitors and committed preadipocytes, we used the *PairWiseJaccardSets()* function of the R package scclusteval (v1.0) (*59*). Preserving numbers of cells classified as progenitor or committed preadipocyte and sizes of Louvain clusters, we randomized these classifications and calculated Pair-wise Jaccard distances of randomized data 1,000 times to generate a background distribution of Jaccard values for each pair of cell type classification and Louvain cluster. Because we could not confirm that each of these distributions was normal using a Shapiro Wilk test for normality (data not shown), we calculated a modified z-score for each Jaccard value against its corresponding background distribution to determine outliers in our correlation data (*60*). The modified z-score procedure considers the median and median absolute deviation (MAD) in place of the mean and standard deviation, respectively, that are used in a traditional z-score procedure. For a Jaccard value $J_i$, the modified z-score $z_i$ is calculated against a background distribution $D_i$ as:

$$z_i = \frac{0.6745 * \left(J_i - median(D_i)\right)}{MAD(D_i)}$$

Outliers are defined as values of $z$ such that $|z| > 3.5$. Further, 0.6745 is used as a constant to correct modified z-scores, as the expectation of the MAD is approximately $0.6745 \cdot \sigma$, or the standard deviation of a normally-distributed randomized background.

Abundance modeling

Using mean EdU absorption data for each sample of our first *Sox9* progenitor perturbation experiment as a proxy for constant proliferation rate in a single doubling or division period, we modeled relative abundance of each population to one another. With $m = 3$ samples each having a unique proliferation rate $p_s$ and initial relative abundances $a_{s,0} = 1/3$, we modeled the relative abundance of sample $s$ at time $t$ using a recursive sequence defined as:

$$a_{s,t} = \frac{p_s * a_{s,t-1}}{\sum_m^n p_n * a_{n,t-1}}$$

According to this model, we estimate that after t = 16 division periods, the total relative abundance of control and *Sox9* knockdown CellTagged cells is less than 10%, or 0.1.

Randomized testing of enrichment of CellTag populations and cell type classifications

To determine Louvain clusters of our P32 data that are enriched for one or both CellTagged population, we used a custom Python-based script described by Biddy, et. al. (*25*). To determine whether cell types are differentially-enriched among our CellTagged populations relative to one another or a control, we performed a randomization procedure of pooled cell type classifications for cells in each pair of populations. After pooling, we randomly reassigned cell types to one population or the other, with population sizes preserved. This was performed 10,000 times, generating a null distribution model representing the difference of fractions of each cell type to the population size in each pair of populations. Finally, a two-tailed p-value was calculated representing the number of the absolute values of these differences that are greater than or equal to the absolute value of the observed difference between the two populations. Bonferroni multiple testing correction was performed to control for dependent alternative hypotheses.

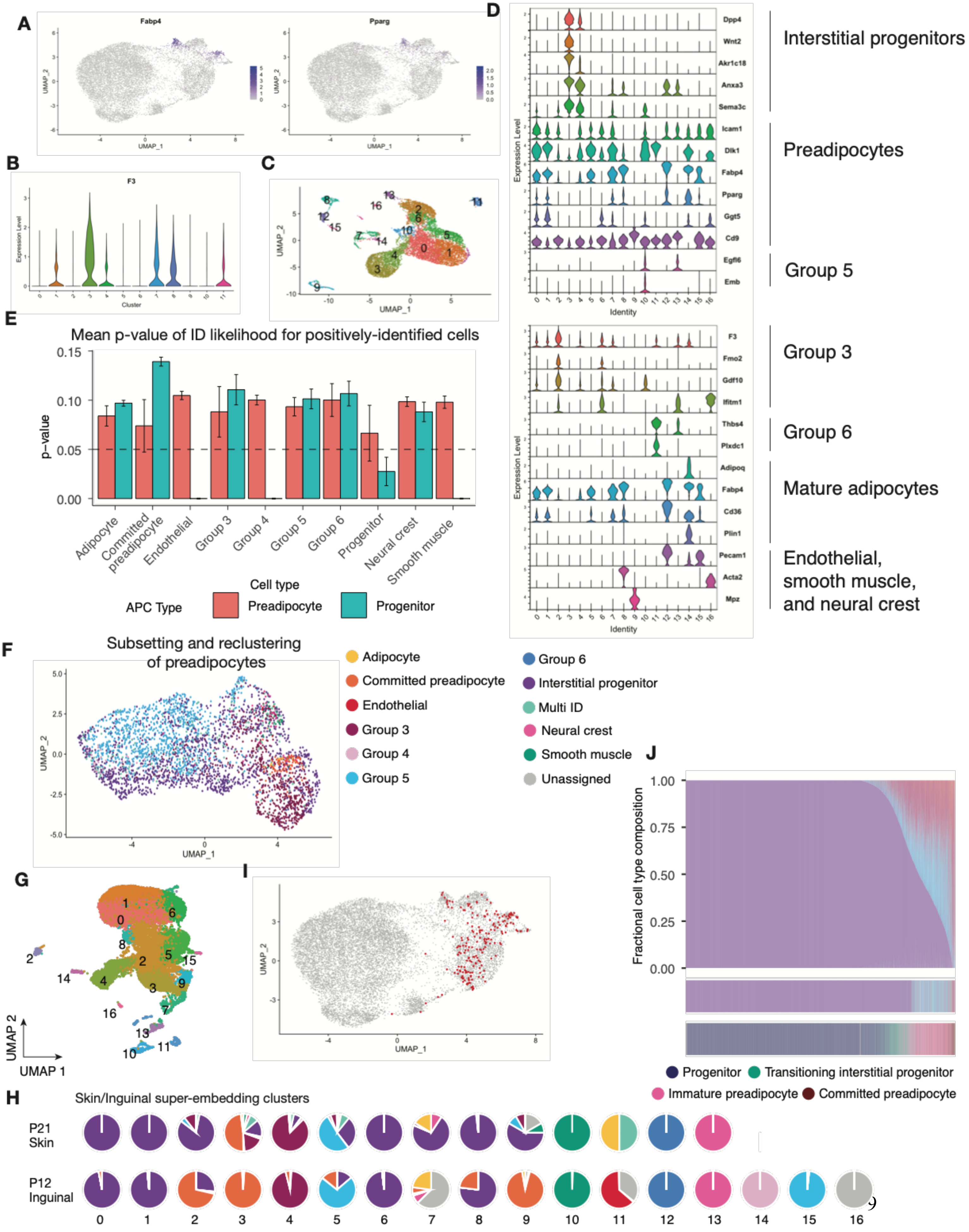
A
Fabp4
Pparg
UMAP_1
UMAP_2
B
F3
Expression Level
Cluster
C
UMAP_1
UMAP_2
D
Dpp4
Wnt2
Akr1c18
Anxa3
Sema3c
Icam1
Dlk1
Fabp4
Pparg
Ggt5
Cd9
Egfl6
Emb
Identity
Expression Level
Interstitial progenitors
Preadipocytes
Group 5
F3
Fmo2
Gdf10
Ifitm1
Thbs4
Plxdc1
Adipoq
Fabp4
Cd36
Plin1
Pecam1
Acta2
Mpz
Identity
Group 3
Group 6
Mature adipocytes
Endothelial,
smooth muscle,
and neural crest
E
Mean p-value of ID likelihood for positively-identified cells
p−value
0.15
0.10
0.05
0.00
Adipocyte
Committed preadipocyte
Endothelial
Group 3
Group 4
Group 5
Group 6
Progenitor
Neural crest
Smooth muscle
Cell type
APC Type
Preadipocyte
Progenitor
F
Subsetting and reclustering
of preadipocytes
UMAP_1
UMAP_2
Adipocyte
Committed preadipocyte
Endothelial
Group 3
Group 4
Group 5
Group 6
Interstitial progenitor
Multi ID
Neural crest
Smooth muscle
Unassigned
G
UMAP 2
UMAP 1
I
UMAP_1
UMAP_2
J
Fractional cell type composition
1.00
0.75
0.50
0.25
0.00
Progenitor
Transitioning interstitial progenitor
Immature preadipocyte
Committed preadipocyte
H
Skin/Inguinal super-embedding clusters
P21
Skin
P12
Inguinal

**Fig. S1. Characterization of skin adipocyte precursor cells. (A)** Feature expression of *Fabp4* and *Pparg* in P21 adipocyte precursor cells. **(B)** Violin plot of *F3* gene expression by cluster in P21 adipocyte precursor cells. **(C)** UMAP of lineage-depleted stromal vascular fraction (SVF) from inguinal adipose tissue (Merrick, D. *et. al.*) (*14*) and UMAP clusters in (C). **(D)** Manual cluster classification of SVF cells based on reported markers from Merrick, D. *et. al.* (*14*). **(E)** Capybara mean p-value of identity likelihood for positively identified preadipocytes and progenitors from the skin using the Merrick, D. *et al*. (*14*) dataset as the reference. **(F)** UMAP of subset and re-clustered skin preadipocytes (clusters 3, 7, 8, and 11). **(G)** UMAP super-embedding of inguinal and skin adipocyte precursors (Merrick, D. *et. al.*) (*14*) and our skin dataset. **(H)** Identity composition of clusters in the super-embedding object. Colors correspond to legend in (F). **(I)** UMAP of P21 skin adipocyte precursors highlighting the location of multi-ID cells. **(J)** Fractional identities of P21 skin APCs assigned by Capybara using inguinal dataset as reference (top), Capybara cell type classifications (middle) and new cell type classifications (bottom). Colors of top and middle panels correspond to legend in (F).

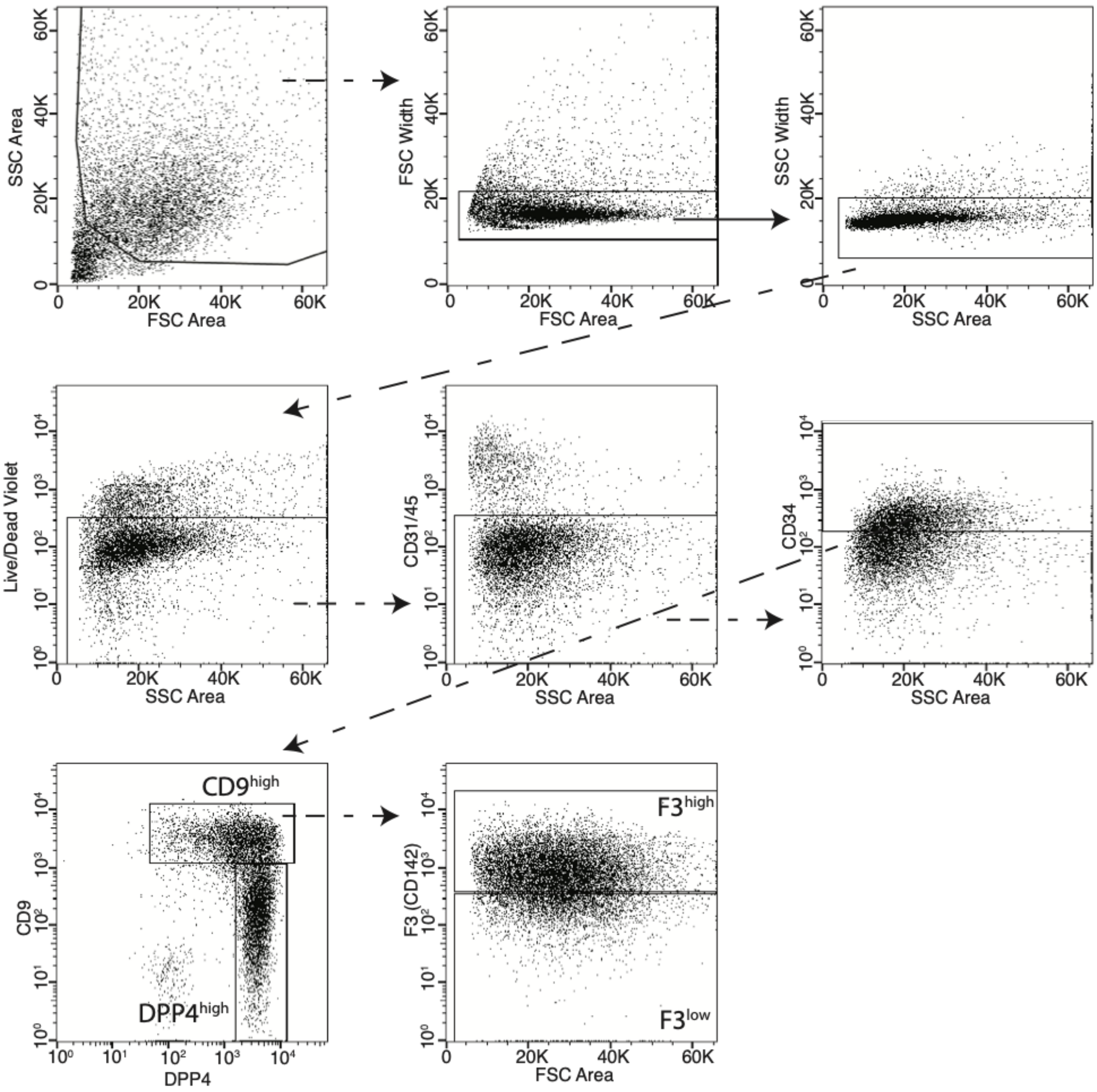
SSC Area
FSC Area
FSC Width
FSC Area
SSC Width
SSC Area
Live/Dead Violet
SSC Area
CD31/45
SSC Area
CD34
SSC Area
CD9
DPP4
CD9high
DPP4high
F3 (CD142)
FSC Area
F3high
F3low

**Fig. S2. FACS strategy to isolate progenitors, immature, and committed/group 3 preadipocytes from C57BL/6J P21 mouse skin.** Following singlet discrimination, live cells were selected and CD45/CD31 cells excluded from this population. Live/lineage negative cells were tested for CD34/CD29 expression and positive cells were then selected and tested for DPP4 and CD9 expression. $DPP4^{high}/CD9^{low}$ cells were isolated as progenitors and $CD9^{high}$ cells were further separated based on F3 expression. $F3^{high}$ cells were isolated as immature preadipocytes and $F3^{low}$ cells isolated as committed/group3 preadipocytes.

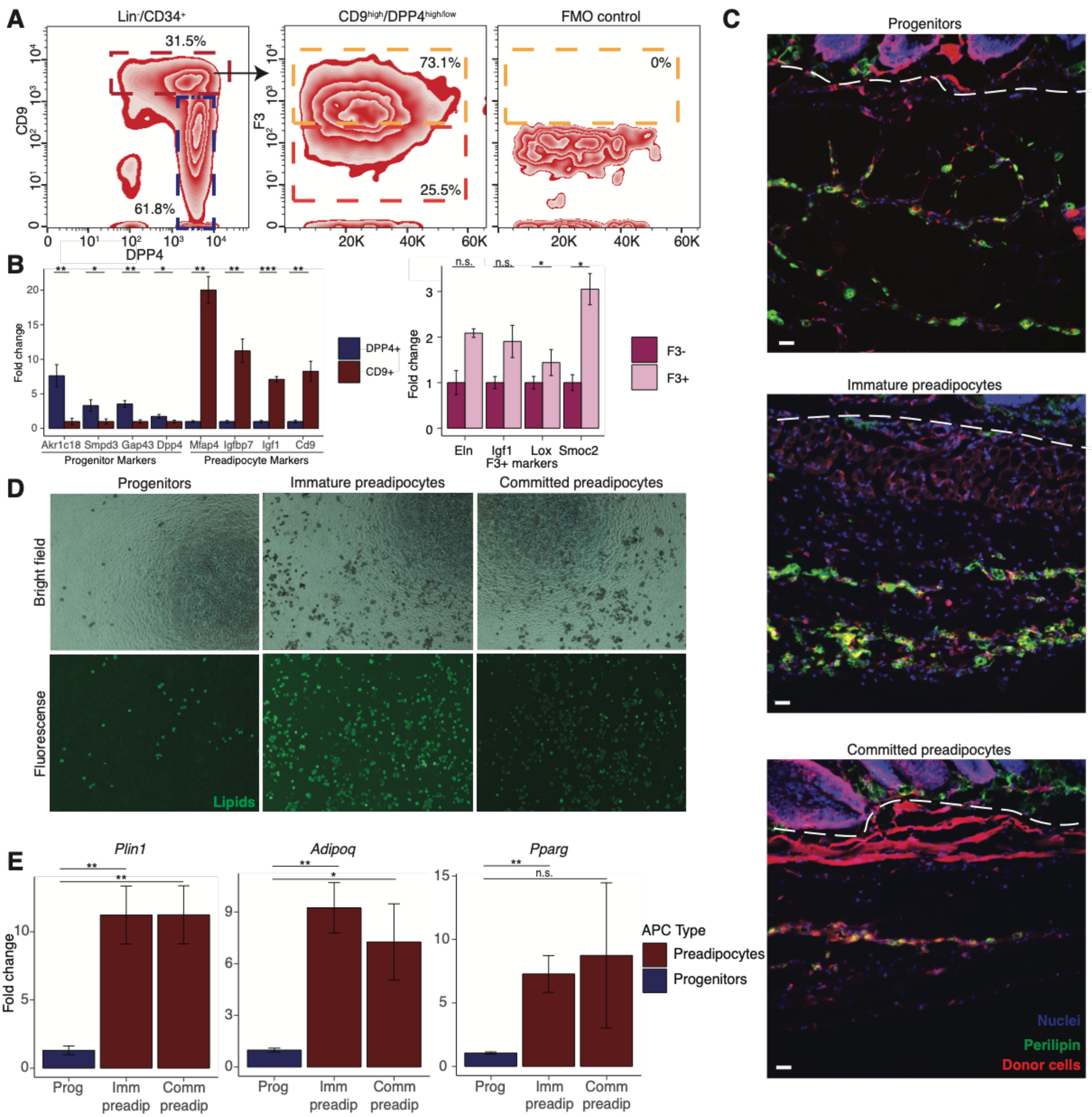
A
Lin-/CD34+
31.5%
61.8%
CD9
DPP4
CD9high/DPP4high/low
73.1%
25.5%
F3
FMO control
0%
20K
40K
60K
B
Fold change
Akr1c18 Smpd3 Gap43 Dpp4 Mfap4 Igfbp7 Igf1 Cd9
Progenitor Markers
Preadipocyte Markers
DPP4+
CD9+
n.s.
Eln Igf1 Lox Smoc2
F3+ markers
F3-
F3+
C
Progenitors
Immature preadipocytes
Committed preadipocytes
Nuclei
Perilipin
Donor cells
D
Progenitors
Immature preadipocytes
Committed preadipocytes
Bright field
Fluorescense
Lipids
E
Plin1
Adipoq
Pparg
n.s.
Fold change
Prog
Imm preadip
Comm preadip
APC Type
Preadipocytes
Progenitors

**Fig. S3. Establishing adipogenic potential of distinct APC populations. (A)** FACS strategy to isolate progenitors, group 5 cells, and committed preadipocytes/group 3 cells. Fluorescence minus one control shows gating for F3-positive cells. **(B)** Representative images of transplant areas in P32-34 mice 11-13 days after cell injection. Donor cells were isolated from the skin of male and female $ROSA^{nTnG}$ mice at P21. Sections were stained for TdTomato, Perilipin1, and DAPI for nuclei. The scale bar is 100µm (*n* = 3 mice for condition). Dotted line delineates the border between dermis and panniculus carnosus. **(C)** Marker gene expression of FACS isolated progenitors, group 5, and committed preadipocyte/group 3 cells from C57BL/6J male mice by qPCR (n = 3 biological replicates (left panel), n = 1 technical replicate collected from 3 biological sources (right panel); unpaired t test, one-tailed; ***p < 0.001, **p < 0.01, *p < 0.05). **(D)** Representative brightfield and fluorescent images (Bodipi staining, green) of differentiating progenitors, immature preadipocytes, and committed/group 3 preadipocytes *in vitro* on day 8 after the start of the differentiation protocol (n = 3 biological replicates from C57BL/6J male mice). **(E)** Quantification of mature adipocyte-specific gene expression by qPCR (n = 3 biological replicates from C57BL/6J male mice; unpaired t test, one-tailed; **p < 0.01, *p < 0.05).

Gene expression of putative markers for in vivo models of lineage tracing

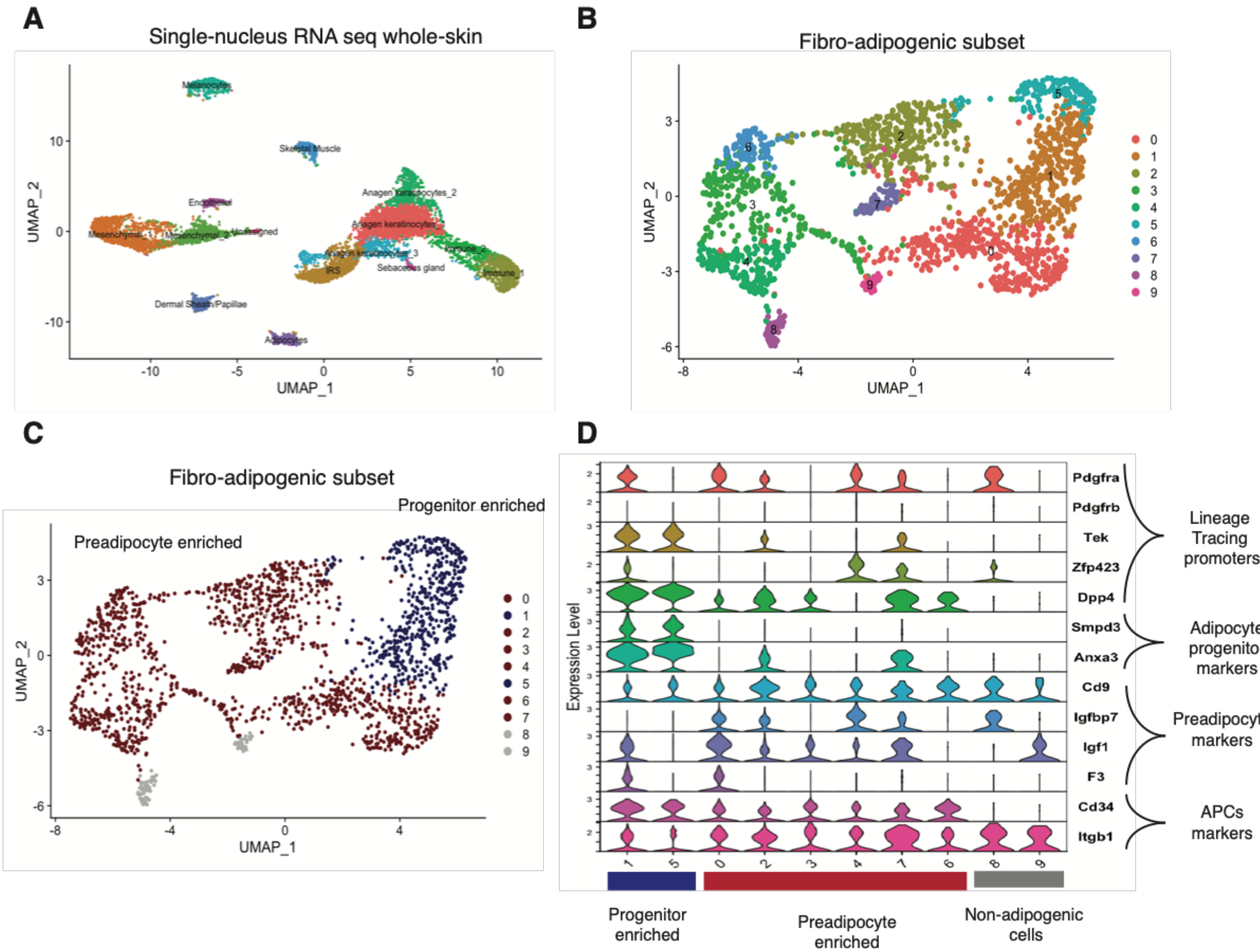

**Fig. S4. Current models for in vivo lineage tracing lack the specificity to track the differentiation of APCs in the skin. (A)** UMAP of cells from whole-skin nuclei isolated from C57BL/6J male P32 mouse tissue. (**B)** UMAP of cells after sub-setting and re-clustering of mesenchymal clusters. (**C)** Manual classification of re-clustered mesenchymal cells into progenitors and preadipocytes based on selected marker gene expression. (**D)** Gene expression comparison of lineage-tracing, adipocyte progenitor, and preadipocyte genes between clusters of progenitors and preadipocytes.

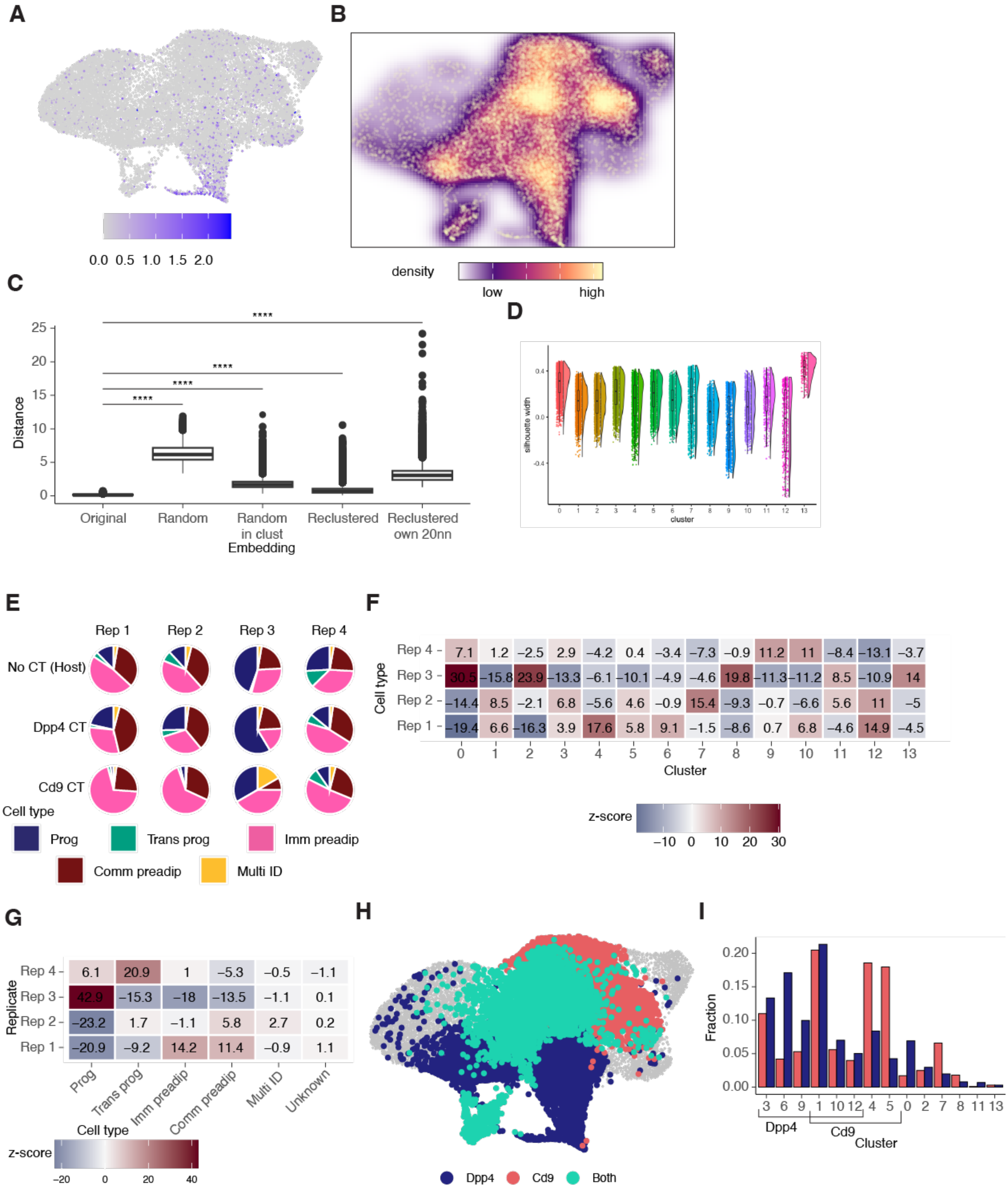
A
0.0 0.5 1.0 1.5 2.0
B
density
low
high
C
Distance
25
20
15
10
5
0
****
****
****
****
Original
Random
Random in clust Embedding
Reclustered
Reclustered own 20nn
D
silhouette width
0.4
0.0
-0.4
0 1 2 3 4 5 6 7 8 9 10 11 12 13
cluster
E
Rep 1
Rep 2
Rep 3
Rep 4
No CT (Host)
Dpp4 CT
Cd9 CT
Cell type
Prog
Trans prog
Imm preadip
Comm preadip
Multi ID
F
Cell type
Rep 4 7.1 1.2 −2.5 2.9 −4.2 0.4 −3.4 −7.3 −0.9 11.2 11 −8.4 −13.1 −3.7
Rep 3 30.5 −15.8 23.9 −13.3 −6.1 −10.1 −4.9 −4.6 19.8 −11.3 −11.2 8.5 −10.9 14
Rep 2 −14.4 8.5 −2.1 6.8 −5.6 4.6 −0.9 15.4 −9.3 −0.7 −6.6 5.6 11 −5
Rep 1 −19.4 6.6 −16.3 3.9 17.6 5.8 9.1 −1.5 −8.6 0.7 6.8 −4.6 14.9 −4.5
0 1 2 3 4 5 6 7 8 9 10 11 12 13
Cluster
z-score
−10 0 10 20 30
G
Replicate
Rep 4 6.1 20.9 1 −5.3 −0.5 −1.1
Rep 3 42.9 −15.3 −18 −13.5 −1.1 0.1
Rep 2 −23.2 1.7 −1.1 5.8 2.7 0.2
Rep 1 −20.9 −9.2 14.2 11.4 −0.9 1.1
Prog
Trans prog
Imm preadip
Comm preadip
Multi ID
Unknown
Cell type
z-score
−20 0 20 40
H
Dpp4
Cd9
Both
I
Fraction
0.20
0.15
0.10
0.05
0.00
3 6 9 1 10 12 4 5 0 2 7 8 11 13
Dpp4
Cd9
Cluster

**Fig. S5. CellTag data enables illustration of divergent potential of distinct preadipocyte populations. (A)** Feature expression of *Cebpa* in P32 adipocyte precursor cells. **(B)** Heatmap of transplanted GFP and/or CellTag expressing cells in the integrated UMAP space. **(C)** Average Euclidean distance of each host cell (GFP-/CellTag-) to, left to right: its 20 nearest neighbors in original embedding, its 20 nearest neighbors in re-embedded data following removal of transplanted cells, 20 random cells in re-embedded data following removal of transplanted cells, 20 random cells in shared Louvain cluster of re-embedded data. **(D)** Silhouette scores of Louvain clusters in P32 adipocyte precursor cells. **(E)** Capybara-assigned cell types using P21 skin data as reference for each biological replicate and CellTagged or host (None) population. **(F) and (G)** Heatmaps of Jaccard similarities for biological replicates and Louvain clusters (F) and Capybara-assigned cell types (G) for P32 adipocyte precursors. **(H)** UMAP of Louvain clusters significantly enriched for one or both CellTag populations using 1,000 randomized permutation tests. **(I)** Fractions of each Louvain cluster comprised of cells with either CellTag population.

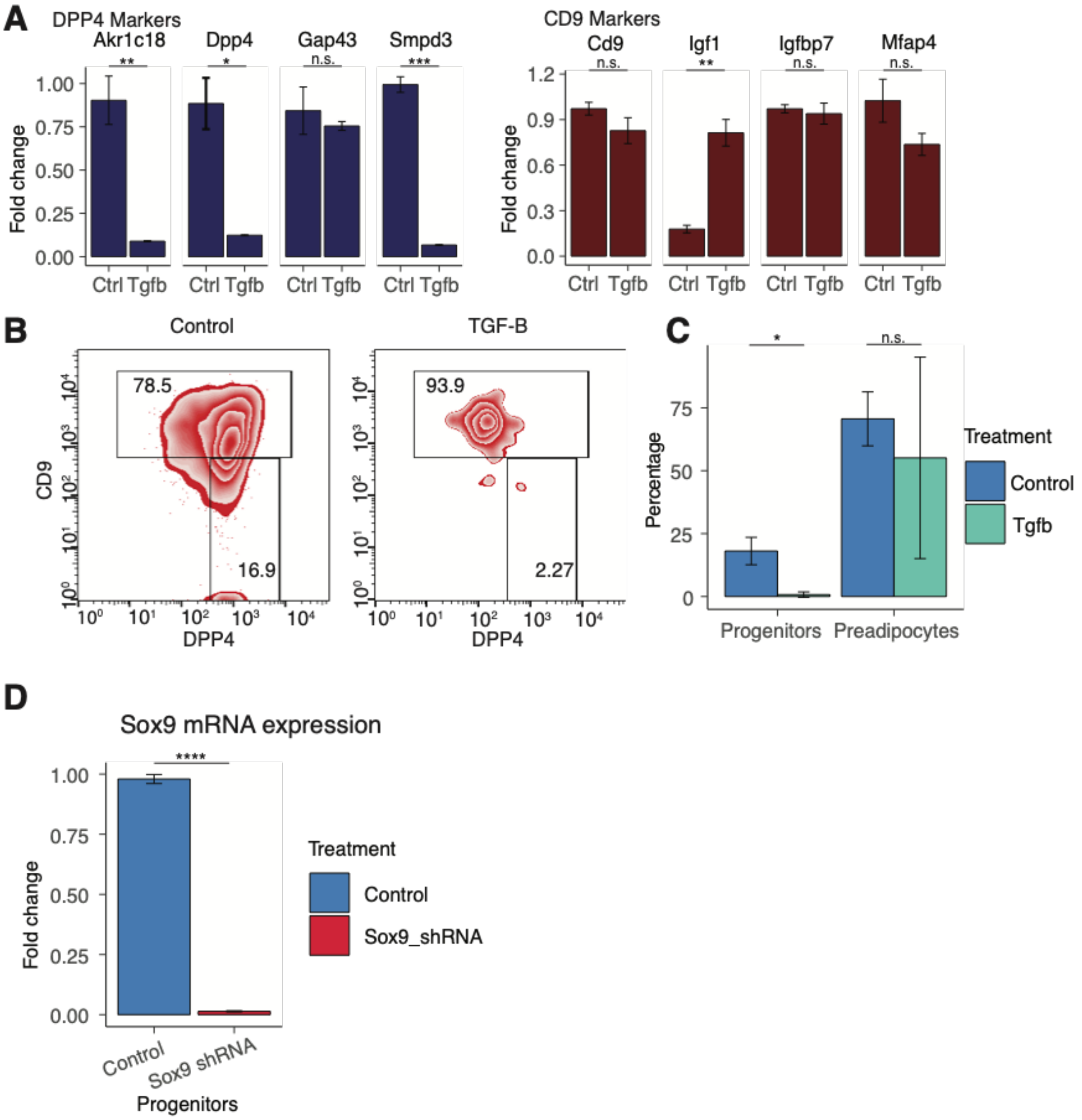
A
DPP4 Markers
Akr1c18
Dpp4
Gap43
Smpd3
**
*
n.s.
***
Fold change
1.00
0.75
0.50
0.25
0.00
Ctrl Tgfb
CD9 Markers
Cd9
Igf1
Igfbp7
Mfap4
n.s.
**
1.2
0.9
0.6
0.3
0.0
B
Control
TGF-B
78.5
16.9
93.9
2.27
CD9
DPP4
C
Percentage
75
50
25
0
Progenitors
Preadipocytes
Treatment
Control
Tgfb
D
Sox9 mRNA expression
****
Fold change
1.00
0.75
0.50
0.25
0.00
Treatment
Control
Sox9_shRNA
Control
Sox9 shRNA
Progenitors

**Fig. S6. TGFB is not sufficient to maintain progenitor identity in the skin. (A)** Relative gene expression of progenitor and preadipocyte marker genes of progenitor cells isolated from P21 C57BL/6J male mice and cultured in the presence of TGFB (10ng/ml) or BSA (10ng/ml, Control) for 5 days (n = 3 biological replicates; unpaired t test with Welch's correction, two-tailed; ***p < 0.001, **p < 0.01, *p < 0.05). **(B) and (C)** Representative FACS plots (B) and quantification (C) of transplanted progenitors treated with BSA (Control) or TGF-B recovered 11-14 days after injection and analyzed for expression of CD9 and DPP4 (n = 3 biological replicates; unpaired t test with Welch's correction, two-tailed; *p < 0.05). **(D)** Relative *Sox9* expression of progenitor cells transduced with a GFP and scramble shRNA (control) or with GFP and *Sox9* shRNA expression cassette after 5-7 days in culture (n = 3 technical replicates; unpaired t test with Welch's correction, two-tailed; ****p < 0.0001).

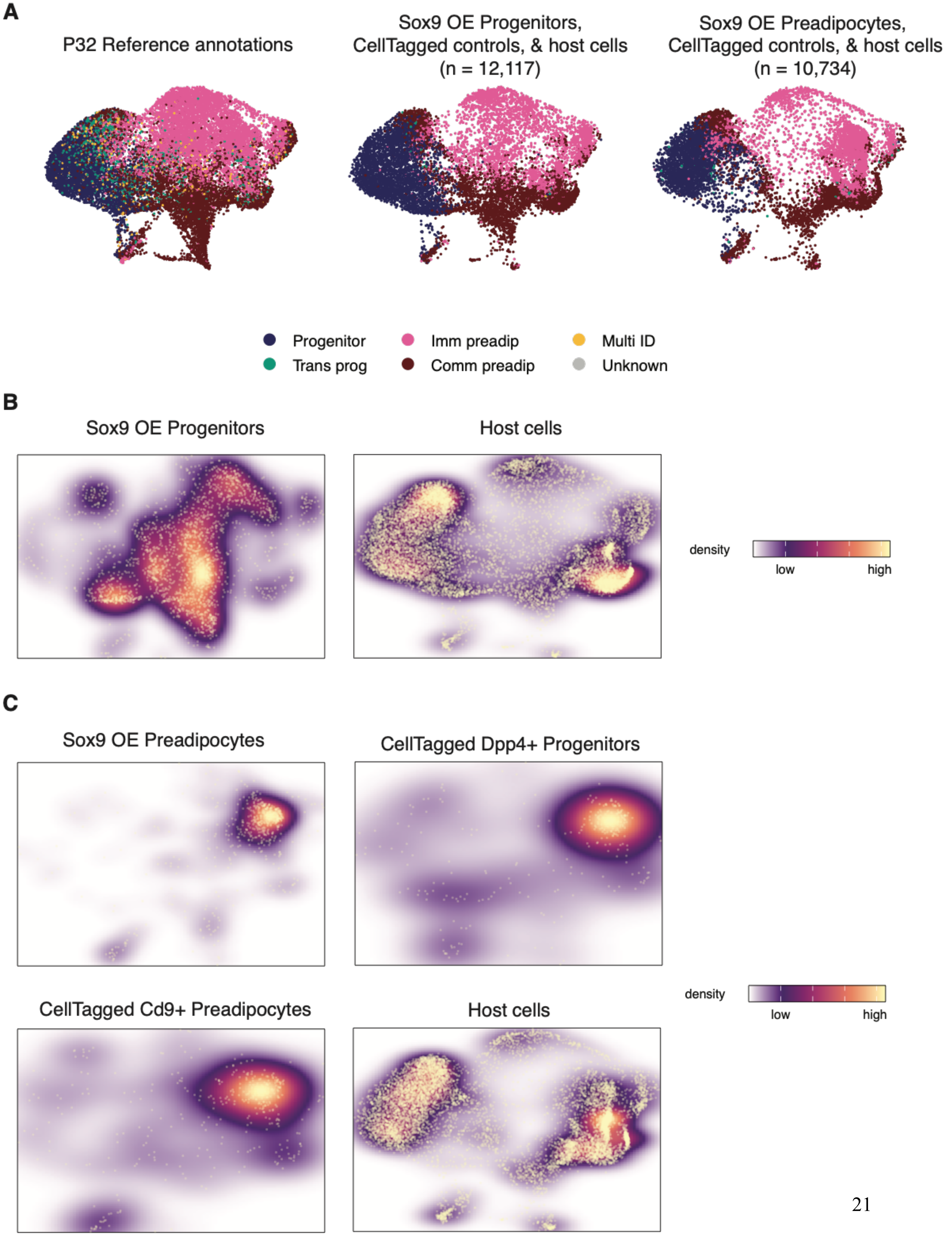

A
P32 Reference annotations
Sox9 OE Progenitors, CellTagged controls, & host cells (n = 12,117)
Sox9 OE Preadipocytes, CellTagged controls, & host cells (n = 10,734)
Progenitor
Trans prog
Imm preadip
Comm preadip
Multi ID
Unknown
B
Sox9 OE Progenitors
Host cells
density
low
high
C
Sox9 OE Preadipocytes
CellTagged Dpp4+ Progenitors
CellTagged Cd9+ Preadipocytes
Host cells
density
low
high

**Fig. S7. Projection of skin precursors with Sox9-overexpressing populations onto P32 homeostatic precursor embedding reveals strong co-clustering mimicking Capybara-identified cell types. (A)** UMAP of P32 skin adipocyte precursors classified using the P21 data as reference (left), *Sox9*-overexpressing progenitors and host cells projected onto the embedding of our P32 data (middle), and projection *Sox9*-overexpressing preadipocytes, controls, and host cells onto the same embedding (right) highlighting the distribution of precursor cell types identified by Capybara with our P21 dataset as reference. **(B)** Heatmaps showing distribution of *Sox9*-overexpressed progenitors (left) and host cells (right) onto P32 homeostatic precursor embedding. **(C)** Heatmaps showing distribution of *Sox9*-overexpressed preadipocytes (top left), control progenitors (top right), control preadipocytes (bottom left), and host cells (bottom right) onto P32 homeostatic precursor embedding.

**Table S1. Differentially-expressed genes for cell types in new classification framework.** Average log fold change between cell type of interest (cluster) and remaining cells in P21 dataset, as well as p-value and bonferroni-corrected p-value. Fraction of cells expressing gene in cell type of interest (pct.1), as well as fraction of cells remaining in dataset expressing the same gene. Differential expression testing is performed using the Wilcoxon Rank Sum Test.

| | p_val | avg_log2FC | pct.1 | pct.2 | p_val_adj | cluster | gene |
|---|---|---|---|---|---|---|---|
| Smpd3 | 0 | 1.95117101 | 0.923 | 0.725 | 0 | Int prog | Smpd3 |
| Akr1c18 | 0 | 1.87930346 | 0.839 | 0.625 | 0 | Int prog | Akr1c18 |
| Pi16 | 0 | 1.81207612 | 0.996 | 0.913 | 0 | Int prog | Pi16 |
| Anxa3 | 0 | 1.68818243 | 0.987 | 0.888 | 0 | Int prog | Anxa3 |
| Sbsn | 0 | 1.67874169 | 0.714 | 0.638 | 0 | Int prog | Sbsn |
| Igfbp5 | 0 | 1.40291419 | 0.964 | 0.845 | 0 | Int prog | Igfbp5 |
| Igfbp4 | 0 | 1.37628388 | 0.927 | 0.749 | 0 | Int prog | Igfbp4 |
| Dpp4 | 0 | 1.34640854 | 0.965 | 0.869 | 0 | Int prog | Dpp4 |
| Cd55 | 0 | 1.23329701 | 0.922 | 0.858 | 0 | Int prog | Cd55 |
| Ackr3 | 0 | 1.18424561 | 0.849 | 0.738 | 0 | Int prog | Ackr3 |
| Ptgs2 | 0 | 1.1441147 | 0.811 | 0.734 | 0 | Int prog | Ptgs2 |
| Wnt2 | 0 | 1.06665408 | 0.774 | 0.681 | 0 | Int prog | Wnt2 |
| Efemp1 | 0 | 0.9964565 | 0.888 | 0.703 | 0 | Int prog | Efemp1 |
| Il1r2 | 0 | 0.99307541 | 0.828 | 0.866 | 0 | Int prog | Il1r2 |
| Krtdap | 0 | 0.97668052 | 0.607 | 0.532 | 0 | Int prog | Krtdap |
| Mustn1 | 0 | 0.96775243 | 0.807 | 0.786 | 0 | Int prog | Mustn1 |
| Il33 | 0 | 0.95530212 | 0.715 | 0.503 | 0 | Int prog | Il33 |
| Prss23 | 0 | 0.95424267 | 0.987 | 0.914 | 0 | Int prog | Prss23 |
| Aldh1a3 | 0 | 0.94583108 | 0.641 | 0.537 | 0 | Int prog | Aldh1a3 |
| C3 | 0 | 0.90839191 | 0.878 | 0.768 | 0 | Int prog | C3 |
| Sfrp2 | 0 | 0.90104535 | 0.983 | 0.818 | 0 | Int prog | Sfrp2 |
| Plac8 | 0 | 0.8151322 | 0.926 | 0.691 | 0 | Int prog | Plac8 |
| Ifi27l2a | 0 | 0.78058294 | 0.99 | 0.932 | 0 | Int prog | Ifi27l2a |
| Lepr | 0 | -0.2921511 | 0.166 | 0.708 | 0 | Int prog | Lepr |
| Nkain4 | 0 | -0.3209841 | 0.424 | 0.751 | 0 | Int prog | Nkain4 |
| Lgals3 | 0 | -0.4659383 | 0.424 | 0.811 | 0 | Int prog | Lgals3 |
| Hmcn1 | 0 | -0.480412 | 0.221 | 0.717 | 0 | Int prog | Hmcn1 |
| Ndrg1 | 0 | -0.4988636 | 0.33 | 0.711 | 0 | Int prog | Ndrg1 |
| Rbp4 | 0 | -0.5030581 | 0.324 | 0.663 | 0 | Int prog | Rbp4 |
| Pltp | 0 | -0.5864527 | 0.22 | 0.618 | 0 | Int prog | Pltp |
| Cav1 | 0 | -0.7013998 | 0.37 | 0.778 | 0 | Int prog | Cav1 |
| Timp1 | 0 | -0.810597 | 0.54 | 0.812 | 0 | Int prog | Timp1 |
| Sdc4 | 0 | -0.8409048 | 0.793 | 0.935 | 0 | Int prog | Sdc4 |
| Thbs1 | 0 | -0.8415122 | 0.405 | 0.764 | 0 | Int prog | Thbs1 |
| Tgfbi | 0 | -0.9747839 | 0.538 | 0.84 | 0 | Int prog | Tgfbi |
| Gadd45g | 0 | -0.98306 | 0.736 | 0.904 | 0 | Int prog | Gadd45g |
| Itgbl1 | 0 | -1.0127078 | 0.549 | 0.826 | 0 | Int prog | Itgbl1 |
| Tnmd | 0 | -1.0279085 | 0.295 | 0.77 | 0 | Int prog | Tnmd |

| | | | | | | | |
|---|---|---|---|---|---|---|---|
| Col4a2 | 0 | -1.0334562 | 0.535 | 0.912 | 0 | Int prog | Col4a2 |
| Emp1 | 0 | -1.0623782 | 0.695 | 0.912 | 0 | Int prog | Emp1 |
| 1500015O10Rik | 0 | -1.0633372 | 0.512 | 0.807 | 0 | Int prog | 1500015O10Rik |
| Icam1 | 0 | -1.083733 | 0.513 | 0.891 | 0 | Int prog | Icam1 |
| Hspa1b | 0 | -1.1149261 | 0.622 | 0.905 | 0 | Int prog | Hspa1b |
| Cd9 | 0 | -1.2351832 | 0.524 | 0.887 | 0 | Int prog | Cd9 |
| Col15a1 | 0 | -1.3067123 | 0.26 | 0.749 | 0 | Int prog | Col15a1 |
| Sfrp1 | 0 | -1.3121097 | 0.483 | 0.779 | 0 | Int prog | Sfrp1 |
| Cxcl14 | 0 | -1.3253704 | 0.481 | 0.792 | 0 | Int prog | Cxcl14 |
| Lox | 0 | -1.3574575 | 0.731 | 0.887 | 0 | Int prog | Lox |
| Col4a1 | 0 | -1.3651885 | 0.65 | 0.939 | 0 | Int prog | Col4a1 |
| Cxcl12 | 0 | -1.3934246 | 0.464 | 0.809 | 0 | Int prog | Cxcl12 |
| C1qtnf3 | 0 | -1.6703305 | 0.26 | 0.675 | 0 | Int prog | C1qtnf3 |
| Igf1 | 0 | -1.7235513 | 0.85 | 0.973 | 0 | Int prog | Igf1 |
| Gas6 | 0 | -1.7321869 | 0.304 | 0.831 | 0 | Int prog | Gas6 |
| Cilp | 0 | -2.0689423 | 0.213 | 0.681 | 0 | Int prog | Cilp |
| Igfbp7 | 0 | -2.4387493 | 0.686 | 0.98 | 0 | Int prog | Igfbp7 |
| Mgp | 0 | -2.5041246 | 0.164 | 0.713 | 0 | Int prog | Mgp |
| Mfap4 | 0 | -3.6052004 | 0.371 | 0.858 | 0 | Int prog | Mfap4 |
| mt-Cytb | 5.37E-303 | -0.4568869 | 0.994 | 0.999 | 7.90E-300 | Int prog | mt-Cytb |
| Pla1a | 5.34E-299 | 0.79172316 | 0.638 | 0.589 | 7.86E-296 | Int prog | Pla1a |
| Ifi205 | 1.64E-290 | 0.6082641 | 0.975 | 0.964 | 2.41E-287 | Int prog | Ifi205 |
| Cst3 | 7.36E-288 | -0.6319864 | 0.974 | 0.987 | 1.08E-284 | Int prog | Cst3 |
| Car3 | 1.51E-284 | -0.4230268 | 0.409 | 0.695 | 2.22E-281 | Int prog | Car3 |
| Abca8a | 8.22E-282 | -0.5244883 | 0.424 | 0.746 | 1.21E-278 | Int prog | Abca8a |
| Fhl1 | 5.02E-281 | 0.55295176 | 0.574 | 0.427 | 7.39E-278 | Int prog | Fhl1 |
| Angptl1 | 1.38E-276 | -0.9934293 | 0.529 | 0.738 | 2.03E-273 | Int prog | Angptl1 |
| Ly6c1 | 4.97E-269 | 0.44882931 | 0.992 | 0.955 | 7.31E-266 | Int prog | Ly6c1 |
| Tpm1 | 2.73E-266 | -0.7152114 | 0.648 | 0.847 | 4.02E-263 | Int prog | Tpm1 |
| F3 | 1.73E-261 | -0.8417303 | 0.395 | 0.699 | 2.55E-258 | Int prog | F3 |
| Gap43 | 1.43E-256 | 0.73450552 | 0.663 | 0.556 | 2.11E-253 | Int prog | Gap43 |
| Peg3 | 1.18E-255 | -0.814316 | 0.571 | 0.816 | 1.74E-252 | Int prog | Peg3 |
| Lpl | 1.58E-255 | -0.9623248 | 0.887 | 0.967 | 2.32E-252 | Int prog | Lpl |
| Mgll | 8.64E-244 | 0.57820438 | 0.774 | 0.714 | 1.27E-240 | Int prog | Mgll |
| Steap4 | 3.36E-241 | -0.3080345 | 0.307 | 0.623 | 4.95E-238 | Int prog | Steap4 |
| Sparcl1 | 1.37E-240 | -0.54151 | 0.292 | 0.613 | 2.02E-237 | Int prog | Sparcl1 |
| Prkg2 | 1.21E-237 | 0.72072785 | 0.629 | 0.605 | 1.77E-234 | Int prog | Prkg2 |
| Serpina3n | 2.83E-227 | -0.426276 | 0.247 | 0.607 | 4.16E-224 | Int prog | Serpina3n |
| Kitl | 1.14E-225 | -0.4075733 | 0.266 | 0.608 | 1.69E-222 | Int prog | Kitl |

| | | | | | | | |
|---|---|---|---|---|---|---|---|
| Clu | 2.86E-211 | -0.3557009 | 0.379 | 0.649 | 4.22E-208 | Int prog | Clu |
| Ccl2 | 2.43E-205 | 1.15063744 | 0.786 | 0.773 | 3.58E-202 | Int prog | Ccl2 |
| Cpxm2 | 5.39E-205 | -0.5271474 | 0.249 | 0.577 | 7.94E-202 | Int prog | Cpxm2 |
| Hspa1a | 6.41E-197 | -0.6200365 | 0.642 | 0.863 | 9.43E-194 | Int prog | Hspa1a |
| Myoc | 4.44E-195 | -0.7605851 | 0.344 | 0.614 | 6.54E-192 | Int prog | Myoc |
| Ptma | 2.07E-189 | -0.3813707 | 0.987 | 0.993 | 3.04E-186 | Int prog | Ptma |
| Nr4a2 | 4.09E-189 | -0.3739063 | 0.41 | 0.697 | 6.03E-186 | Int prog | Nr4a2 |
| Hpgd | 7.95E-189 | -0.7235378 | 0.619 | 0.847 | 1.17E-185 | Int prog | Hpgd |
| Apoe | 9.54E-188 | -1.331133 | 0.309 | 0.615 | 1.40E-184 | Int prog | Apoe |
| Cotl1 | 1.90E-183 | 0.33950659 | 0.433 | 0.288 | 2.80E-180 | Int prog | Cotl1 |
| Pcsk6 | 1.48E-182 | 0.71629858 | 0.63 | 0.62 | 2.18E-179 | Int prog | Pcsk6 |
| Hbb-bt | 3.76E-178 | 0.36782885 | 0.864 | 0.94 | 5.53E-175 | Int prog | Hbb-bt |
| Fxyd6 | 4.67E-178 | -0.8271772 | 0.281 | 0.586 | 6.87E-175 | Int prog | Fxyd6 |
| Ptx3 | 6.60E-178 | 0.86218622 | 0.796 | 0.809 | 9.72E-175 | Int prog | Ptx3 |
| mt-Nd4 | 3.76E-175 | -0.3265166 | 0.993 | 0.998 | 5.54E-172 | Int prog | mt-Nd4 |
| Wisp2 | 1.10E-174 | -0.9389048 | 0.459 | 0.683 | 1.62E-171 | Int prog | Wisp2 |
| Pcsk5 | 1.20E-173 | -0.6836584 | 0.497 | 0.739 | 1.77E-170 | Int prog | Pcsk5 |
| Ctsk | 1.83E-173 | -0.6065847 | 0.868 | 0.924 | 2.70E-170 | Int prog | Ctsk |
| Tspo | 1.97E-167 | -0.5305637 | 0.713 | 0.882 | 2.89E-164 | Int prog | Tspo |
| Slit2 | 4.35E-165 | -0.585599 | 0.282 | 0.589 | 6.40E-162 | Int prog | Slit2 |
| Tmsb4x | 8.55E-164 | 0.35363612 | 0.986 | 0.984 | 1.26E-160 | Int prog | Tmsb4x |
| Fmo2 | 1.18E-163 | -0.7405246 | 0.415 | 0.657 | 1.73E-160 | Int prog | Fmo2 |
| Hbb-bs | 3.58E-163 | 0.3953124 | 0.999 | 0.998 | 5.28E-160 | Int prog | Hbb-bs |
| Gpm6b | 6.64E-163 | -0.5714728 | 0.217 | 0.566 | 9.77E-160 | Int prog | Gpm6b |
| Fabp4 | 6.05E-159 | -0.7745398 | 0.583 | 0.831 | 8.91E-156 | Int prog | Fabp4 |
| Crip2 | 5.31E-152 | -0.4947109 | 0.371 | 0.632 | 7.82E-149 | Int prog | Crip2 |
| Pdgfrl | 1.70E-151 | -0.6248662 | 0.677 | 0.817 | 2.50E-148 | Int prog | Pdgfrl |
| Hba-a2 | 4.61E-151 | 0.38676062 | 0.95 | 0.975 | 6.79E-148 | Int prog | Hba-a2 |
| Arhgdib | 4.82E-149 | -0.5238814 | 0.303 | 0.607 | 7.09E-146 | Int prog | Arhgdib |
| Hmgcs1 | 1.33E-141 | -0.2766631 | 0.685 | 0.887 | 1.96E-138 | Int prog | Hmgcs1 |
| Hba-a1 | 7.98E-139 | 0.36867885 | 0.976 | 0.985 | 1.17E-135 | Int prog | Hba-a1 |
| Dnajb1 | 3.36E-138 | -0.6368926 | 0.778 | 0.922 | 4.95E-135 | Int prog | Dnajb1 |
| Plagl1 | 2.25E-137 | 0.55373558 | 0.675 | 0.613 | 3.31E-134 | Int prog | Plagl1 |
| AW112010 | 1.01E-127 | -0.3617771 | 0.488 | 0.73 | 1.49E-124 | Int prog | AW112010 |
| Lgr5 | 1.30E-127 | -0.4503742 | 0.283 | 0.557 | 1.92E-124 | Int prog | Lgr5 |
| Mif | 2.00E-127 | -0.4892616 | 0.583 | 0.77 | 2.94E-124 | Int prog | Mif |
| Gas1 | 9.07E-125 | -0.6083408 | 0.656 | 0.78 | 1.34E-121 | Int prog | Gas1 |
| Adamts4 | 1.04E-118 | -0.4212275 | 0.554 | 0.744 | 1.53E-115 | Int prog | Adamts4 |
| Ndufa4l2 | 2.66E-116 | -0.9040646 | 0.259 | 0.551 | 3.92E-113 | Int prog | Ndufa4l2 |

| | | | | | | | |
|---|---|---|---|---|---|---|---|
| Rrad | 1.04E-109 | -1.7305391 | 0.385 | 0.589 | 1.53E-106 | Int prog | Rrad |
| Tubb5 | 8.73E-109 | -0.3811537 | 0.786 | 0.898 | 1.28E-105 | Int prog | Tubb5 |
| Fgl2 | 1.66E-107 | -0.4545537 | 0.844 | 0.949 | 2.45E-104 | Int prog | Fgl2 |
| Pmepa1 | 1.83E-105 | -0.5387117 | 0.869 | 0.897 | 2.69E-102 | Int prog | Pmepa1 |
| Osr2 | 2.87E-105 | 0.37544134 | 0.617 | 0.546 | 4.23E-102 | Int prog | Osr2 |
| Ccl11 | 4.20E-101 | 0.28280451 | 0.523 | 0.452 | 6.18E-98 | Int prog | Ccl11 |
| Col12a1 | 2.69E-99 | -0.5234923 | 0.311 | 0.572 | 3.96E-96 | Int prog | Col12a1 |
| Ier3 | 3.75E-99 | -0.5731317 | 0.799 | 0.896 | 5.52E-96 | Int prog | Ier3 |
| Ccl7 | 3.60E-97 | 0.39401612 | 0.87 | 0.862 | 5.30E-94 | Int prog | Ccl7 |
| Eln | 5.50E-95 | -1.6301379 | 0.834 | 0.847 | 8.09E-92 | Int prog | Eln |
| Gas7 | 1.77E-92 | 0.58448077 | 0.722 | 0.781 | 2.61E-89 | Int prog | Gas7 |
| Cxcl1 | 3.39E-92 | 0.64771234 | 0.856 | 0.846 | 4.99E-89 | Int prog | Cxcl1 |
| Thbs2 | 1.36E-90 | -0.3927618 | 0.586 | 0.759 | 2.01E-87 | Int prog | Thbs2 |
| Gpx1 | 5.45E-89 | -0.3378756 | 0.87 | 0.956 | 8.02E-86 | Int prog | Gpx1 |
| Has1 | 6.68E-87 | 0.39791329 | 0.709 | 0.671 | 9.83E-84 | Int prog | Has1 |
| Bmp2 | 1.96E-84 | 0.26286602 | 0.506 | 0.553 | 2.89E-81 | Int prog | Bmp2 |
| Irf7 | 3.18E-83 | 0.37587555 | 0.643 | 0.666 | 4.69E-80 | Int prog | Irf7 |
| Tuba1b | 5.36E-80 | -0.2605868 | 0.662 | 0.875 | 7.89E-77 | Int prog | Tuba1b |
| Cited2 | 2.74E-79 | 0.38607069 | 0.474 | 0.452 | 4.03E-76 | Int prog | Cited2 |
| Vcam1 | 1.72E-76 | -0.4053158 | 0.574 | 0.742 | 2.54E-73 | Int prog | Vcam1 |
| Ldha | 4.02E-73 | -0.3360648 | 0.746 | 0.851 | 5.92E-70 | Int prog | Ldha |
| Socs3 | 3.57E-68 | -0.4546062 | 0.647 | 0.794 | 5.25E-65 | Int prog | Socs3 |
| Tnfsf9 | 1.52E-67 | 0.30044457 | 0.683 | 0.759 | 2.24E-64 | Int prog | Tnfsf9 |
| Ddah1 | 2.47E-67 | -0.3317421 | 0.341 | 0.567 | 3.64E-64 | Int prog | Ddah1 |
| Grem2 | 9.82E-66 | -0.4067722 | 0.341 | 0.541 | 1.45E-62 | Int prog | Grem2 |
| Edn1 | 1.11E-65 | 0.51510896 | 0.533 | 0.646 | 1.64E-62 | Int prog | Edn1 |
| Igfbp3 | 8.66E-64 | -0.5675049 | 0.444 | 0.655 | 1.27E-60 | Int prog | Igfbp3 |
| Mafb | 2.25E-58 | -0.7977121 | 0.444 | 0.604 | 3.31E-55 | Int prog | Mafb |
| Cyr61 | 1.75E-57 | -0.3676024 | 0.525 | 0.708 | 2.58E-54 | Int prog | Cyr61 |
| Tgm2 | 1.85E-57 | -0.4764374 | 0.362 | 0.556 | 2.72E-54 | Int prog | Tgm2 |
| Ccl8 | 2.75E-57 | -0.553221 | 0.458 | 0.64 | 4.05E-54 | Int prog | Ccl8 |
| Aspn | 3.01E-57 | -0.3788754 | 0.954 | 0.95 | 4.43E-54 | Int prog | Aspn |
| Clec11a | 1.74E-56 | -0.4075382 | 0.416 | 0.601 | 2.56E-53 | Int prog | Clec11a |
| Bst2 | 9.39E-56 | 0.28915037 | 0.835 | 0.798 | 1.38E-52 | Int prog | Bst2 |
| Rnf19b | 1.35E-55 | -0.3822638 | 0.262 | 0.528 | 1.99E-52 | Int prog | Rnf19b |
| Mgst3 | 1.39E-54 | -0.2993616 | 0.221 | 0.503 | 2.05E-51 | Int prog | Mgst3 |
| Gem | 1.49E-54 | -0.5070368 | 0.66 | 0.755 | 2.19E-51 | Int prog | Gem |
| Tmem176a | 2.08E-54 | -0.2787699 | 0.126 | 0.431 | 3.06E-51 | Int prog | Tmem176a |
| Hmcn2 | 1.26E-53 | -0.6556655 | 0.107 | 0.472 | 1.85E-50 | Int prog | Hmcn2 |

| | | | | | | | |
|---|---|---|---|---|---|---|---|
| Rgs16 | 3.15E-52 | -0.3500433 | 0.168 | 0.291 | 4.64E-49 | Int prog | Rgs16 |
| Pappa2 | 8.19E-52 | -0.4171229 | 0.128 | 0.469 | 1.21E-48 | Int prog | Pappa2 |
| Fxyd5 | 3.74E-48 | -0.3915114 | 0.462 | 0.632 | 5.50E-45 | Int prog | Fxyd5 |
| Phlda1 | 7.07E-45 | -0.4740725 | 0.67 | 0.792 | 1.04E-41 | Int prog | Phlda1 |
| Crispld2 | 1.18E-43 | -0.3778227 | 0.453 | 0.571 | 1.74E-40 | Int prog | Crispld2 |
| Dmkn | 2.52E-42 | 0.3440151 | 0.464 | 0.537 | 3.71E-39 | Int prog | Dmkn |
| Cdkn1c | 9.13E-42 | -0.4164212 | 0.719 | 0.809 | 1.34E-38 | Int prog | Cdkn1c |
| Gadd45b | 1.96E-40 | -0.3512723 | 0.728 | 0.807 | 2.89E-37 | Int prog | Gadd45b |
| Id2 | 2.70E-40 | -0.4243072 | 0.408 | 0.598 | 3.97E-37 | Int prog | Id2 |
| Sepp1 | 1.81E-39 | -0.261692 | 0.946 | 0.98 | 2.66E-36 | Int prog | Sepp1 |
| Dbi | 3.03E-38 | -0.3672795 | 0.881 | 0.944 | 4.46E-35 | Int prog | Dbi |
| Sema3e | 9.64E-37 | 0.25186773 | 0.411 | 0.46 | 1.42E-33 | Int prog | Sema3e |
| C4b | 6.60E-36 | -0.4284836 | 0.533 | 0.68 | 9.72E-33 | Int prog | C4b |
| Fbn2 | 2.28E-34 | -0.4497006 | 0.501 | 0.637 | 3.35E-31 | Int prog | Fbn2 |
| Gbp2 | 5.25E-34 | -0.3278698 | 0.532 | 0.688 | 7.73E-31 | Int prog | Gbp2 |
| Tmem176b | 1.38E-33 | -0.6031654 | 0.283 | 0.501 | 2.04E-30 | Int prog | Tmem176b |
| Mdk | 7.33E-33 | -0.5411237 | 0.274 | 0.5 | 1.08E-29 | Int prog | Mdk |
| Ccnd1 | 8.43E-33 | -0.2553824 | 0.392 | 0.592 | 1.24E-29 | Int prog | Ccnd1 |
| Matn2 | 2.52E-31 | -0.3676953 | 0.335 | 0.552 | 3.70E-28 | Int prog | Matn2 |
| Csrp1 | 5.46E-30 | -0.3476818 | 0.324 | 0.518 | 8.04E-27 | Int prog | Csrp1 |
| Fmod | 6.10E-30 | -0.4915358 | 0.127 | 0.44 | 8.99E-27 | Int prog | Fmod |
| CellTag.UTR | 4.83E-28 | -0.3991167 | 0.232 | 0.508 | 7.11E-25 | Int prog | CellTag.UTR |
| Iigp1 | 1.76E-26 | -0.4137538 | 0.456 | 0.603 | 2.60E-23 | Int prog | Iigp1 |
| Sned1 | 1.42E-22 | -0.2660763 | 0.452 | 0.621 | 2.09E-19 | Int prog | Sned1 |
| Itpkc | 1.97E-22 | -0.2852653 | 0.327 | 0.51 | 2.90E-19 | Int prog | Itpkc |
| Mmp3 | 5.89E-22 | -0.3040367 | 0.611 | 0.742 | 8.67E-19 | Int prog | Mmp3 |
| Vcan | 1.24E-21 | -0.2757492 | 0.765 | 0.829 | 1.83E-18 | Int prog | Vcan |
| Ptn | 1.83E-21 | -0.7120949 | 0.439 | 0.573 | 2.70E-18 | Int prog | Ptn |
| Dlk1 | 6.87E-20 | -0.5167883 | 0.239 | 0.488 | 1.01E-16 | Int prog | Dlk1 |
| Adamtsl2 | 1.75E-19 | -0.3381727 | 0.073 | 0.412 | 2.57E-16 | Int prog | Adamtsl2 |
| Nov | 4.10E-18 | -0.2777655 | 0.39 | 0.437 | 6.04E-15 | Int prog | Nov |
| Col8a1 | 8.76E-18 | -0.6783301 | 0.099 | 0.422 | 1.29E-14 | Int prog | Col8a1 |
| Nr4a3 | 4.07E-17 | 0.32031969 | 0.604 | 0.697 | 5.99E-14 | Int prog | Nr4a3 |
| Tubb4b | 5.27E-16 | -0.272168 | 0.461 | 0.604 | 7.76E-13 | Int prog | Tubb4b |
| Il1rl1 | 6.04E-16 | 0.37091104 | 0.533 | 0.686 | 8.89E-13 | Int prog | Il1rl1 |
| Serpine1 | 1.64E-14 | -0.4809451 | 0.326 | 0.49 | 2.42E-11 | Int prog | Serpine1 |
| Socs1 | 2.91E-13 | -0.3352938 | 0.342 | 0.51 | 4.28E-10 | Int prog | Socs1 |
| Runx1 | 1.13E-12 | -0.2561765 | 0.295 | 0.471 | 1.67E-09 | Int prog | Runx1 |
| Plvap | 2.78E-12 | -0.2689119 | 0.264 | 0.466 | 4.09E-09 | Int prog | Plvap |

| | | | | | | | |
|---|---|---|---|---|---|---|---|
| Gpx3 | 1.04E-11 | -0.3972813 | 0.987 | 0.974 | 1.53E-08 | Int prog | Gpx3 |
| Mylk | 1.11E-11 | -0.3171966 | 0.299 | 0.475 | 1.63E-08 | Int prog | Mylk |
| Meox1 | 1.33E-10 | -0.2691373 | 0.115 | 0.334 | 1.95E-07 | Int prog | Meox1 |
| Ctla2a | 6.99E-10 | -0.5441933 | 0.281 | 0.401 | 1.03E-06 | Int prog | Ctla2a |
| Ncam1 | 8.09E-08 | -0.3877835 | 0.235 | 0.422 | 0.00011904 | Int prog | Ncam1 |
| Cxcl2 | 8.69E-08 | 0.51303375 | 0.6 | 0.712 | 0.00012788 | Int prog | Cxcl2 |
| Cxcl9 | 1.48E-06 | -0.8129508 | 0.189 | 0.356 | 0.00217653 | Int prog | Cxcl9 |
| GFP.CDS | 1.05E-05 | -0.3815611 | 0.237 | 0.449 | 0.01539209 | Int prog | GFP.CDS |
| Emp2 | 1.98E-05 | -0.3228668 | 0.267 | 0.443 | 0.02913338 | Int prog | Emp2 |
| Wisp1 | 4.08E-05 | -0.2762315 | 0.16 | 0.361 | 0.06009326 | Int prog | Wisp1 |
| Epha4 | 4.37E-05 | -0.3402833 | 0.222 | 0.437 | 0.0642734 | Int prog | Epha4 |
| Fbln7 | 9.18E-05 | -0.4890292 | 0.141 | 0.424 | 0.13509415 | Int prog | Fbln7 |
| Gdf10 | 0.00012849 | -0.2514019 | 0.175 | 0.361 | 0.18913874 | Int prog | Gdf10 |
| Gfra1 | 0.00178027 | -0.289532 | 0.21 | 0.432 | 1 | Int prog | Gfra1 |
| Smpd3.1 | 1.23E-60 | 0.31114293 | 0.898 | 0.764 | 1.80E-57 | Trans int prog | Smpd3 |
| Pi16.1 | 2.01E-54 | 0.41376344 | 0.982 | 0.929 | 2.96E-51 | Trans int prog | Pi16 |
| Dpp4.1 | 2.33E-54 | 0.34551034 | 0.966 | 0.887 | 3.43E-51 | Trans int prog | Dpp4 |
| Igfbp7.1 | 3.69E-45 | -0.8060859 | 0.956 | 0.908 | 5.43E-42 | Trans int prog | Igfbp7 |
| Anxa3.1 | 7.09E-42 | 0.30754593 | 0.979 | 0.907 | 1.04E-38 | Trans int prog | Anxa3 |
| Ifi205.1 | 1.87E-39 | 0.44859705 | 0.995 | 0.964 | 2.75E-36 | Trans int prog | Ifi205 |
| Postn | 5.42E-39 | 0.40950168 | 0.987 | 0.957 | 7.97E-36 | Trans int prog | Postn |
| Rbp4.1 | 1.43E-37 | -0.3109817 | 0.547 | 0.586 | 2.11E-34 | Trans int prog | Rbp4 |
| Cpxm2.1 | 2.52E-37 | -0.3677555 | 0.461 | 0.502 | 3.71E-34 | Trans int prog | Cpxm2 |
| Prss23.1 | 3.93E-37 | 0.37399484 | 0.979 | 0.929 | 5.79E-34 | Trans int prog | Prss23 |
| Hspb1 | 9.05E-33 | -0.5393191 | 0.659 | 0.701 | 1.33E-29 | Trans int prog | Hspb1 |
| Mgp.1 | 1.96E-32 | -1.0998258 | 0.563 | 0.585 | 2.88E-29 | Trans int prog | Mgp |
| Inmt | 4.44E-31 | 0.27001395 | 0.758 | 0.632 | 6.54E-28 | Trans int prog | Inmt |
| Ptma.1 | 5.17E-30 | -0.2845933 | 0.994 | 0.992 | 7.61E-27 | Trans int prog | Ptma |
| Fmod.1 | 1.42E-26 | -0.3527236 | 0.262 | 0.373 | 2.09E-23 | Trans int prog | Fmod |
| Sparcl1.1 | 1.73E-26 | -0.3339194 | 0.464 | 0.542 | 2.55E-23 | Trans int prog | Sparcl1 |
| Cst3.1 | 3.06E-26 | -0.363848 | 0.979 | 0.984 | 4.51E-23 | Trans int prog | Cst3 |
| mt-Cytb.1 | 1.74E-25 | -0.267939 | 0.999 | 0.997 | 2.56E-22 | Trans int prog | mt-Cytb |
| Gap43.1 | 2.64E-25 | 0.31789768 | 0.736 | 0.572 | 3.88E-22 | Trans int prog | Gap43 |
| Sfrp2.1 | 2.68E-25 | 0.33498367 | 0.907 | 0.854 | 3.95E-22 | Trans int prog | Sfrp2 |
| AY036118 | 1.93E-24 | -1.2416977 | 0.865 | 0.824 | 2.85E-21 | Trans int prog | AY036118 |
| Gpx3.1 | 2.49E-24 | 0.25209494 | 0.991 | 0.976 | 3.67E-21 | Trans int prog | Gpx3 |
| Cd9.1 | 4.61E-23 | -0.4555827 | 0.801 | 0.801 | 6.79E-20 | Trans int prog | Cd9 |
| Gpm6b.1 | 5.78E-22 | -0.2920339 | 0.434 | 0.487 | 8.51E-19 | Trans int prog | Gpm6b |
| Has1.1 | 1.46E-20 | 0.33665453 | 0.791 | 0.673 | 2.15E-17 | Trans int prog | Has1 |

| | | | | | | | |
|---|---|---|---|---|---|---|---|
| Dbi.1 | 2.82E-20 | -0.4397966 | 0.946 | 0.928 | 4.15E-17 | Trans int prog | Dbi |
| Cav1.1 | 1.20E-19 | -0.3139793 | 0.626 | 0.685 | 1.76E-16 | Trans int prog | Cav1 |
| Plac8.1 | 1.48E-19 | 0.26809841 | 0.849 | 0.74 | 2.18E-16 | Trans int prog | Plac8 |
| Igf1.1 | 1.56E-19 | -0.4810838 | 0.967 | 0.942 | 2.29E-16 | Trans int prog | Igf1 |
| Apoe.1 | 5.91E-19 | -0.9108334 | 0.517 | 0.544 | 8.69E-16 | Trans int prog | Apoe |
| Ccl2.1 | 1.15E-18 | 0.38297358 | 0.865 | 0.771 | 1.69E-15 | Trans int prog | Ccl2 |
| Rrad.1 | 6.73E-17 | -0.6491907 | 0.524 | 0.542 | 9.91E-14 | Trans int prog | Rrad |
| Ptgs2.1 | 2.12E-16 | 0.27600877 | 0.86 | 0.746 | 3.12E-13 | Trans int prog | Ptgs2 |
| Ccl7.1 | 5.98E-15 | 0.34504882 | 0.942 | 0.859 | 8.80E-12 | Trans int prog | Ccl7 |
| C1qtnf3.1 | 1.78E-14 | -0.6714975 | 0.6 | 0.576 | 2.62E-11 | Trans int prog | C1qtnf3 |
| Col8a1.1 | 7.21E-14 | -0.2857057 | 0.321 | 0.347 | 1.06E-10 | Trans int prog | Col8a1 |
| Ndufa4l2.1 | 9.75E-14 | -0.2793501 | 0.467 | 0.483 | 1.43E-10 | Trans int prog | Ndufa4l2 |
| Tspo.1 | 2.81E-13 | -0.3196669 | 0.857 | 0.841 | 4.13E-10 | Trans int prog | Tspo |
| Col15a1.1 | 8.89E-13 | -0.3299985 | 0.629 | 0.634 | 1.31E-09 | Trans int prog | Col15a1 |
| Mdk.1 | 3.26E-12 | -0.2875579 | 0.432 | 0.448 | 4.80E-09 | Trans int prog | Mdk |
| Car3.1 | 1.09E-10 | -0.3543922 | 0.671 | 0.625 | 1.61E-07 | Trans int prog | Car3 |
| Hba-a2.1 | 2.14E-10 | -0.9689779 | 0.993 | 0.967 | 3.15E-07 | Trans int prog | Hba-a2 |
| Thbs4 | 2.16E-10 | -0.43858 | 0.207 | 0.31 | 3.18E-07 | Trans int prog | Thbs4 |
| Ptn.1 | 2.39E-10 | -0.4745173 | 0.542 | 0.541 | 3.52E-07 | Trans int prog | Ptn |
| Hspa1b.1 | 3.16E-10 | -0.4284939 | 0.852 | 0.838 | 4.65E-07 | Trans int prog | Hspa1b |
| Cilp.1 | 9.50E-10 | -0.7537852 | 0.637 | 0.567 | 1.40E-06 | Trans int prog | Cilp |
| Gadd45b.1 | 2.23E-09 | -0.279054 | 0.803 | 0.787 | 3.29E-06 | Trans int prog | Gadd45b |
| Cxcl2.1 | 3.96E-09 | 0.39410228 | 0.787 | 0.679 | 5.82E-06 | Trans int prog | Cxcl2 |
| Cxcl12.1 | 5.99E-09 | -0.4279813 | 0.729 | 0.727 | 8.81E-06 | Trans int prog | Cxcl12 |
| Tmem176b.1 | 2.01E-08 | -0.3946437 | 0.491 | 0.447 | 2.96E-05 | Trans int prog | Tmem176b |
| Mfap4.1 | 1.19E-07 | -1.1178657 | 0.844 | 0.737 | 0.00017559 | Trans int prog | Mfap4 |
| Prlr | 1.56E-07 | -0.27002 | 0.366 | 0.336 | 0.00023035 | Trans int prog | Prlr |
| Fxyd6.1 | 6.54E-07 | -0.4167396 | 0.572 | 0.511 | 0.00096274 | Trans int prog | Fxyd6 |
| Klf2 | 1.51E-06 | -0.3537954 | 0.827 | 0.8 | 0.00222634 | Trans int prog | Klf2 |
| Tpm1.1 | 1.65E-06 | -0.2732674 | 0.849 | 0.798 | 0.00242925 | Trans int prog | Tpm1 |
| Isg15 | 1.94E-06 | -0.3672644 | 0.763 | 0.723 | 0.002858 | Trans int prog | Isg15 |
| Sdc4.1 | 2.03E-06 | -0.2516957 | 0.969 | 0.898 | 0.00298818 | Trans int prog | Sdc4 |
| Hspd1 | 8.33E-06 | -0.2686894 | 0.856 | 0.81 | 0.01225652 | Trans int prog | Hspd1 |
| Hba-a1.1 | 2.14E-05 | -1.0689096 | 0.991 | 0.982 | 0.03147762 | Trans int prog | Hba-a1 |
| Hspa1a.1 | 3.74E-05 | -0.3060296 | 0.836 | 0.809 | 0.0549886 | Trans int prog | Hspa1a |
| Hbb-bt.1 | 4.52E-05 | -0.8038387 | 0.978 | 0.919 | 0.06659855 | Trans int prog | Hbb-bt |
| Igfbp5.1 | 0.00022662 | -0.3608882 | 0.909 | 0.871 | 0.33358444 | Trans int prog | Igfbp5 |
| Col4a1.1 | 0.00037397 | -0.4262471 | 0.961 | 0.865 | 0.55048454 | Trans int prog | Col4a1 |
| Sfrp1.1 | 0.00048716 | -0.3267633 | 0.772 | 0.706 | 0.71709227 | Trans int prog | Sfrp1 |

| | | | | | | | |
|---|---|---|---|---|---|---|---|
| Hbb-bs.1 | 0.00058256 | -1.0723489 | 0.996 | 0.998 | 0.85752393 | Trans int prog | Hbb-bs |
| Gadd45g.1 | 0.00088065 | -0.3173464 | 0.888 | 0.863 | 1 | Trans int prog | Gadd45g |
| Gas6.1 | 0.00193456 | -0.2568314 | 0.799 | 0.701 | 1 | Trans int prog | Gas6 |
| Col4a2.1 | 0.00253007 | -0.3348376 | 0.957 | 0.815 | 1 | Trans int prog | Col4a2 |
| Fabp4.1 | 0.00261713 | -0.737483 | 0.927 | 0.763 | 1 | Trans int prog | Fabp4 |
| Sbsn.1 | 0.00284939 | -0.3359721 | 0.744 | 0.651 | 1 | Trans int prog | Sbsn |
| Gem.1 | 0.00288323 | -0.2551602 | 0.791 | 0.729 | 1 | Trans int prog | Gem |
| 1500015O10Rik.1 | 0.00305618 | -0.3048097 | 0.796 | 0.734 | 1 | Trans int prog | 1500015O10Rik |
| Itgbl1.1 | 0.00606212 | -0.2686445 | 0.795 | 0.759 | 1 | Trans int prog | Itgbl1 |
| Mfap4.2 | 0 | 2.93759394 | 0.968 | 0.595 | 0 | Imm preadip | Mfap4 |
| Eln.1 | 0 | 2.5604756 | 0.947 | 0.776 | 0 | Imm preadip | Eln |
| Cilp.2 | 0 | 1.81637493 | 0.832 | 0.399 | 0 | Imm preadip | Cilp |
| C1qtnf3.2 | 0 | 1.76554247 | 0.803 | 0.428 | 0 | Imm preadip | C1qtnf3 |
| Lox.1 | 0 | 1.67972566 | 0.964 | 0.775 | 0 | Imm preadip | Lox |
| Itgbl1.2 | 0 | 1.29731268 | 0.903 | 0.667 | 0 | Imm preadip | Itgbl1 |
| Wisp2.1 | 0 | 1.28245985 | 0.771 | 0.537 | 0 | Imm preadip | Wisp2 |
| Mgp.2 | 0 | 1.124641 | 0.815 | 0.431 | 0 | Imm preadip | Mgp |
| Tgfbi.1 | 0 | 0.99523714 | 0.922 | 0.668 | 0 | Imm preadip | Tgfbi |
| Angptl1.1 | 0 | 0.97891446 | 0.826 | 0.599 | 0 | Imm preadip | Angptl1 |
| Gas6.2 | 0 | 0.95486182 | 0.897 | 0.581 | 0 | Imm preadip | Gas6 |
| Pdgfrl.1 | 0 | 0.95147143 | 0.894 | 0.712 | 0 | Imm preadip | Pdgfrl |
| Pmepa1.1 | 0 | 0.94885013 | 0.943 | 0.856 | 0 | Imm preadip | Pmepa1 |
| Igf1.2 | 0 | 0.94300816 | 0.983 | 0.918 | 0 | Imm preadip | Igf1 |
| Pcsk5.1 | 0 | 0.86788386 | 0.792 | 0.609 | 0 | Imm preadip | Pcsk5 |
| Tnmd.1 | 0 | 0.8520092 | 0.838 | 0.54 | 0 | Imm preadip | Tnmd |
| Fbn2.1 | 0 | 0.82944427 | 0.756 | 0.506 | 0 | Imm preadip | Fbn2 |
| Aspn.1 | 0 | 0.82822251 | 0.983 | 0.929 | 0 | Imm preadip | Aspn |
| Emp1.1 | 0 | 0.76774886 | 0.946 | 0.805 | 0 | Imm preadip | Emp1 |
| F3.1 | 0 | 0.76509102 | 0.772 | 0.532 | 0 | Imm preadip | F3 |
| Hpgd.1 | 0 | 0.67853251 | 0.898 | 0.724 | 0 | Imm preadip | Hpgd |
| Cd9.2 | 0 | 0.67658302 | 0.923 | 0.721 | 0 | Imm preadip | Cd9 |
| Cpxm2.2 | 0 | 0.64870991 | 0.65 | 0.4 | 0 | Imm preadip | Cpxm2 |
| Lgr5.1 | 0 | 0.62750036 | 0.629 | 0.402 | 0 | Imm preadip | Lgr5 |
| Hmcn1.1 | 0 | 0.59064606 | 0.776 | 0.485 | 0 | Imm preadip | Hmcn1 |
| Nkain4.1 | 0 | 0.56281516 | 0.813 | 0.582 | 0 | Imm preadip | Nkain4 |
| Cav1.2 | 0 | 0.53712299 | 0.808 | 0.598 | 0 | Imm preadip | Cav1 |
| Ramp1 | 0 | 0.50008479 | 0.739 | 0.467 | 0 | Imm preadip | Ramp1 |
| Clu.1 | 0 | 0.49165391 | 0.743 | 0.482 | 0 | Imm preadip | Clu |
| Nr4a2.1 | 0 | 0.47044143 | 0.741 | 0.557 | 0 | Imm preadip | Nr4a2 |

| | | | | | | | |
|---|---|---|---|---|---|---|---|
| Fgf9 | 0 | 0.28779112 | 0.688 | 0.406 | 0 | Imm preadip | Fgf9 |
| Wnt2.1 | 0 | -0.6304013 | 0.614 | 0.761 | 0 | Imm preadip | Wnt2 |
| Ifi205.2 | 0 | -0.7096807 | 0.954 | 0.974 | 0 | Imm preadip | Ifi205 |
| Il1r2.1 | 0 | -0.7361154 | 0.85 | 0.862 | 0 | Imm preadip | Il1r2 |
| Ifi27l2a.1 | 0 | -0.9329942 | 0.935 | 0.953 | 0 | Imm preadip | Ifi27l2a |
| Plac8.2 | 0 | -0.9358065 | 0.627 | 0.825 | 0 | Imm preadip | Plac8 |
| Efemp1.1 | 0 | -1.0821855 | 0.651 | 0.81 | 0 | Imm preadip | Efemp1 |
| Igfbp5.2 | 0 | -1.1039994 | 0.825 | 0.905 | 0 | Imm preadip | Igfbp5 |
| Igfbp4.1 | 0 | -1.3283269 | 0.716 | 0.84 | 0 | Imm preadip | Igfbp4 |
| Akr1c18.1 | 0 | -1.339977 | 0.586 | 0.735 | 0 | Imm preadip | Akr1c18 |
| Ptx3.1 | 0 | -1.4042839 | 0.758 | 0.837 | 0 | Imm preadip | Ptx3 |
| Smpd3.2 | 0 | -1.5622051 | 0.693 | 0.824 | 0 | Imm preadip | Smpd3 |
| C3.1 | 0 | -1.5996575 | 0.724 | 0.84 | 0 | Imm preadip | C3 |
| Tnfsf9.1 | 1.27E-304 | -0.4908523 | 0.723 | 0.753 | 1.86E-301 | Imm preadip | Tnfsf9 |
| Ndufa4l2.2 | 2.02E-301 | 0.75577401 | 0.646 | 0.374 | 2.97E-298 | Imm preadip | Ndufa4l2 |
| Tnfaip6 | 5.81E-295 | -0.8040632 | 0.92 | 0.941 | 8.55E-292 | Imm preadip | Tnfaip6 |
| Mmp3.1 | 5.66E-293 | -0.9304149 | 0.649 | 0.752 | 8.33E-290 | Imm preadip | Mmp3 |
| Sfrp1.2 | 5.08E-290 | 0.62113124 | 0.815 | 0.64 | 7.48E-287 | Imm preadip | Sfrp1 |
| Fmo2.1 | 3.70E-284 | 0.96837614 | 0.695 | 0.538 | 5.44E-281 | Imm preadip | Fmo2 |
| Col1a1 | 1.21E-282 | 0.51459771 | 1 | 0.999 | 1.78E-279 | Imm preadip | Col1a1 |
| Cxcl13 | 1.93E-279 | -0.578204 | 0.696 | 0.739 | 2.84E-276 | Imm preadip | Cxcl13 |
| Fxyd6.2 | 1.93E-275 | 0.81838915 | 0.641 | 0.43 | 2.84E-272 | Imm preadip | Fxyd6 |
| Arhgdib.1 | 3.79E-265 | 0.49815321 | 0.681 | 0.439 | 5.58E-262 | Imm preadip | Arhgdib |
| Icam1.1 | 9.56E-261 | 0.40379107 | 0.896 | 0.74 | 1.41E-257 | Imm preadip | Icam1 |
| Hspa1b.2 | 4.13E-258 | 0.50911243 | 0.917 | 0.787 | 6.08E-255 | Imm preadip | Hspa1b |
| Rbp4.2 | 2.14E-253 | 0.44678123 | 0.687 | 0.515 | 3.15E-250 | Imm preadip | Rbp4 |
| Cxcl14.1 | 8.72E-251 | 0.62301353 | 0.792 | 0.67 | 1.28E-247 | Imm preadip | Cxcl14 |
| Ccl11.1 | 4.75E-246 | -0.5673638 | 0.341 | 0.553 | 6.99E-243 | Imm preadip | Ccl11 |
| Ndrg1.1 | 4.22E-238 | 0.33447707 | 0.756 | 0.532 | 6.21E-235 | Imm preadip | Ndrg1 |
| Mgst1 | 3.66E-233 | -0.4788954 | 0.886 | 0.921 | 5.39E-230 | Imm preadip | Mgst1 |
| Cxcl1.1 | 9.22E-232 | -0.9754334 | 0.818 | 0.869 | 1.36E-228 | Imm preadip | Cxcl1 |
| Sdc4.2 | 1.27E-229 | 0.51029907 | 0.944 | 0.874 | 1.87E-226 | Imm preadip | Sdc4 |
| Gas1.1 | 9.41E-229 | 0.58715618 | 0.831 | 0.698 | 1.39E-225 | Imm preadip | Gas1 |
| Cst3.2 | 7.88E-226 | 0.35315593 | 0.992 | 0.979 | 1.16E-222 | Imm preadip | Cst3 |
| Npy1r | 1.47E-224 | 0.30886878 | 0.73 | 0.644 | 2.17E-221 | Imm preadip | Npy1r |
| Angptl7 | 1.50E-213 | 0.3056199 | 0.569 | 0.326 | 2.21E-210 | Imm preadip | Angptl7 |
| Pla1a.1 | 2.09E-209 | -0.5199366 | 0.548 | 0.635 | 3.07E-206 | Imm preadip | Pla1a |
| Cd55.1 | 4.72E-208 | -0.7350489 | 0.84 | 0.895 | 6.94E-205 | Imm preadip | Cd55 |
| Tpm1.2 | 3.81E-202 | 0.47708626 | 0.867 | 0.757 | 5.60E-199 | Imm preadip | Tpm1 |

| | | | | | | | |
|---|---|---|---|---|---|---|---|
| Col12a1.1 | 2.62E-200 | 0.67497979 | 0.623 | 0.436 | 3.85E-197 | Imm preadip | Col12a1 |
| Aldh1a3.1 | 3.89E-195 | -0.5779439 | 0.497 | 0.604 | 5.73E-192 | Imm preadip | Aldh1a3 |
| Gpx3.2 | 1.58E-194 | 0.62078572 | 0.984 | 0.973 | 2.32E-191 | Imm preadip | Gpx3 |
| Ptgs2.2 | 2.57E-187 | -0.8078184 | 0.738 | 0.762 | 3.78E-184 | Imm preadip | Ptgs2 |
| Myoc.1 | 6.88E-187 | 0.56310493 | 0.664 | 0.475 | 1.01E-183 | Imm preadip | Myoc |
| Sbsn.2 | 1.83E-186 | -1.0116598 | 0.631 | 0.673 | 2.69E-183 | Imm preadip | Sbsn |
| Ly6c1.1 | 1.97E-184 | -0.4056106 | 0.954 | 0.969 | 2.90E-181 | Imm preadip | Ly6c1 |
| Postn.1 | 5.65E-182 | -0.4108145 | 0.931 | 0.977 | 8.32E-179 | Imm preadip | Postn |
| Adamtsl2.1 | 4.64E-181 | 0.53334031 | 0.529 | 0.202 | 6.82E-178 | Imm preadip | Adamtsl2 |
| Prkg2.1 | 3.29E-179 | -0.4035855 | 0.563 | 0.643 | 4.85E-176 | Imm preadip | Prkg2 |
| Thbs1.1 | 5.28E-178 | 0.57882207 | 0.773 | 0.618 | 7.78E-175 | Imm preadip | Thbs1 |
| Fmod.2 | 8.94E-172 | 0.6743341 | 0.524 | 0.263 | 1.32E-168 | Imm preadip | Fmod |
| Ackr3.1 | 6.78E-169 | -0.6842731 | 0.733 | 0.785 | 9.99E-166 | Imm preadip | Ackr3 |
| C4b.1 | 9.32E-167 | -0.6574663 | 0.59 | 0.682 | 1.37E-163 | Imm preadip | C4b |
| Pmp22 | 1.85E-163 | 0.36071933 | 0.946 | 0.899 | 2.72E-160 | Imm preadip | Pmp22 |
| Sned1.1 | 3.61E-161 | -0.6576279 | 0.545 | 0.605 | 5.32E-158 | Imm preadip | Sned1 |
| Car3.2 | 6.34E-157 | -0.475215 | 0.738 | 0.555 | 9.33E-154 | Imm preadip | Car3 |
| Tmsb4x.1 | 1.78E-148 | -0.339995 | 0.986 | 0.983 | 2.62E-145 | Imm preadip | Tmsb4x |
| Fgl2.1 | 2.27E-147 | 0.38045598 | 0.965 | 0.897 | 3.34E-144 | Imm preadip | Fgl2 |
| Hba-a1.2 | 4.21E-145 | 0.3912352 | 0.987 | 0.98 | 6.20E-142 | Imm preadip | Hba-a1 |
| Cthrc1 | 4.99E-139 | 0.55531745 | 0.697 | 0.555 | 7.34E-136 | Imm preadip | Cthrc1 |
| Rgcc | 2.41E-134 | 0.31847983 | 0.69 | 0.608 | 3.55E-131 | Imm preadip | Rgcc |
| Peg3.1 | 2.92E-132 | 0.33877255 | 0.841 | 0.704 | 4.30E-129 | Imm preadip | Peg3 |
| Hbb-bs.2 | 3.02E-129 | 0.39277907 | 0.999 | 0.998 | 4.44E-126 | Imm preadip | Hbb-bs |
| Mustn1.1 | 1.73E-128 | -0.4496115 | 0.78 | 0.799 | 2.55E-125 | Imm preadip | Mustn1 |
| Ptn.2 | 7.66E-126 | -0.8471305 | 0.497 | 0.571 | 1.13E-122 | Imm preadip | Ptn |
| Hmcn2.1 | 2.39E-125 | 0.58897061 | 0.54 | 0.284 | 3.51E-122 | Imm preadip | Hmcn2 |
| Ccl2.2 | 2.79E-121 | -0.9962383 | 0.761 | 0.786 | 4.10E-118 | Imm preadip | Ccl2 |
| Plod2 | 5.76E-121 | -0.2696684 | 0.439 | 0.566 | 8.47E-118 | Imm preadip | Plod2 |
| Ctgf | 5.40E-120 | 0.42840878 | 0.532 | 0.359 | 7.95E-117 | Imm preadip | Ctgf |
| Irf7.1 | 2.62E-118 | -0.3101275 | 0.629 | 0.681 | 3.85E-115 | Imm preadip | Irf7 |
| Clec11a.1 | 4.78E-118 | 0.4305984 | 0.644 | 0.501 | 7.03E-115 | Imm preadip | Clec11a |
| Fibin | 5.25E-116 | 0.26466259 | 0.553 | 0.375 | 7.73E-113 | Imm preadip | Fibin |
| Slit2.1 | 6.54E-116 | 0.44883479 | 0.609 | 0.456 | 9.63E-113 | Imm preadip | Slit2 |
| Art3 | 9.00E-114 | 0.26516016 | 0.509 | 0.321 | 1.33E-110 | Imm preadip | Art3 |
| Tnfaip2 | 3.64E-111 | -0.4286838 | 0.735 | 0.769 | 5.36E-108 | Imm preadip | Tnfaip2 |
| Ccl7.2 | 8.48E-111 | -0.6480871 | 0.847 | 0.875 | 1.25E-107 | Imm preadip | Ccl7 |
| Npm1 | 1.28E-104 | -0.2688494 | 0.93 | 0.923 | 1.89E-101 | Imm preadip | Npm1 |
| Prss23.2 | 2.34E-101 | -0.4078104 | 0.922 | 0.938 | 3.44E-98 | Imm preadip | Prss23 |

| | | | | | | | |
|---|---|---|---|---|---|---|---|
| Ccl5 | 3.48E-99 | -0.338316 | 0.35 | 0.429 | 5.13E-96 | Imm preadip | Ccl5 |
| Dpp4.2 | 5.47E-99 | -0.6473173 | 0.895 | 0.89 | 8.05E-96 | Imm preadip | Dpp4 |
| Ctsk.1 | 5.95E-99 | 0.29885287 | 0.935 | 0.894 | 8.76E-96 | Imm preadip | Ctsk |
| Il33.1 | 2.43E-97 | -0.6052416 | 0.506 | 0.584 | 3.58E-94 | Imm preadip | Il33 |
| Cryab | 1.97E-96 | 0.31890478 | 0.909 | 0.845 | 2.89E-93 | Imm preadip | Cryab |
| Col15a1.2 | 7.02E-96 | 0.2545071 | 0.739 | 0.564 | 1.03E-92 | Imm preadip | Col15a1 |
| Cited2.1 | 3.31E-94 | -0.3098732 | 0.395 | 0.499 | 4.88E-91 | Imm preadip | Cited2 |
| Cyr61.1 | 9.86E-93 | 0.31666005 | 0.74 | 0.615 | 1.45E-89 | Imm preadip | Cyr61 |
| Hspa1a.2 | 4.67E-91 | 0.29777762 | 0.871 | 0.772 | 6.88E-88 | Imm preadip | Hspa1a |
| Hbb-bt.2 | 8.03E-86 | 0.41699251 | 0.943 | 0.909 | 1.18E-82 | Imm preadip | Hbb-bt |
| Phlda1.1 | 3.90E-81 | -0.5189539 | 0.756 | 0.768 | 5.74E-78 | Imm preadip | Phlda1 |
| Pappa2.1 | 8.56E-81 | 0.52421591 | 0.511 | 0.308 | 1.26E-77 | Imm preadip | Pappa2 |
| Anxa3.2 | 2.03E-79 | -0.7927783 | 0.911 | 0.912 | 2.99E-76 | Imm preadip | Anxa3 |
| Tmem176b.2 | 6.80E-79 | -0.2570545 | 0.416 | 0.472 | 1.00E-75 | Imm preadip | Tmem176b |
| Tgm2.1 | 9.07E-78 | 0.30111603 | 0.591 | 0.457 | 1.33E-74 | Imm preadip | Tgm2 |
| Gadd45b.2 | 1.61E-75 | 0.34393976 | 0.826 | 0.763 | 2.37E-72 | Imm preadip | Gadd45b |
| Ier3.1 | 2.18E-74 | 0.40043148 | 0.906 | 0.852 | 3.20E-71 | Imm preadip | Ier3 |
| Col6a5 | 6.45E-71 | -0.3705724 | 0.38 | 0.442 | 9.50E-68 | Imm preadip | Col6a5 |
| Ncam1.1 | 8.30E-68 | 0.29335412 | 0.474 | 0.315 | 1.22E-64 | Imm preadip | Ncam1 |
| Grem2.1 | 5.53E-67 | 0.36235666 | 0.557 | 0.452 | 8.14E-64 | Imm preadip | Grem2 |
| Thbs2.1 | 3.43E-66 | 0.31220476 | 0.763 | 0.689 | 5.04E-63 | Imm preadip | Thbs2 |
| Fabp4.2 | 5.00E-66 | -1.0106058 | 0.818 | 0.742 | 7.36E-63 | Imm preadip | Fabp4 |
| Sdpr | 5.38E-59 | 0.28098873 | 0.648 | 0.536 | 7.92E-56 | Imm preadip | Sdpr |
| Rrad.2 | 6.56E-57 | 0.66438484 | 0.6 | 0.501 | 9.65E-54 | Imm preadip | Rrad |
| Rgs16.1 | 4.44E-56 | -0.2920413 | 0.248 | 0.271 | 6.54E-53 | Imm preadip | Rgs16 |
| Sfrp4 | 6.48E-53 | 0.68636594 | 0.818 | 0.806 | 9.54E-50 | Imm preadip | Sfrp4 |
| Igfbp3.1 | 2.03E-51 | 0.32809177 | 0.68 | 0.556 | 2.99E-48 | Imm preadip | Igfbp3 |
| Hba-a2.2 | 1.58E-50 | 0.38918511 | 0.976 | 0.964 | 2.32E-47 | Imm preadip | Hba-a2 |
| Sod2 | 4.42E-48 | -0.2717356 | 0.687 | 0.699 | 6.50E-45 | Imm preadip | Sod2 |
| Krtdap.1 | 6.13E-48 | -0.4798072 | 0.583 | 0.527 | 9.02E-45 | Imm preadip | Krtdap |
| Socs3.1 | 1.90E-47 | -0.3971115 | 0.765 | 0.756 | 2.80E-44 | Imm preadip | Socs3 |
| Matn2.1 | 9.57E-46 | 0.26774531 | 0.577 | 0.45 | 1.41E-42 | Imm preadip | Matn2 |
| Pi16.2 | 2.46E-45 | -0.7357032 | 0.95 | 0.921 | 3.62E-42 | Imm preadip | Pi16 |
| Psmb8 | 8.84E-45 | -0.2977034 | 0.641 | 0.658 | 1.30E-41 | Imm preadip | Psmb8 |
| Cxcl9.1 | 2.35E-44 | -0.2895558 | 0.321 | 0.314 | 3.46E-41 | Imm preadip | Cxcl9 |
| Lgmn | 9.39E-44 | -0.2591274 | 0.467 | 0.518 | 1.38E-40 | Imm preadip | Lgmn |
| Mafb.1 | 2.38E-42 | -0.3274243 | 0.527 | 0.592 | 3.50E-39 | Imm preadip | Mafb |
| AW112010.1 | 6.68E-41 | -0.4509532 | 0.687 | 0.664 | 9.83E-38 | Imm preadip | AW112010 |
| AY036118.1 | 1.78E-38 | 0.27899116 | 0.853 | 0.809 | 2.62E-35 | Imm preadip | AY036118 |

| | | | | | | | |
|---|---|---|---|---|---|---|---|
| Hmox1 | 8.83E-38 | -0.3307777 | 0.664 | 0.645 | 1.30E-34 | Imm preadip | Hmox1 |
| Fabp5 | 5.06E-37 | -0.5172773 | 0.479 | 0.522 | 7.44E-34 | Imm preadip | Fabp5 |
| Cotl1.1 | 1.46E-36 | -0.2757762 | 0.255 | 0.366 | 2.15E-33 | Imm preadip | Cotl1 |
| Cxcl2.2 | 9.26E-36 | -0.6223598 | 0.689 | 0.683 | 1.36E-32 | Imm preadip | Cxcl2 |
| Gem.2 | 1.04E-35 | -0.2790675 | 0.703 | 0.752 | 1.53E-32 | Imm preadip | Gem |
| Apoe.2 | 1.55E-35 | -1.0931547 | 0.539 | 0.546 | 2.28E-32 | Imm preadip | Apoe |
| Steap4.1 | 4.66E-33 | -0.2799103 | 0.56 | 0.542 | 6.86E-30 | Imm preadip | Steap4 |
| Gdf10.1 | 2.20E-32 | -0.3177651 | 0.29 | 0.335 | 3.24E-29 | Imm preadip | Gdf10 |
| Col18a1 | 1.33E-31 | -0.2636718 | 0.709 | 0.65 | 1.96E-28 | Imm preadip | Col18a1 |
| Abca8a.1 | 4.19E-31 | 0.31260513 | 0.727 | 0.633 | 6.17E-28 | Imm preadip | Abca8a |
| Gas7.1 | 4.35E-30 | -0.3900254 | 0.796 | 0.748 | 6.40E-27 | Imm preadip | Gas7 |
| Trf | 1.36E-29 | -0.2726003 | 0.573 | 0.529 | 2.01E-26 | Imm preadip | Trf |
| Il6 | 4.84E-29 | -0.4408031 | 0.672 | 0.638 | 7.13E-26 | Imm preadip | Il6 |
| Cxcl10 | 2.31E-25 | -0.2617766 | 0.581 | 0.622 | 3.39E-22 | Imm preadip | Cxcl10 |
| Ddah1.1 | 1.88E-23 | 0.29277254 | 0.551 | 0.489 | 2.77E-20 | Imm preadip | Ddah1 |
| Plagl1.1 | 1.41E-22 | -0.2865323 | 0.627 | 0.628 | 2.07E-19 | Imm preadip | Plagl1 |
| Nov.1 | 6.45E-22 | 0.71949483 | 0.485 | 0.386 | 9.50E-19 | Imm preadip | Nov |
| Epha4.1 | 2.26E-19 | 0.26398463 | 0.462 | 0.337 | 3.33E-16 | Imm preadip | Epha4 |
| Hspd1.1 | 6.69E-19 | -0.27589 | 0.833 | 0.798 | 9.85E-16 | Imm preadip | Hspd1 |
| Cxcl12.2 | 2.97E-18 | -0.4821937 | 0.76 | 0.706 | 4.37E-15 | Imm preadip | Cxcl12 |
| Col8a1.2 | 3.52E-15 | 0.68215545 | 0.447 | 0.279 | 5.18E-12 | Imm preadip | Col8a1 |
| Maf | 5.91E-14 | -0.2536844 | 0.539 | 0.522 | 8.70E-11 | Imm preadip | Maf |
| Ccl8.1 | 3.47E-13 | -0.3239749 | 0.612 | 0.588 | 5.11E-10 | Imm preadip | Ccl8 |
| Angpt4 | 1.47E-11 | 0.35672046 | 0.462 | 0.389 | 2.16E-08 | Imm preadip | Angpt4 |
| Dbi.2 | 4.45E-11 | -0.3211524 | 0.949 | 0.916 | 6.56E-08 | Imm preadip | Dbi |
| Wisp1.1 | 7.10E-11 | 0.30829855 | 0.421 | 0.243 | 1.04E-07 | Imm preadip | Wisp1 |
| Fhl1.1 | 8.52E-11 | -0.2652 | 0.459 | 0.463 | 1.25E-07 | Imm preadip | Fhl1 |
| Npr3 | 2.32E-05 | 0.26244224 | 0.376 | 0.255 | 0.03417642 | Imm preadip | Npr3 |
| Crispld2.1 | 3.46E-05 | 0.32104313 | 0.531 | 0.552 | 0.05089077 | Imm preadip | Crispld2 |
| Thbs4.1 | 0.00926057 | 0.72106122 | 0.395 | 0.245 | 1 | Imm preadip | Thbs4 |
| Apoe.3 | 0 | 1.97394108 | 0.75 | 0.462 | 0 | Comm preadip | Apoe |
| Fabp4.3 | 0 | 1.63033915 | 0.835 | 0.748 | 0 | Comm preadip | Fabp4 |
| Cxcl12.3 | 0 | 1.37458762 | 0.9 | 0.66 | 0 | Comm preadip | Cxcl12 |
| Ptn.3 | 0 | 1.36693992 | 0.688 | 0.484 | 0 | Comm preadip | Ptn |
| Mmp3.2 | 0 | 1.14871836 | 0.848 | 0.658 | 0 | Comm preadip | Mmp3 |
| Igfbp7.2 | 0 | 1.08878407 | 0.977 | 0.885 | 0 | Comm preadip | Igfbp7 |
| Col4a1.2 | 0 | 1.0510851 | 0.961 | 0.835 | 0 | Comm preadip | Col4a1 |
| C4b.2 | 0 | 0.95049186 | 0.787 | 0.59 | 0 | Comm preadip | C4b |
| Sned1.2 | 0 | 0.89388562 | 0.73 | 0.523 | 0 | Comm preadip | Sned1 |

| | | | | | | | |
|---|---|---|---|---|---|---|---|
| Col4a2.2 | 0 | 0.77656694 | 0.928 | 0.782 | 0 | Comm preadip | Col4a2 |
| Steap4.2 | 0 | 0.58459508 | 0.715 | 0.484 | 0 | Comm preadip | Steap4 |
| Agt | 0 | 0.28948218 | 0.701 | 0.346 | 0 | Comm preadip | Agt |
| Col1a1.1 | 0 | -0.6152587 | 0.999 | 1 | 0 | Comm preadip | Col1a1 |
| Pmepa1.2 | 0 | -0.72503 | 0.823 | 0.917 | 0 | Comm preadip | Pmepa1 |
| Prss23.3 | 0 | -0.737385 | 0.887 | 0.949 | 0 | Comm preadip | Prss23 |
| Aspn.2 | 0 | -0.7586605 | 0.897 | 0.971 | 0 | Comm preadip | Aspn |
| Sfrp4.1 | 0 | -0.9597647 | 0.731 | 0.841 | 0 | Comm preadip | Sfrp4 |
| Dpp4.3 | 0 | -1.0229261 | 0.809 | 0.924 | 0 | Comm preadip | Dpp4 |
| Lox.2 | 0 | -1.2266581 | 0.77 | 0.881 | 0 | Comm preadip | Lox |
| Anxa3.3 | 0 | -1.4196999 | 0.836 | 0.941 | 0 | Comm preadip | Anxa3 |
| Sfrp2.2 | 0 | -1.4197411 | 0.778 | 0.888 | 0 | Comm preadip | Sfrp2 |
| Pi16.3 | 0 | -1.9075493 | 0.843 | 0.967 | 0 | Comm preadip | Pi16 |
| Eln.2 | 0 | -2.3856405 | 0.697 | 0.901 | 0 | Comm preadip | Eln |
| AW112010.2 | 8.48E-291 | 0.8029451 | 0.785 | 0.63 | 1.25E-287 | Comm preadip | AW112010 |
| Fbn2.2 | 3.36E-287 | -0.7307522 | 0.455 | 0.664 | 4.95E-284 | Comm preadip | Fbn2 |
| Sparcl1.2 | 3.35E-279 | 0.73347441 | 0.666 | 0.487 | 4.93E-276 | Comm preadip | Sparcl1 |
| Lpl.1 | 3.28E-271 | 0.88318217 | 0.967 | 0.941 | 4.82E-268 | Comm preadip | Lpl |
| Mfap4.3 | 1.17E-270 | -1.8635344 | 0.715 | 0.754 | 1.73E-267 | Comm preadip | Mfap4 |
| Kitl.1 | 6.26E-269 | 0.67437069 | 0.676 | 0.469 | 9.22E-266 | Comm preadip | Kitl |
| Pdgfrl.2 | 4.22E-265 | -0.6746683 | 0.708 | 0.814 | 6.21E-262 | Comm preadip | Pdgfrl |
| Itgbl1.3 | 3.74E-263 | -0.8583723 | 0.729 | 0.773 | 5.50E-260 | Comm preadip | Itgbl1 |
| Tnfaip6.1 | 6.54E-262 | 0.84990075 | 0.955 | 0.924 | 9.62E-259 | Comm preadip | Tnfaip6 |
| Tmem176b.3 | 2.15E-260 | 0.80555882 | 0.623 | 0.382 | 3.17E-257 | Comm preadip | Tmem176b |
| Phlda1.2 | 3.49E-244 | 0.91117801 | 0.836 | 0.735 | 5.13E-241 | Comm preadip | Phlda1 |
| Ndnf | 1.00E-241 | 0.31891168 | 0.597 | 0.366 | 1.48E-238 | Comm preadip | Ndnf |
| Cldn10 | 1.71E-239 | -0.4740208 | 0.759 | 0.791 | 2.52E-236 | Comm preadip | Cldn10 |
| Col15a1.3 | 3.13E-238 | 0.6233674 | 0.801 | 0.568 | 4.61E-235 | Comm preadip | Col15a1 |
| Cxcl13.1 | 2.72E-231 | 0.41275222 | 0.83 | 0.68 | 4.01E-228 | Comm preadip | Cxcl13 |
| Gpx3.3 | 8.34E-231 | -0.5653613 | 0.955 | 0.985 | 1.23E-227 | Comm preadip | Gpx3 |
| Socs3.2 | 5.64E-223 | 0.75985708 | 0.84 | 0.728 | 8.31E-220 | Comm preadip | Socs3 |
| Enpp2 | 3.11E-221 | 0.33128622 | 0.634 | 0.424 | 4.58E-218 | Comm preadip | Enpp2 |
| Gap43.2 | 7.94E-218 | -0.6337795 | 0.457 | 0.63 | 1.17E-214 | Comm preadip | Gap43 |
| Gucy1a3 | 7.50E-217 | 0.44777718 | 0.546 | 0.24 | 1.10E-213 | Comm preadip | Gucy1a3 |
| Mgst1.1 | 1.65E-206 | 0.54552068 | 0.938 | 0.895 | 2.43E-203 | Comm preadip | Mgst1 |
| Cilp.3 | 1.30E-203 | -0.9026734 | 0.489 | 0.603 | 1.91E-200 | Comm preadip | Cilp |
| Gem.3 | 4.95E-199 | 0.73898109 | 0.822 | 0.698 | 7.28E-196 | Comm preadip | Gem |
| Wisp2.2 | 3.17E-195 | -0.9497479 | 0.566 | 0.655 | 4.67E-192 | Comm preadip | Wisp2 |
| Ramp1.1 | 8.02E-195 | -0.3794136 | 0.492 | 0.608 | 1.18E-191 | Comm preadip | Ramp1 |

| | | | | | | | |
|---|---|---|---|---|---|---|---|
| Mafb.2 | 1.50E-192 | 0.968133 | 0.692 | 0.517 | 2.20E-189 | Comm preadip | Mafb |
| Hp | 1.23E-184 | 0.30508043 | 0.553 | 0.326 | 1.82E-181 | Comm preadip | Hp |
| Col18a1.1 | 2.18E-182 | 0.38894282 | 0.754 | 0.642 | 3.21E-179 | Comm preadip | Col18a1 |
| Mgll.1 | 3.61E-181 | -0.5001491 | 0.691 | 0.743 | 5.31E-178 | Comm preadip | Mgll |
| Ackr3.2 | 1.41E-170 | -0.6261869 | 0.72 | 0.782 | 2.07E-167 | Comm preadip | Ackr3 |
| Cgref1 | 2.11E-168 | -0.3989302 | 0.635 | 0.695 | 3.11E-165 | Comm preadip | Cgref1 |
| Fhl1.2 | 2.79E-167 | -0.26443 | 0.351 | 0.505 | 4.10E-164 | Comm preadip | Fhl1 |
| Mgst3.1 | 1.79E-165 | 0.39064345 | 0.608 | 0.37 | 2.64E-162 | Comm preadip | Mgst3 |
| Tgfbi.2 | 8.83E-162 | -0.576745 | 0.72 | 0.788 | 1.30E-158 | Comm preadip | Tgfbi |
| Osr2.1 | 1.96E-160 | -0.4780361 | 0.446 | 0.608 | 2.89E-157 | Comm preadip | Osr2 |
| mt-Cytb.2 | 8.91E-153 | 0.30743196 | 0.998 | 0.997 | 1.31E-149 | Comm preadip | mt-Cytb |
| C1qtnf3.3 | 2.40E-152 | -1.0002507 | 0.515 | 0.602 | 3.54E-149 | Comm preadip | C1qtnf3 |
| Cd55.2 | 2.66E-149 | -0.6208313 | 0.863 | 0.877 | 3.91E-146 | Comm preadip | Cd55 |
| Nkain4.2 | 6.33E-148 | -0.3825752 | 0.679 | 0.672 | 9.31E-145 | Comm preadip | Nkain4 |
| Dbi.3 | 4.86E-143 | 0.73581627 | 0.937 | 0.926 | 7.15E-140 | Comm preadip | Dbi |
| 1500015O10Rik.2 | 7.16E-139 | 0.86889541 | 0.805 | 0.711 | 1.05E-135 | Comm preadip | 1500015O10Rik |
| Cthrc1.1 | 7.26E-131 | -0.5512134 | 0.544 | 0.638 | 1.07E-127 | Comm preadip | Cthrc1 |
| Tmem176a.1 | 3.53E-128 | 0.48425719 | 0.523 | 0.296 | 5.20E-125 | Comm preadip | Tmem176a |
| Krtdap.2 | 8.02E-126 | -0.4106989 | 0.433 | 0.595 | 1.18E-122 | Comm preadip | Krtdap |
| Hpgd.2 | 2.46E-123 | -0.2523143 | 0.761 | 0.806 | 3.62E-120 | Comm preadip | Hpgd |
| Mustn1.2 | 5.48E-123 | -0.569961 | 0.776 | 0.797 | 8.07E-120 | Comm preadip | Mustn1 |
| Adamts4.1 | 6.92E-121 | 0.49734177 | 0.78 | 0.667 | 1.02E-117 | Comm preadip | Adamts4 |
| Sowahc | 9.31E-120 | 0.38268915 | 0.607 | 0.475 | 1.37E-116 | Comm preadip | Sowahc |
| Angptl4 | 2.67E-118 | 0.49257516 | 0.475 | 0.262 | 3.93E-115 | Comm preadip | Angptl4 |
| Angptl1.2 | 1.42E-116 | -0.4393442 | 0.619 | 0.716 | 2.08E-113 | Comm preadip | Angptl1 |
| Sox9 | 7.82E-112 | 0.4298802 | 0.476 | 0.248 | 1.15E-108 | Comm preadip | Sox9 |
| Serpinb1a | 2.69E-111 | 0.28876472 | 0.668 | 0.497 | 3.96E-108 | Comm preadip | Serpinb1a |
| Smpd3.3 | 7.14E-111 | -0.8948428 | 0.728 | 0.789 | 1.05E-107 | Comm preadip | Smpd3 |
| Ccl8.2 | 2.62E-97 | 0.71188508 | 0.672 | 0.568 | 3.86E-94 | Comm preadip | Ccl8 |
| Cryab.1 | 8.40E-95 | -0.3709834 | 0.835 | 0.884 | 1.24E-91 | Comm preadip | Cryab |
| Gbp2.1 | 7.77E-93 | 0.55385674 | 0.733 | 0.619 | 1.14E-89 | Comm preadip | Gbp2 |
| Prlr.1 | 3.74E-92 | 0.4687884 | 0.487 | 0.28 | 5.50E-89 | Comm preadip | Prlr |
| Gadd45g.2 | 1.16E-87 | 0.52642943 | 0.897 | 0.852 | 1.71E-84 | Comm preadip | Gadd45g |
| Icam1.2 | 6.10E-87 | 0.2984747 | 0.888 | 0.769 | 8.98E-84 | Comm preadip | Icam1 |
| Tnfsf9.2 | 6.76E-87 | 0.28221695 | 0.785 | 0.724 | 9.95E-84 | Comm preadip | Tnfsf9 |
| Akr1c18.2 | 2.58E-84 | -0.8543153 | 0.662 | 0.681 | 3.79E-81 | Comm preadip | Akr1c18 |
| Hmox1.1 | 8.67E-83 | 0.36829288 | 0.749 | 0.615 | 1.28E-79 | Comm preadip | Hmox1 |
| Gdf10.2 | 4.04E-81 | 0.54235528 | 0.47 | 0.258 | 5.95E-78 | Comm preadip | Gdf10 |
| Tnfaip2.1 | 4.59E-80 | 0.43504065 | 0.801 | 0.738 | 6.76E-77 | Comm preadip | Tnfaip2 |

| | | | | | | | |
|---|---|---|---|---|---|---|---|
| Timp1.1 | 2.14E-78 | 0.52650846 | 0.805 | 0.725 | 3.15E-75 | Comm preadip | Timp1 |
| Vcam1.1 | 1.46E-77 | 0.42706807 | 0.767 | 0.677 | 2.16E-74 | Comm preadip | Vcam1 |
| Gbp5 | 1.60E-77 | 0.37431706 | 0.576 | 0.446 | 2.35E-74 | Comm preadip | Gbp5 |
| Adamtsl2.2 | 1.41E-76 | -0.3033417 | 0.283 | 0.351 | 2.07E-73 | Comm preadip | Adamtsl2 |
| Gpm6b.2 | 7.26E-75 | 0.48713259 | 0.589 | 0.443 | 1.07E-71 | Comm preadip | Gpm6b |
| Iigp1.1 | 1.35E-74 | 0.63221089 | 0.646 | 0.538 | 1.99E-71 | Comm preadip | Iigp1 |
| Nov.2 | 2.57E-74 | -0.6985995 | 0.359 | 0.452 | 3.78E-71 | Comm preadip | Nov |
| Pcsk5.2 | 1.07E-72 | -0.4997607 | 0.671 | 0.686 | 1.57E-69 | Comm preadip | Pcsk5 |
| Hspd1.2 | 1.07E-72 | 0.50377853 | 0.848 | 0.798 | 1.58E-69 | Comm preadip | Hspd1 |
| Sod2.1 | 1.63E-71 | 0.29995872 | 0.77 | 0.665 | 2.40E-68 | Comm preadip | Sod2 |
| Saa3 | 2.42E-69 | 0.47798251 | 0.443 | 0.242 | 3.56E-66 | Comm preadip | Saa3 |
| C1qbp | 6.21E-69 | 0.2669868 | 0.686 | 0.548 | 9.14E-66 | Comm preadip | C1qbp |
| Ptx3.2 | 8.47E-66 | 0.44457978 | 0.864 | 0.783 | 1.25E-62 | Comm preadip | Ptx3 |
| Cp | 1.70E-62 | 0.2891086 | 0.789 | 0.722 | 2.50E-59 | Comm preadip | Cp |
| Ptgs2.3 | 1.46E-60 | -0.4463425 | 0.697 | 0.774 | 2.15E-57 | Comm preadip | Ptgs2 |
| Has1.2 | 2.44E-59 | -0.2719947 | 0.613 | 0.705 | 3.59E-56 | Comm preadip | Has1 |
| Ccl19 | 5.49E-59 | 0.39546803 | 0.511 | 0.356 | 8.08E-56 | Comm preadip | Ccl19 |
| Plagl1.2 | 6.80E-59 | -0.3020918 | 0.568 | 0.651 | 1.00E-55 | Comm preadip | Plagl1 |
| Ccl5.1 | 1.41E-58 | 0.56661485 | 0.494 | 0.36 | 2.08E-55 | Comm preadip | Ccl5 |
| Trf.1 | 1.89E-58 | 0.29215245 | 0.606 | 0.523 | 2.78E-55 | Comm preadip | Trf |
| Psmb8.1 | 6.88E-58 | 0.37962508 | 0.703 | 0.631 | 1.01E-54 | Comm preadip | Psmb8 |
| C3.2 | 1.50E-57 | 0.53469275 | 0.802 | 0.79 | 2.21E-54 | Comm preadip | C3 |
| Ldha.1 | 9.44E-57 | 0.29840398 | 0.85 | 0.817 | 1.39E-53 | Comm preadip | Ldha |
| Ccl11.2 | 1.83E-55 | 0.34714 | 0.584 | 0.424 | 2.70E-52 | Comm preadip | Ccl11 |
| Pcsk6.1 | 1.66E-53 | -0.5217372 | 0.646 | 0.613 | 2.45E-50 | Comm preadip | Pcsk6 |
| Tubb5.1 | 2.02E-52 | 0.28573389 | 0.893 | 0.864 | 2.97E-49 | Comm preadip | Tubb5 |
| Sbsn.3 | 7.19E-47 | -0.7485542 | 0.627 | 0.667 | 1.06E-43 | Comm preadip | Sbsn |
| Cd74 | 1.03E-44 | 0.34432703 | 0.47 | 0.337 | 1.51E-41 | Comm preadip | Cd74 |
| Tspo.2 | 1.63E-44 | 0.32941649 | 0.875 | 0.83 | 2.39E-41 | Comm preadip | Tspo |
| Hes1 | 2.45E-44 | -0.3176075 | 0.528 | 0.56 | 3.61E-41 | Comm preadip | Hes1 |
| Socs1.1 | 7.87E-43 | 0.46341469 | 0.561 | 0.436 | 1.16E-39 | Comm preadip | Socs1 |
| Lgr5.2 | 2.41E-42 | -0.3349035 | 0.49 | 0.494 | 3.55E-39 | Comm preadip | Lgr5 |
| Hspa1b.3 | 3.18E-42 | 0.2853956 | 0.903 | 0.813 | 4.69E-39 | Comm preadip | Hspa1b |
| Lmo4 | 2.61E-39 | 0.25613945 | 0.648 | 0.561 | 3.84E-36 | Comm preadip | Lmo4 |
| Fmod.3 | 1.74E-38 | -0.3269668 | 0.371 | 0.364 | 2.56E-35 | Comm preadip | Fmod |
| Fmo2.2 | 1.53E-37 | -0.5523059 | 0.6 | 0.6 | 2.26E-34 | Comm preadip | Fmo2 |
| Hnrnpa1 | 1.25E-36 | 0.25473357 | 0.706 | 0.619 | 1.85E-33 | Comm preadip | Hnrnpa1 |
| Serpina3n.1 | 6.74E-35 | 0.33533042 | 0.59 | 0.496 | 9.92E-32 | Comm preadip | Serpina3n |
| Nrk | 1.59E-34 | -0.2679525 | 0.54 | 0.574 | 2.35E-31 | Comm preadip | Nrk |

| | | | | | | | |
|---|---|---|---|---|---|---|---|
| Fabp5.1 | 2.21E-34 | 0.8423207 | 0.563 | 0.482 | 3.25E-31 | Comm preadip | Fabp5 |
| H2-Q7 | 8.34E-34 | 0.31560914 | 0.479 | 0.377 | 1.23E-30 | Comm preadip | H2-Q7 |
| Hba-a2.3 | 8.72E-34 | -0.7668703 | 0.97 | 0.968 | 1.28E-30 | Comm preadip | Hba-a2 |
| Cxcl1.2 | 1.32E-33 | 0.33410446 | 0.87 | 0.84 | 1.94E-30 | Comm preadip | Cxcl1 |
| Hspb1.1 | 1.75E-33 | 0.4799145 | 0.751 | 0.679 | 2.57E-30 | Comm preadip | Hspb1 |
| Il6.1 | 2.69E-32 | 0.41688894 | 0.683 | 0.639 | 3.95E-29 | Comm preadip | Il6 |
| Stat1 | 9.80E-32 | 0.28115546 | 0.6 | 0.515 | 1.44E-28 | Comm preadip | Stat1 |
| Cxcl9.2 | 3.86E-31 | 0.95841331 | 0.408 | 0.281 | 5.68E-28 | Comm preadip | Cxcl9 |
| Gbp7 | 7.79E-31 | 0.27314887 | 0.579 | 0.482 | 1.15E-27 | Comm preadip | Gbp7 |
| Mif.1 | 2.67E-29 | 0.25277573 | 0.763 | 0.711 | 3.93E-26 | Comm preadip | Mif |
| Ctla2a.1 | 4.27E-29 | -0.4162173 | 0.333 | 0.389 | 6.29E-26 | Comm preadip | Ctla2a |
| Fxyd5.1 | 2.45E-27 | 0.30474575 | 0.65 | 0.569 | 3.61E-24 | Comm preadip | Fxyd5 |
| Col6a5.1 | 3.27E-25 | 0.45367087 | 0.489 | 0.389 | 4.81E-22 | Comm preadip | Col6a5 |
| Dnajb1.1 | 4.43E-24 | 0.37091648 | 0.917 | 0.877 | 6.52E-21 | Comm preadip | Dnajb1 |
| Car3.3 | 8.27E-24 | 0.90903228 | 0.648 | 0.62 | 1.22E-20 | Comm preadip | Car3 |
| Rgs16.2 | 1.69E-22 | 0.65326489 | 0.386 | 0.213 | 2.48E-19 | Comm preadip | Rgs16 |
| Wnt2.2 | 1.77E-22 | -0.5322207 | 0.746 | 0.686 | 2.61E-19 | Comm preadip | Wnt2 |
| Plin2 | 4.21E-21 | 0.26072481 | 0.463 | 0.351 | 6.20E-18 | Comm preadip | Plin2 |
| Isg15.1 | 5.83E-21 | 0.30539238 | 0.759 | 0.712 | 8.58E-18 | Comm preadip | Isg15 |
| Rrad.3 | 5.81E-20 | 0.41230851 | 0.595 | 0.519 | 8.55E-17 | Comm preadip | Rrad |
| Plvap.1 | 4.05E-18 | 0.3784208 | 0.481 | 0.394 | 5.96E-15 | Comm preadip | Plvap |
| Hbb-bt.3 | 4.54E-17 | -0.7008083 | 0.93 | 0.919 | 6.68E-14 | Comm preadip | Hbb-bt |
| Emb | 1.37E-16 | 0.43153269 | 0.443 | 0.348 | 2.01E-13 | Comm preadip | Emb |
| Mylk.1 | 4.11E-16 | 0.27265512 | 0.494 | 0.41 | 6.04E-13 | Comm preadip | Mylk |
| Ccl2.3 | 7.86E-16 | -0.2713828 | 0.771 | 0.778 | 1.16E-12 | Comm preadip | Ccl2 |
| Sox4 | 3.24E-15 | 0.28019455 | 0.572 | 0.515 | 4.78E-12 | Comm preadip | Sox4 |
| Aldh1a3.2 | 2.71E-14 | -0.3719338 | 0.574 | 0.556 | 3.99E-11 | Comm preadip | Aldh1a3 |
| Hmcn1.2 | 2.96E-14 | -0.29257 | 0.666 | 0.575 | 4.36E-11 | Comm preadip | Hmcn1 |
| Dlk1.1 | 1.73E-13 | 0.27051546 | 0.503 | 0.4 | 2.54E-10 | Comm preadip | Dlk1 |
| Col12a1.2 | 3.28E-12 | -0.3550702 | 0.531 | 0.502 | 4.83E-09 | Comm preadip | Col12a1 |
| Cpxm2.3 | 4.66E-12 | -0.2555577 | 0.512 | 0.494 | 6.86E-09 | Comm preadip | Cpxm2 |
| Id2.1 | 8.66E-12 | 0.25248018 | 0.597 | 0.536 | 1.27E-08 | Comm preadip | Id2 |
| Lbp | 3.83E-11 | 0.26670471 | 0.811 | 0.778 | 5.64E-08 | Comm preadip | Lbp |
| Cxcl10.1 | 2.64E-10 | 0.42095513 | 0.622 | 0.6 | 3.88E-07 | Comm preadip | Cxcl10 |
| S1pr3 | 3.51E-10 | 0.30408289 | 0.422 | 0.279 | 5.16E-07 | Comm preadip | S1pr3 |
| Prkg2.2 | 8.71E-07 | -0.3225937 | 0.644 | 0.598 | 0.0012817 | Comm preadip | Prkg2 |
| Ifi47 | 0.00188135 | 0.28056025 | 0.49 | 0.461 | 1 | Comm preadip | Ifi47 |

**Table S2. Canonical gene expression markers for non-APC cell types in the skin used for manually classifying single-nucleus RNA-seq data.** Identification based on Joost, et al., 2020 scRNA-seq reference dataset (*56*).

| Cell type | Markers |
|---|---|
| Anagen keratinocytes 1 | Robo1$^{high}$, Cux1$^{high}$ |
| Anagen keratinocytes 2 | Tgm6, Prdm1 |
| Anagen keratinocytes 3 | Lgr5, Barx2 |
| Mesenchymal 1 | Pdgfra, Ebf2, Lgr5$^{low}$, Igtbl1 |
| Mesenchymal 2 | Lgr5$^{low}$, Igtbl1, Col1a2$^{high}$, |
| IRS | Fhod3$^{high}$, Lgr5, Efna5$^{high}$, Barx2 |
| Immune 1 | Ptprc |
| Immune 2 | Ptprc |
| Melanocytes | Dct$^{high}$, Tyr$^{high}$ |
| Skeletal muscle | Barx2, Ttn, Myh1 |
| Dermal sheath/papillae | Pdgfra, Col11a1, Ncam2 |
| Adipocytes | Ebf2, Plin1, Adipoq |
| Endothelial | Ptprb, Pecam1 |
| Sebaceous gland | Far2, Serpina3j |
| Unassigned | Armc4, Tpd52l1 |

**Table S3. Canonical gene expression markers for non-APC cell types in the skin used for manually classifying single-cell RNA-seq data.**

| Cell type | Markers |
|---|---|
| Immune | Ptprc |
| Endothelial | Pecam1 |
| Epithelial | Krt14, Krt8 |
| Mature adipocytes | Plin1 |
| Muscle | Myom2 |
| Smooth muscle | Acta2, Myo— |
| Proliferating | Top2a, Mki67, Racgap1 |

**Table S4. Differentially-expressed features of host progenitors vs. transplant progenitors in homeostatic P32 dataset.**

| | p_val | avg_log2FC | pct.1 | pct.2 | p_val_adj |
|---|---|---|---|---|---|
| GFP.CDS | 0 | -1.6978394 | 0.047 | 0.843 | 0 |
| Igfbp4 | 2.82E-209 | 1.3735663 | 0.939 | 0.892 | 4.15E-206 |
| CellTag.UTR | 9.78E-195 | -1.7647919 | 0.066 | 0.759 | 1.44E-191 |
| Postn | 2.61E-162 | -0.7711218 | 0.991 | 0.999 | 3.84E-159 |
| C3 | 1.46E-160 | 1.13339046 | 0.891 | 0.837 | 2.16E-157 |
| Pi16 | 8.46E-130 | 0.72247254 | 0.996 | 0.996 | 1.25E-126 |
| Ptn | 3.29E-93 | -1.2209852 | 0.342 | 0.748 | 4.85E-90 |
| Gpx3 | 1.53E-92 | 0.67263749 | 0.986 | 0.992 | 2.26E-89 |
| Dlk1 | 8.95E-84 | -0.9167945 | 0.107 | 0.657 | 1.32E-80 |
| Ifi205 | 2.05E-82 | 0.73030614 | 0.972 | 0.983 | 3.01E-79 |
| Inmt | 2.01E-67 | 0.28744571 | 0.396 | 0.833 | 2.95E-64 |
| Dmkn | 7.61E-66 | 0.28245153 | 0.367 | 0.773 | 1.12E-62 |
| Cxcl12 | 7.95E-62 | -0.6182448 | 0.372 | 0.758 | 1.17E-58 |
| Il1r2 | 1.24E-60 | 0.91774066 | 0.796 | 0.93 | 1.82E-57 |
| Gap43 | 4.96E-59 | -0.5747273 | 0.598 | 0.868 | 7.30E-56 |
| Sepp1 | 1.64E-55 | 0.51463876 | 0.944 | 0.951 | 2.41E-52 |
| Col6a5 | 2.22E-53 | -0.5593636 | 0.259 | 0.653 | 3.27E-50 |
| Sfrp2 | 3.36E-53 | 0.50944071 | 0.982 | 0.987 | 4.95E-50 |
| Sfrp4 | 1.39E-47 | 0.63444672 | 0.866 | 0.908 | 2.05E-44 |
| Ifi27l2a | 1.89E-45 | 0.46394588 | 0.989 | 0.991 | 2.78E-42 |
| Eln | 2.37E-45 | 0.70729565 | 0.825 | 0.864 | 3.49E-42 |
| S100a4 | 1.33E-43 | -0.2536574 | 0.295 | 0.655 | 1.96E-40 |
| Dpp4 | 9.87E-43 | 0.36946418 | 0.957 | 0.99 | 1.45E-39 |
| Ptgs2 | 1.38E-41 | 0.78948046 | 0.789 | 0.881 | 2.03E-38 |
| Itgbl1 | 4.85E-41 | -0.2654589 | 0.466 | 0.814 | 7.14E-38 |
| Smpd3 | 5.36E-41 | 0.51608401 | 0.921 | 0.931 | 7.89E-38 |
| Cd9 | 9.94E-41 | -0.3994699 | 0.447 | 0.768 | 1.46E-37 |
| Fbn2 | 9.84E-39 | -0.4247932 | 0.418 | 0.766 | 1.45E-35 |
| Hbb-bs | 5.07E-38 | 1.19781533 | 0.999 | 1 | 7.46E-35 |
| Anxa3 | 1.14E-36 | 0.31480204 | 0.985 | 0.993 | 1.67E-33 |
| Cthrc1 | 2.27E-36 | -0.5760565 | 0.463 | 0.763 | 3.34E-33 |
| Hbb-bt | 6.91E-36 | 0.82841927 | 0.833 | 0.962 | 1.02E-32 |
| Ackr3 | 1.03E-35 | 0.47401025 | 0.835 | 0.895 | 1.52E-32 |
| 1500015O10Rik | 1.63E-30 | -0.5929047 | 0.448 | 0.716 | 2.41E-27 |
| Il33 | 5.18E-30 | 0.92705562 | 0.687 | 0.805 | 7.63E-27 |

| | | | | | |
|---|---|---|---|---|---|
| Cdkn1c | 1.98E-28 | -0.4640105 | 0.668 | 0.881 | 2.91E-25 |
| H19 | 7.88E-27 | -0.983408 | 0.325 | 0.62 | 1.16E-23 |
| Timp1 | 5.70E-26 | -0.2850217 | 0.47 | 0.76 | 8.39E-23 |
| Pmepa1 | 6.54E-26 | 0.44400356 | 0.848 | 0.936 | 9.63E-23 |
| Edn1 | 1.51E-25 | 0.34514755 | 0.449 | 0.8 | 2.22E-22 |
| Nrk | 2.33E-25 | -0.4573813 | 0.453 | 0.715 | 3.43E-22 |
| Mustn1 | 8.78E-25 | 0.53074123 | 0.77 | 0.921 | 1.29E-21 |
| Hba-a2 | 2.18E-24 | 1.13001283 | 0.938 | 0.985 | 3.21E-21 |
| Plac8 | 1.70E-23 | 0.37952927 | 0.926 | 0.926 | 2.50E-20 |
| Ccl5 | 5.67E-23 | -0.2969148 | 0.265 | 0.583 | 8.34E-20 |
| Tnfsf9 | 6.51E-19 | 0.27063185 | 0.612 | 0.91 | 9.58E-16 |
| Vcam1 | 1.18E-18 | -0.2900041 | 0.514 | 0.764 | 1.73E-15 |
| Krtdap | 2.42E-18 | 0.47102714 | 0.532 | 0.847 | 3.56E-15 |
| Ccl2 | 3.49E-18 | -0.442025 | 0.737 | 0.944 | 5.13E-15 |
| Fgl2 | 1.62E-17 | 0.53898089 | 0.817 | 0.932 | 2.39E-14 |
| Matn2 | 3.59E-17 | -0.4750697 | 0.257 | 0.582 | 5.29E-14 |
| Hba-a1 | 3.81E-17 | 1.13522835 | 0.972 | 0.99 | 5.61E-14 |
| Tubb5 | 9.64E-17 | -0.2638018 | 0.752 | 0.896 | 1.42E-13 |
| Rrad | 1.64E-15 | -0.355179 | 0.314 | 0.61 | 2.41E-12 |
| Nov | 1.73E-15 | -0.5100504 | 0.322 | 0.605 | 2.55E-12 |
| Has1 | 9.57E-15 | 0.54088597 | 0.68 | 0.801 | 1.41E-11 |
| Mgp | 4.78E-14 | -0.2579504 | 0.115 | 0.32 | 7.03E-11 |
| Nr4a3 | 1.24E-13 | 0.34288713 | 0.532 | 0.832 | 1.82E-10 |
| Osr2 | 1.59E-13 | 0.27506825 | 0.608 | 0.647 | 2.34E-10 |
| Cxcl13 | 4.80E-13 | 0.65563631 | 0.503 | 0.859 | 7.06E-10 |
| H2-Q7 | 9.84E-13 | -0.3060783 | 0.275 | 0.563 | 1.45E-09 |
| Thbs1 | 1.37E-12 | -0.3836758 | 0.343 | 0.602 | 2.01E-09 |
| Col4a1 | 1.82E-12 | 0.4358938 | 0.574 | 0.89 | 2.67E-09 |
| Igf1 | 1.20E-11 | 0.37885123 | 0.82 | 0.945 | 1.76E-08 |
| Igfbp7 | 1.29E-11 | -0.2610588 | 0.654 | 0.789 | 1.89E-08 |
| Cxcl9 | 6.80E-11 | -0.4283394 | 0.139 | 0.346 | 1.00E-07 |
| Mmp3 | 7.66E-11 | 0.63706183 | 0.538 | 0.844 | 1.13E-07 |
| Sbsn | 1.34E-08 | 0.91910207 | 0.666 | 0.867 | 1.98E-05 |
| Hmox1 | 1.05E-07 | 0.34226821 | 0.444 | 0.759 | 0.00015493 |
| Gas1 | 1.35E-07 | 0.3610028 | 0.635 | 0.723 | 0.0001989 |
| Ccl8 | 2.72E-07 | -0.2573871 | 0.406 | 0.625 | 0.0003997 |
| Stmn1 | 4.63E-07 | -0.3042611 | 0.3 | 0.554 | 0.00068143 |
| Atf3 | 4.90E-07 | 0.29613133 | 0.924 | 0.958 | 0.00072119 |
| Mfap4 | 4.98E-07 | -0.5693244 | 0.311 | 0.562 | 0.00073306 |

| | | | | | |
|---|---|---|---|---|---|
| Akr1c18 | 5.35E-07 | -0.3702399 | 0.822 | 0.896 | 0.00078685 |
| Iigp1 | 1.48E-06 | -0.4514139 | 0.41 | 0.605 | 0.00217468 |
| Thbs2 | 1.94E-06 | 0.36219054 | 0.559 | 0.674 | 0.00285018 |
| Lbp | 2.18E-05 | 0.33006848 | 0.693 | 0.866 | 0.03203745 |
| Emb | 5.95E-05 | -0.358784 | 0.183 | 0.48 | 0.08760305 |
| Pla1a | 7.75E-05 | 0.26806486 | 0.575 | 0.838 | 0.11415171 |
| Fst | 0.00013115 | 0.2949532 | 0.778 | 0.889 | 0.19305431 |
| Cxcl1 | 0.00016144 | 0.37177031 | 0.829 | 0.942 | 0.23763521 |
| C7 | 0.00019552 | 0.36965829 | 0.59 | 0.9 | 0.28781241 |
| Fabp5 | 0.0010871 | -0.5057688 | 0.404 | 0.573 | 1 |
| Wisp2 | 0.00153252 | 0.37498357 | 0.427 | 0.56 | 1 |
| Gadd45b | 0.00262158 | 0.29207636 | 0.684 | 0.867 | 1 |
| Hspa1b | 0.00375499 | 0.36208142 | 0.554 | 0.841 | 1 |
| Cryab | 0.01780611 | 0.35280283 | 0.801 | 0.914 | 1 |
| Hspa1a | 0.08331532 | 0.28726058 | 0.582 | 0.831 | 1 |
| AY036118 | 0.12207633 | 0.57563466 | 0.768 | 0.862 | 1 |
| Pcsk6 | 0.14183255 | 0.27905765 | 0.581 | 0.783 | 1 |
| Spon1 | 0.16810613 | 0.25044314 | 0.501 | 0.72 | 1 |
| Gja1 | 0.16863412 | 0.28911575 | 0.647 | 0.844 | 1 |
| Prkg2 | 0.41466798 | 0.43424206 | 0.564 | 0.835 | 1 |
| Cldn10 | 0.43240889 | 0.32431145 | 0.641 | 0.865 | 1 |
| Il6 | 0.59014673 | 0.35550727 | 0.517 | 0.688 | 1 |

**Table S5. P-values for two-tailed randomized testing identifying cell types significantly enriched by CellTagged populations.**

| Comparison (two-tailed) | α (following Bonferroni correction) | Cell type | p-value |
|---|---|---|---|
| Progenitor-CellTagged vs. Preadipocyte-CellTagged P32 precursors (Figure 2E) | 0.01 | Progenitor | < 0.0001 |
| | | Transitioning progenitor | 0.2932 |
| | | Immature preadipocyte | < 0.0001 |
| | | Committed preadipocyte | < 0.0001 |
| | | Multi ID | 0.1178 |
| Host APCs vs. *Sox9*-OE CellTagged progenitors (Figure 4E) | 0.01 | Progenitor | < 0.0001 |
| | | Transitioning progenitor | < 0.0001 |
| | | Immature preadipocyte | < 0.0001 |
| | | Committed preadipocyte | < 0.0001 |
| | | Multi ID | 1.0000 |
| Control-CellTagged preadipocytes vs. *Sox9*-OE preadipocytes (Figure 4H) | 0.01 | Progenitor | 0.0044 |
| | | Transitioning progenitor | < 0.0001 |
| | | Immature preadipocyte | < 0.0001 |
| | | Committed preadipocyte | 0.2693 |
| | | Multi ID | 1.0000 |

**Table S6. P-values for permutation testing identifying Louvain clusters significantly enriched for populations of CellTagged cells in homeostatic P32 dataset.**

| Cluster | CellTagged cells of progenitor-origin | CellTagged cells of preadipocyte- origin |
|---|---|---|
| 0 | 1.00000 | 1.00000 |
| 1 | 0.00000 | 0.00000 |
| 2 | 1.00000 | 1.00000 |
| 3 | 0.00015 | 0.12090 |
| 4 | 0.70565 | 0.00000 |
| 5 | 1.00000 | 0.00000 |
| 6 | 0.00000 | 1.00000 |
| 7 | 1.00000 | 0.17575 |
| 8 | 1.00000 | 1.00000 |
| 9 | 0.00000 | 0.45205 |
| 10 | 0.00005 | 0.02820 |
| 11 | 0.99985 | 1.00000 |
| 12 | 0.00000 | 0.00000 |
| 13 | 0.99620 | 0.99610 |
